# The genomic origins and evolutionary path to a key innovation in the world’s most venomous snakes

**DOI:** 10.64898/2026.06.23.733212

**Authors:** Jory van Thiel, Noah L. Dowell, Cara F. Smith, Elda E. Sanchez, Sean B. Carroll

**Author notes:** Corresponding author: Sean B. Carroll. **Email:**.

## Abstract

Evolutionary innovation is a key driver of the colonization of new environments and the adaptive radiations of major groups. Novel traits typically evolve through the modification of pre-existing characters but the genetic paths underlying their origin have been challenging to trace, and the general requirements for and relative order of different kinds of gene mutations have been difficult to assess. Here, we trace the genomic origins of four procoagulant venom toxins (factor X, factor V, group I phospholipase A_₂_, and Kunitz-type toxins) that collectively underlie a novel, especially potent blood-clotting venom type in the recently evolved Australian brown snake and taipan clade. We discover evidence for a previously unknown fifth toxin, coagulation factor VII, and show that the toxins evolved through two distinct genetic paths. The factor X and factor V toxins evolved through the sequential *de novo* co-option of ancestral clotting factor proteins that entailed their heterotopic expression in the venom gland, the fixation of segmental duplications containing each locus, and subsequent gain-of-function mutations that rendered factor X and factor V constitutively active. In contrast, the phospholipase A_₂_ and Kunitz-type toxins evolved by modifying the functions of neurotoxins that were part of the venom arsenal. Our findings support models in which innovative mutations in single-copy genes precede gene duplication in the evolution of novel proteins and offer a rare view into the genesis of a complex trait that has played a central role in a major adaptive radiation.

**Significance Statement:** This study investigates how an entirely new blood-clotting venom type evolved during the recent radiation of Australia’s iconic venomous snakes. We traced the key genetic events that occurred on the evolutionary path to one of the world’s most potent venoms. We found that the novel venom activity evolved through the sequential co-option of multiple proteins of the snake’s own blood-clotting system, followed by the modification of two venom neurotoxins into proteins with procoagulant activities. We suggest that these unique de novo gene co-options are seminal events that can unlock new ecological strategies, which in turn, may enable major adaptive radiations.

## Introduction

The evolution of key innovations such as the tetrapod limb, the wings of insects, birds, and bats, or C4 photosynthesis in plants has enabled new ways of living and catalyzed the emergence and radiation of major new clades. Hence, understanding the *evolutionary path* to novelty *-* the sequence of changes through which a new trait emerges over time, is a central quest of evolutionary biology. And one of the major objectives is to understand the *genetic paths* through which novel traits arise, which entails the identification of the genes involved, the kinds of changes that have occurred in these genes, and if possible, the order in which they have occurred. In addition, to gain a more general understanding of the origins of novelties, it is also important to consider why evolution has taken a particular genetic path, that is, why particular genes (and not others) are involved, why certain kinds of genetic changes have occurred (and not others), and why a particular order may have unfolded.

For example, with respect to the evolution of new morphological traits, a common genetic path has been identified (1, 2). The primary mechanism of morphological innovation is gene co-option, in which a gene or pre-existing genetic regulatory network (GRN) is deployed in a new developmental context (3). A variety of morphological novelties in plants and animals have been shown to arise via changes in the spatiotemporal expression of developmental regulatory genes (4–13) (reviewed in (14)). These gene expression changes have been shown or inferred to result largely from mutations in cis-regulatory elements (enhancers) that control gene transcription (6–9, 15, 16). By contrast, coding sequences changes that modify regulatory protein function are relatively rare, at least in animals, presumably because such mutations have deleterious pleiotropic effects. In addition, the fixation of new duplicates is uncommon among animal developmental regulatory genes perhaps due to their disruption of the balance of inputs in GRNs (1).

On the other hand, the fixation of new gene duplicates is widespread in the evolution of new proteins involved in a wide range of novel physiological traits (17–20). However, the path from ancestral gene to novel biochemical protein is often challenging to trace, with the requirements for and order of gene duplication events, coding sequence changes, and gene expression changes difficult to untangle. The increasing availability of high-quality genome sequences is making the evolution of many novel traits much more accessible and raises the prospect that a larger body of well-studied examples will reveal some generalities about the genetic paths to biochemical innovation.

Animal venoms are key innovations that fundamentally change how predators and prey interact, shifting them from physical combat to chemical warfare. Venom proteins enable animals to capture different types of prey and provide a new line of defense against predators. Because of these ecological advantages, venoms have evolved independently in numerous animal lineages, from various cnidarians to several major groups of arthropods, cephalopods, and vertebrates (21).

In snakes, several venom toxin families typify the two major clades of venomous snakes such as the three-finger toxins and group I phospholipase A_2_ toxins of elapids, and the metalloproteinase, serine protease, and group II phospholipase A_2_ toxins of vipers. These venom toxins have evolved from pre-existing proteins that function within the body and expanded through gene duplication and divergence (22–25). This ancestry allows for the potential tracing of the genetic events that transformed physiological proteins to biochemical weapons, however, all of these toxins evolved early within their respective lineages and from proteins whose ancestral functions are not well understood, making it is difficult to decipher the order in which events unfolded.

One striking example of a recent key innovation in venoms occurred when venomous snakes first colonized the Australian continent. Devoid of venomous snakes for most of its history (vipers never reached the continent), Australia’s elapids (Elapidae: Hydrophiinae), are inferred to have descended from an aquatic or semi-aquatic Asian ancestor (26, 27), which invaded Australasia as recently as 12 million years ago (mya) (28, 29) (Fig. 1). This colonization event spurred a continent-wide radiation of terrestrial elapids, and later, one of these lineages transitioned into marine habitats, triggering a second radiation of sea snakes (28) (Fig. 1).

**Fig. 1.**
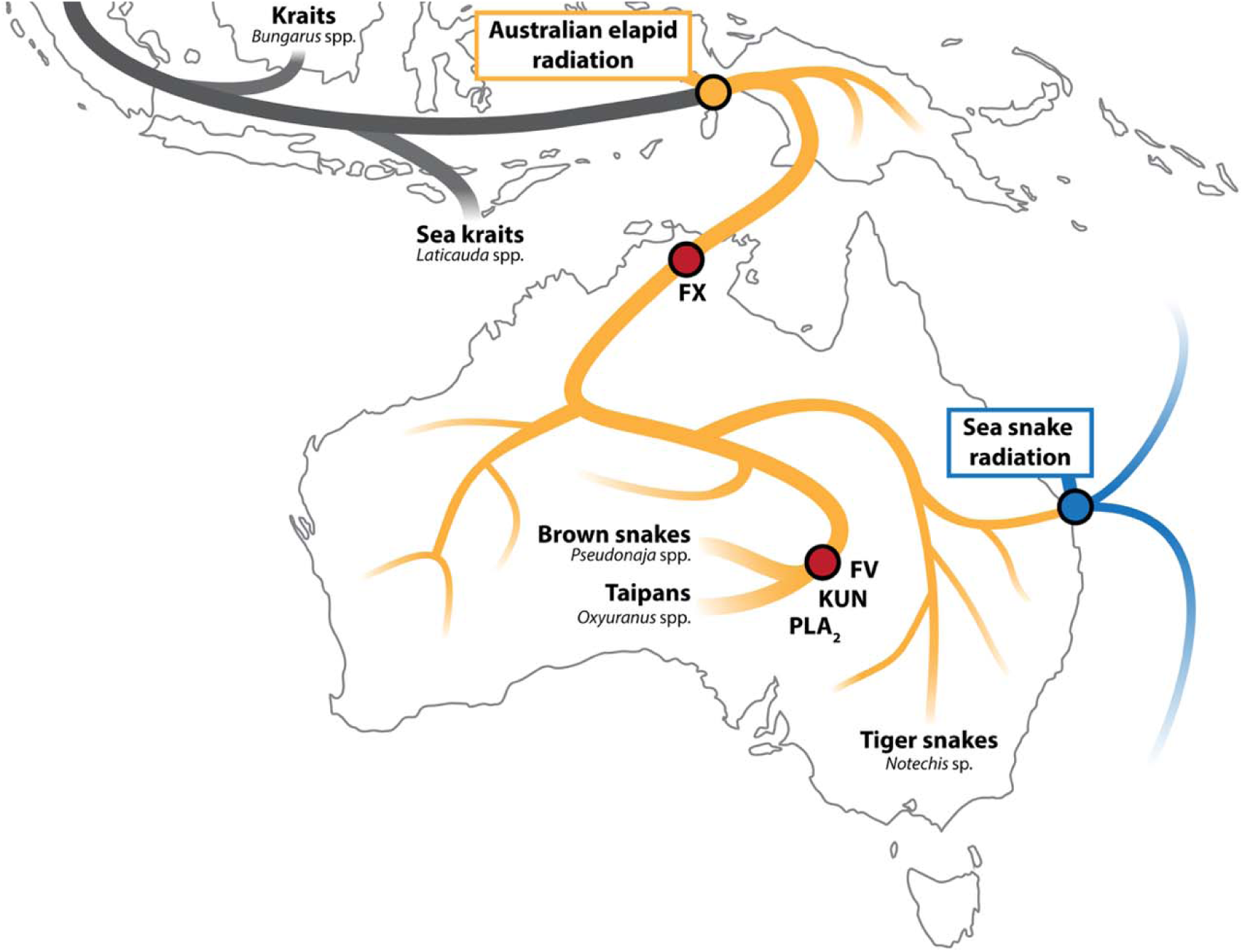
Evolutionary history of the Australian elapid radiation (Hydrophiinae) and the inferred origins of their procoagulant toxins. The Asian kraits (*Bungarus* spp.) and sea kraits (*Laticauda* spp.) are inferred to be sister taxa to all Australian elapids, with a common Asian ancestor of the sea kraits thought to have migrated to Melanesia and subsequently radiated into Australia (yellow circle). One of these terrestrial lineages moved into the marine environment, which spurred a second radiation of sea snakes (blue circle). Coagulation factor X (FX) was recruited early in the diversification of Australian terrestrial snakes (red circle). The monophyletic brown snake (*Pseudonaja* ssp.) – taipan (*Oxyuranus* spp.) clade evolved three additional procoagulant toxins: coagulation factor V (FV), group I phospholipase A_₂_ (PLA_2_), and Kunitz-type toxin (KUN; red circle).

While all elapid snakes produce venoms that cause paralysis, the terrestrial Australian elapid lineage evolved a unique additional activity: the induction of blood clotting. This novel procoagulant venom type is due to the presence of a homolog of the factor X clotting protein, a key enzyme that activates prothrombin in the vertebrate blood clotting pathway (30–35). The advent of venom factor X appears to have occurred early in their evolutionary history (36) and played a role in the success of the Australian elapid radiation because all extant clades produce this toxin (37) (Fig. 1).

As Australian elapids diverged across the continent, the monophyletic clade comprising brown snakes (*Pseudonaja* spp.) and taipans (*Oxyuranus* spp.) further expanded the arsenal of procoagulant toxins (Fig. 1). One of these new toxins is venom factor V, a homolog of blood coagulation factor V (38, 39), a non-enzymatic protein that forms a complex with coagulation factor X that enhances prothrombin-activation efficiency over 250,000-fold (40). In addition, two other classes of procoagulant toxins evolved in the brown snake-taipan clade including type I phospholipase A_2_ proteins (PLA_2_) (41) and Kunitz-type toxins, the latter of which selectively inhibit plasmin, the enzyme responsible for breaking down blood clots (42–44). By producing a novel set of toxins that both rapidly trigger blood clotting and also prevent the breakdown of these dangerous clots, brown snakes and taipans have evolved the most potent venoms among all land snakes (45).

While the individual procoagulant toxins have been well characterized biochemically, little is known about the nature or order of the genetic events underlying the evolution of procoagulant venom in Australian elapids. Here, we generated high-quality genome assemblies of the Eastern brown snake (*Pseudonaja textilis*), and inland taipan (*Oxyuranus microlepidotus*) and traced the genomic origins of the four elapid procoagulant toxins (venom factor X, venom factor V, group I PLA_2_, and Kunitz-type toxins) and discovered a putative fifth toxin, coagulation factor VII. We show that the different procoagulant toxins evolved through two distinct genetic paths. Venom factor X and venom factor V both arose via the *de novo* co-option of single copy liver-expressed clotting factor genes followed by their segmental gene duplication, whereas the procoagulant PLA_2_ and plasmin-inhibiting toxins evolved through the modification of members of protein families expressed in the venom gland (neofunctionalization). Our findings support models in which gene co-option precedes gene duplication in the evolution of novel proteins, and we suggest that the co-opted coagulation factors have played a similar role in the evolutionary success of Australian elapids as the major toxin types co-opted in other venomous snake families.

## Results

### Venom factor X evolved via gene co-option and a segmental duplication event

The first procoagulant toxin that was recruited into the venom arsenal of the Australian elapid radiation was venom factor X. The homology between the venom toxin and coagulation factor X has long been recognized (32–35), but the genomic origin of the toxin, and the genetic mechanism(s) through which it evolved have not been elucidated. To trace the origin of venom factor X, we first examined the genomic locus encoding coagulation factor X (*F10*) in two outgroup species, a colubrid (*Natrix helvetica*) and an Asian elapid (*Bungarus candidus*). We identified a single *F10* gene that is situated in a conserved syntenic region flanked on the 5′ end by genes encoding guanine nucleotide exchange factor DBS (*MCF2L*) and coagulation factor VII (*F7*), and on the 3′ end by Vitamin K-dependent protein Z (*PROZ*), PCI domain-containing protein 2 (*PCID2*), and Lysosome-associated membrane glycoprotein 1 (*LAMP1*) (Fig. 2A).

**Fig. 2.**
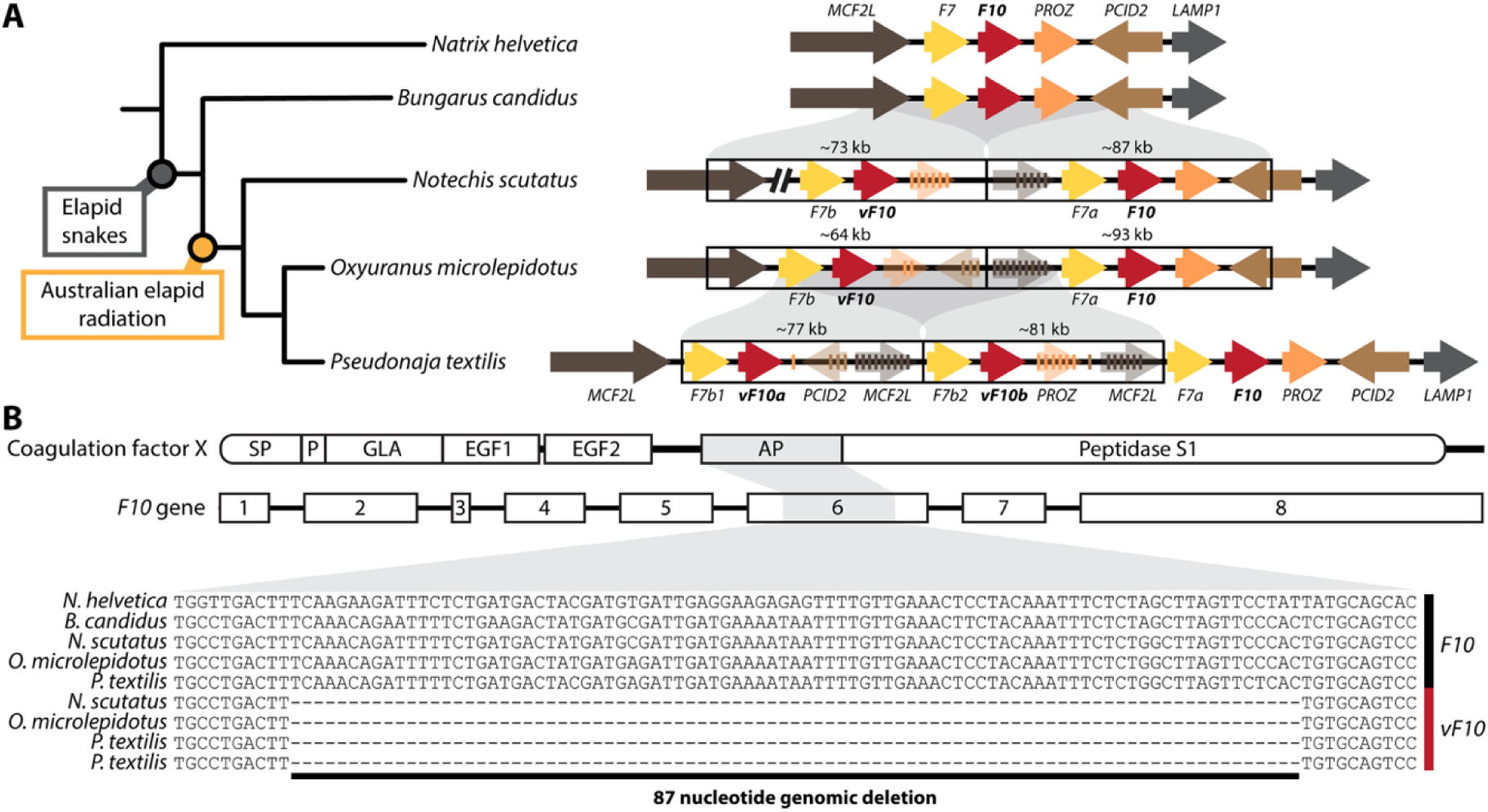
The venom factor X gene arose via a segmental duplication. (A) Schematic species phylogeny showing the genomic regions of the *F10* locus and its flanking genes in several representative snake species. The *F10* gene (red arrow) is located in a syntenic region flanked by *MCF2L* (dark brown), *F7* (yellow), *PROZ* (orange), *PCID2* (light brown), and *LAMP1* (grey). A single *F10* gene exists in non-Australian snakes (in *N*. *helvetica*, *B*. *candidus*), whereas Australian terrestrial elapids possess two *F10* genes (in *N*. *scutatus* and *O*. *microlepidotus*) or three *F10* genes (in *P*. *textilis*). The duplicated *F10* genes share the same flanking genes as the non-duplicated loci, which indicates they arose by segmental duplication events. The paralogs expressed in the venom gland are denoted *vF10*, those expressed primarily in the liver are denoted *F10.* The duplicated *F7* loci remain intact in both lineages, are expressed in the venom gland, and are denoted *F7a* and *F7b1* and *F7b2*. *PROZ* is found as a pseudogene only in the duplicated segments (indicated by transparent arrows). The break-line symbol represents a genomic region with low confidence. The species phylogeny is based on reference (49). (B) Schematic of the domain structure of coagulation factor X: signal peptide (SP), propeptide (P), GLA domain (GLA), EGF1 and EGF2 domains, activation peptide (AP), and a peptidase S1 domain. The *F10* gene comprises eight exons (sizes drawn to relative scale), with the activation peptide encoded by exon 6 (highlighted in grey). In *vF10* genes, exon 6 contains an 87-nucleotide deletion, whereas it remains intact in the ancestral *F10* genes.

We then identified the *F10* locus in several procoagulant species that express the F10 toxin in their venoms (*N*. *scutatus*, *O. microlepidotus* and *P. textilis*) (46). We found that these species differ in *F10* gene number, with two copies in *N*. *scutatus* and *O. microlepidotus* and three copies in *P. textilis*. These *F10* genes are not arranged as tandem duplicates but are instead flanked by the same set of neighboring genes found in the outgroup species *N. helvetica* and *B. candidus* (Fig. 2A). This arrangement reveals that two syntenic segments are present in the *O*. *microlepidotus* (∼64 and ∼93 kb) and *N*. *scutatus* (∼73 and ∼87 kb) genomes (Fig. 2A), and that there is an additional, third *F10*-containing segment in *P*. *textilis* (Fig. 2A). Based on these observations, we deduce that an intrachromosomal duplication including *F10* and all or part of three surrounding loci occurred in an ancestral Australian elapid lineage, followed by a second segmental duplication within the brown snake lineage (*SI Appendix*, Fig. S1).

To assess the evolutionary fate of genes within these duplicated segments, we examined the flanking genes in more detail. We found that the *F10* and *F7* paralogs remain intact in all species (Fig. 2A). In contrast, the *PROZ* locus has been pseudogenized by mutations that disrupt its coding sequence (transparent arrows) (Fig. 2A). Since *PROZ* is intact in the outgroup species (*N*. *helvetica* and *B*. *candidus*; Fig. 2A), the emergence of these pseudogenes after the segmental duplication events suggests that the duplicates of some genes were dispensable or perhaps deleterious while those that have been preserved may be beneficial.

To identify which of the *F10* paralogs encode the hemostatic factor X protein and which the venom toxin, we examined the transcripts encoded by each gene (*SI Appendix*, Fig. S2). We confirmed these assignments by analyses of long-read isoform sequencing (Iso-seq) of *O. microlepidotus* mRNA and short-read sequencing (RNA-seq) of *P*. *textilis* mRNA from liver and venom gland tissues. We found that one *F10* gene is primarily expressed in the liver and encodes the hemostatic protein (Table 1), and the second *F10* gene in *O. microlepidotus* (*vF10*) and the two genes in *P. textilis* (*vF10a* and *vF10b*) are expressed in the venom gland and encode venom toxins (Table 1).

**Table 1.**
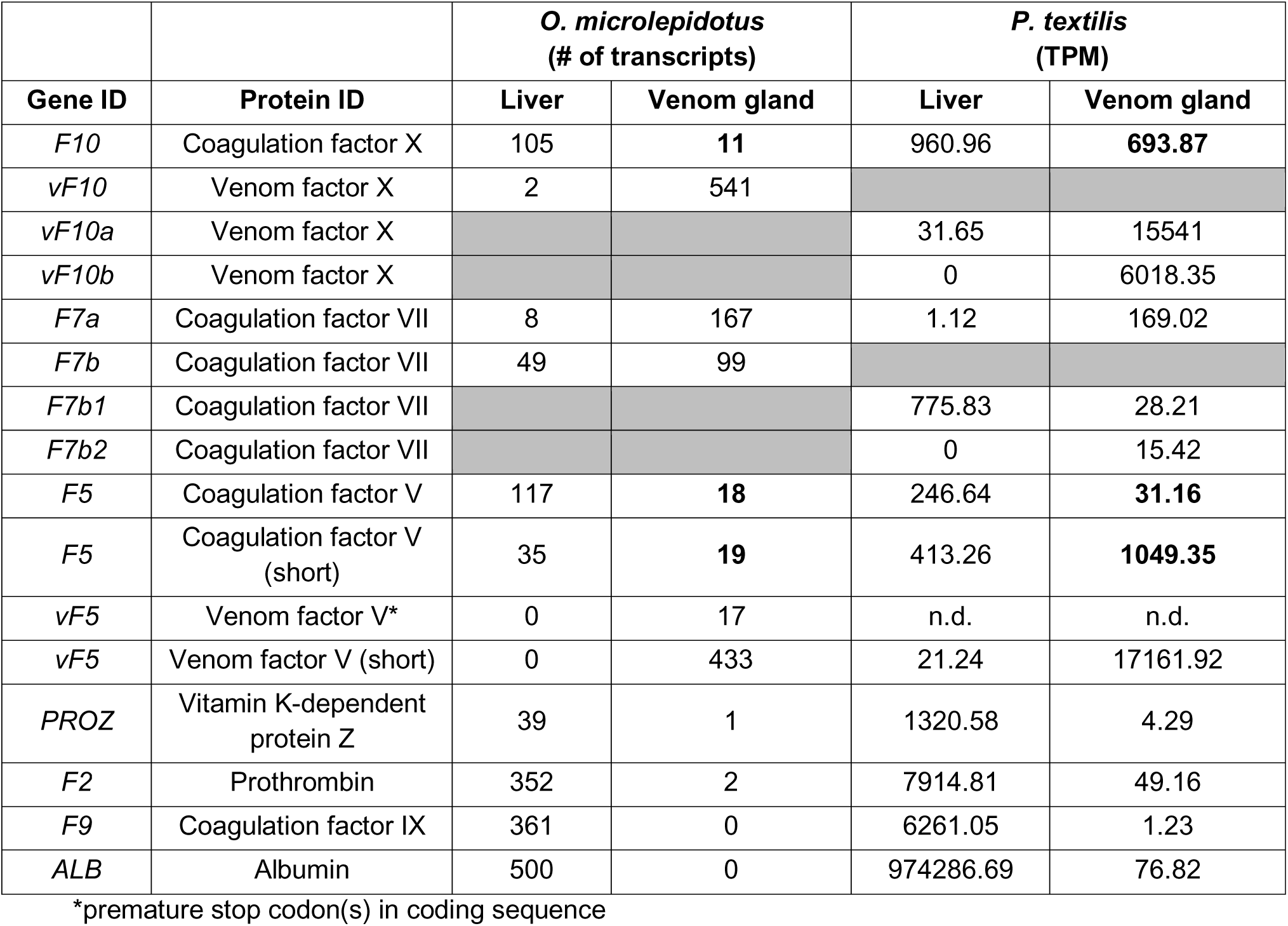
Coagulation factor gene expression in venom gland and liver tissues.

|  |  | *O. microlepidotus*<br>(# of transcripts) |  | *P. textilis*<br>(TPM) |  |
| --- | --- | --- | --- | --- | --- |
| Gene ID | Protein ID | Liver | Venom gland | Liver | Venom gland |
| *F10* | Coagulation factor X | 105 | **11** | 960.96 | **693.87** |
| *vF10* | Venom factor X | 2 | 541 |  |  |
| *vF10a* | Venom factor X |  |  | 31.65 | 15541 |
| *vF10b* | Venom factor X |  |  | 0 | 6018.35 |
| *F7a* | Coagulation factor VII | 8 | 167 | 1.12 | 169.02 |
| *F7b* | Coagulation factor VII | 49 | 99 |  |  |
| *F7b1* | Coagulation factor VII |  |  | 775.83 | 28.21 |
| *F7b2* | Coagulation factor VII |  |  | 0 | 15.42 |
| *F5* | Coagulation factor V | 117 | **18** | 246.64 | **31.16** |
| *F5* | Coagulation factor V<br>(short) | 35 | **19** | 413.26 | **1049.35** |
| *vF5* | Venom factor V* | 0 | 17 | n.d. | n.d. |
| *vF5* | Venom factor V (short) | 0 | 433 | 21.24 | 17161.92 |
| *PROZ* | Vitamin K-dependent<br>protein Z | 39 | 1 | 1320.58 | 4.29 |
| *F2* | Prothrombin | 352 | 2 | 7914.81 | 49.16 |
| *F9* | Coagulation factor IX | 361 | 0 | 6261.05 | 1.23 |
| *ALB* | Albumin | 500 | 0 | 974286.69 | 76.82 |

\*premature stop codon(s) in coding sequence

These results show that the *F10* gene was co-opted and duplicated in the evolution of the Australian elapid clade, but they do not reveal the order in which these events occurred. One important clue that co-option appears to have occurred first emerged from our detection of transcripts from the hemostatic *F10* gene in the venom glands of both *O. microlepidotus* and *P. textilis* (Table 1, bold). We confirmed that these signals were due to bona fide *F10* transcripts (and not *<u>vF10</u>* transcripts) by both Iso-seq transcripts of *O. microlepidotus* venom gland RNA (*SI Appendix*, Fig. S3) and short-reads of *P. textilis* venom gland RNA (*SI Appendix*, Fig. S4).The level of *F10* transcription in venom glands was not observed for several other liver-expressed genes including the *F2*, *F9*, and *PROZ* coagulation pathway genes and serum albumin (*ALB*) (Table 1). In addition, we also found evidence that the hemostatic factor X protein is present in venom from mass spectrometry analysis (*SI Appendix*, Fig. S5). These observations collectively indicate that *F10* expression in the venom gland is unlikely to simply reflect background noise. Rather, it is consistent with *F10* expression evolving in the venom gland when *F10* was a single-copy gene, before the duplication of the locus, and then persisting after gene duplication.

### The factor VII protein is a putative fifth procoagulant toxin

It is notable that the *F7* genes that were co-duplicated with *F10* remain intact while adjacent co-duplicated loci were pseudogenized. In normal hemostasis, coagulation factor VII activates coagulation factor X (47), so we considered the possibility that factor VII may have a role in venom that has not been previously reported. Indeed, we detected significant RNA expression of the co-duplicated *F7* paralogs (*F7a* and *F7b*) in venom gland tissues from both *P. textilis* and *O. microlepidotus* (Table 1), which we confirmed to encompass full-length transcripts by long-read Iso-seq analysis of *O. microlepidotus* venom gland RNA (*SI Appendix*, Fig. S6). We also sought to verify factor VII’s presence by mass spectroscopy analysis of multiple *P. textilis* venom samples and found several tryptic peptides unique to factor VII protein, suggesting that this coagulation factor is translated and secreted into venom (*SI Appendix*, Fig. S7). Taken together, the co-duplication of *F7* with *F10*, preservation of the *F7* gene duplicates, the transcription of *F7* in venom gland tissue, and the presence of factor VII protein in crude venom collectively suggest *that F7* was also co-opted early in the evolution of procoagulant venom. Understanding the potential functional role of factor VII in venom requires further biochemical investigation beyond the scope of this study.

### Neofunctionalization of the venom factor X toxin by partial genetic deletion of its activation peptide

The hemostatic form of factor X is secreted as an inactive protein (zymogen) that requires proteolytic removal of an activation peptide to become active, whereas the procoagulant factor X toxin is present as a constitutively active serine protease in venom (30–32). One notable difference between the venom-expressed toxin and the paralogous clotting factor, is that its activation peptide is truncated by 29 residues (27 versus 56 amino acids; *SI Appendix*, Fig. S2) (34, 35). This shortened activation peptide has been suggested to enable factor V–dependent activation of prothrombin (48), but it has been unclear whether this truncation results from genomic, post-transcriptional, or post-translational changes. Comparison of *F10* and *vF10* genomic loci from several species reveals an 87-nucleotide deletion in *vF10* exon 6, the region that encodes the activation peptide (Fig. 2B). This finding reveals that the constitutive activity of the venom protein is due to a deletion in the genome, and thus the venom protein has been neofunctionalized through a gain of function mutation.

### The *vF10* and *F7b* genes were lost early in the radiation of sea snakes

Sea snakes (part of the subfamily Hydrophiinae) evolved from a terrestrial Australian elapid ancestor and radiated into a clade with fully aquatic lifestyles (28, 49). This marine invasion appears to have had important consequences on venom composition, as previous work suggests that the venoms of multiple sea snake species (e.g., *Aipysurus laevis* and *Hydrophis* spp.) lack the ability to clot human plasma (36). Because sea snakes evolved from a clade that produces the procoagulant factor X toxin, this prompts the question of the fate of the *vF10* gene in sea snakes.

We compared the *F10* locus of the *O*. *microlepidotus* genome with those of five phylogenetically distinct sea snake species. We found a syntenic gene arrangement in all sea snake genomes in which a single *F10* gene was flanked on the 5′ end by the *MCF2L* and *F7* genes, and on the 3′ end by the *PROZ*, *PCID2*, and *LAMP1* genes (Fig. 3). The organization of this gene complex largely resembles that observed in the non-procoagulant outgroup *B*. *candidus* (Fig. 3). However, detailed inspection of the sea snake genomes revealed a unique genomic segment containing a *MCF2L* pseudogene spanning either exons 22–25 (in *A. laevis*) or exons 22-26 (in all *Hydrophis* spp.) (Fig. 3). The most parsimonious explanation for the presence of this *MCF2L* pseudogene is that it is a remnant of the original *F10/F7* segmental duplication in terrestrial Australian elapids (as represented by *O*. *microlepidotus*; Fig. 3). This specific genomic arrangement indicates that the major portion of the duplicated region, spanning *F7b*, *vF10*, *PROZ*, and pseudogene fragments of *PCID2* gene is no longer present, and has been deleted. The great similarity of this genomic region among multiple distinct sea snake species indicates that the deletion event(s) that led to the loss of the *vF10* gene occurred early in this marine radiation. Taken together, these observations suggest that the transition of a common ancestor of extant sea snakes to a marine environment, where they adapted to feeding on distinct prey, was accompanied by the genetic loss of venom factors X and VII.

**Fig. 3.**
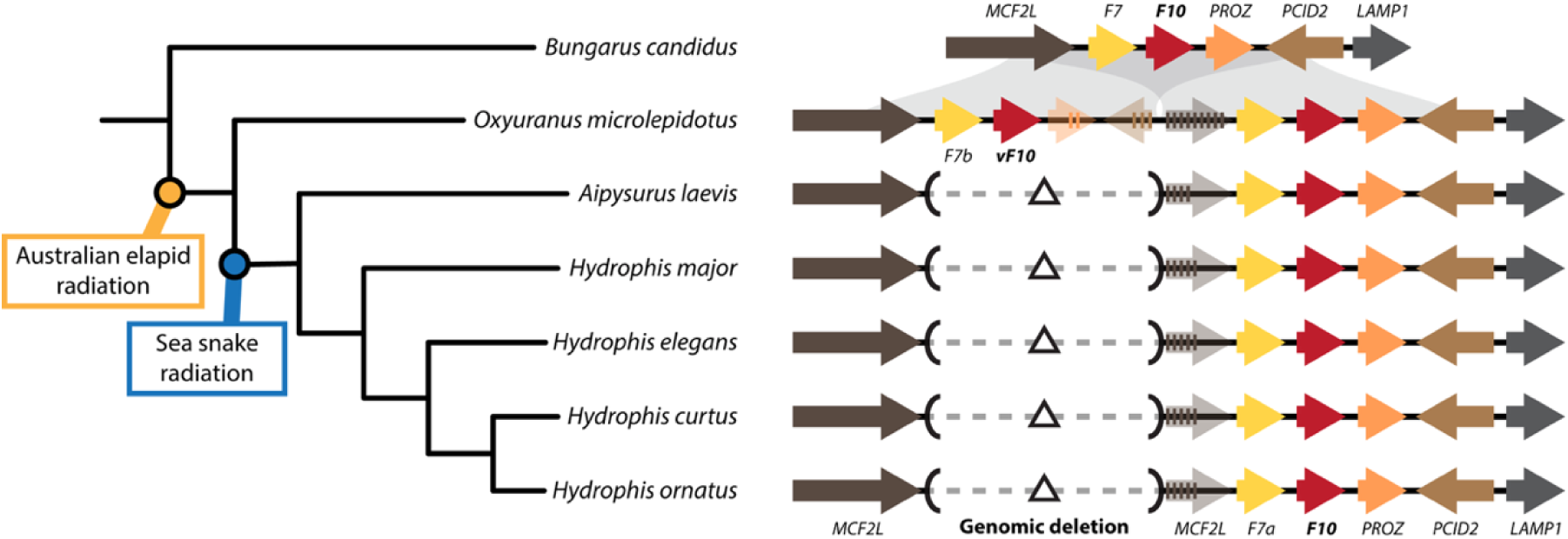
Genomic deletion of the *vF10* and *F7b* genes occurred early in sea snake radiation. Genomic comparison of the *F10* locus in an Asian elapid (*B*. *candidus*), terrestrial Australian elapid (*O*. *microlepidotus*) and several sea snake species (*Aipysurus laevis*, *Hydrophis major*, *H*. *elegans*, *H*. *curtus*, and *H*. *ornatus*). While the liver-expressed *F10* gene is present in all sea snake species, a large genomic segment containing the venom-expressed *vF10* and *F7b* genes is absent (as indicated by black triangles). Because sea snakes evolved from a clade of terrestrial Australian elapids that possess the *vF10* and *F7b* genes, we infer that both genes were lost during the ecological transition from land to sea. Pseudogene fragments are indicated by transparent arrows. The species phylogeny is based on reference (49).

### Venom factor V evolved via co-option and a segmental duplication event

One key protein found exclusively in the venoms of the Australian brown snake and taipan clade is the procoagulant toxin venom factor V. This component appears to have evolved from hepatic coagulation factor V (38, 39), but its genomic origin and the genetic mechanisms that shaped its evolution have not been resolved. We found that the evolutionary history of the venom factor V locus bears striking parallels to that of venom factor X because it also involved a segmental duplication event, pseudogenization of flanking genes, co-option of the clotting factor gene, and mutations that rendered it constitutively active.

We traced the origin of venom factor V by first examining several species that lack the venom toxin (*N. helvetica, B. candidus* and *N. scutatus*), in which we found a single coagulation factor V gene (*F5*) flanked on its 5′ end by genes encoding cell surface glycoprotein receptor 1 (*CD200R1*), coiled-coil domain-containing protein 80 (*CCDC80*), UDP-sugar transporter protein SLC35A5 (*SLC35A5*) and ubiquitin-like-conjugating enzyme ATG3 (*ATG3*), and its 3′ end by the thiamine transporter 1 (*SLC19A2*), the coiled-coil domain-containing protein 181 (*CCDC181*), and the golgin-45 (*BLZF1*) genes (Fig. 4A).

**Fig. 4.**
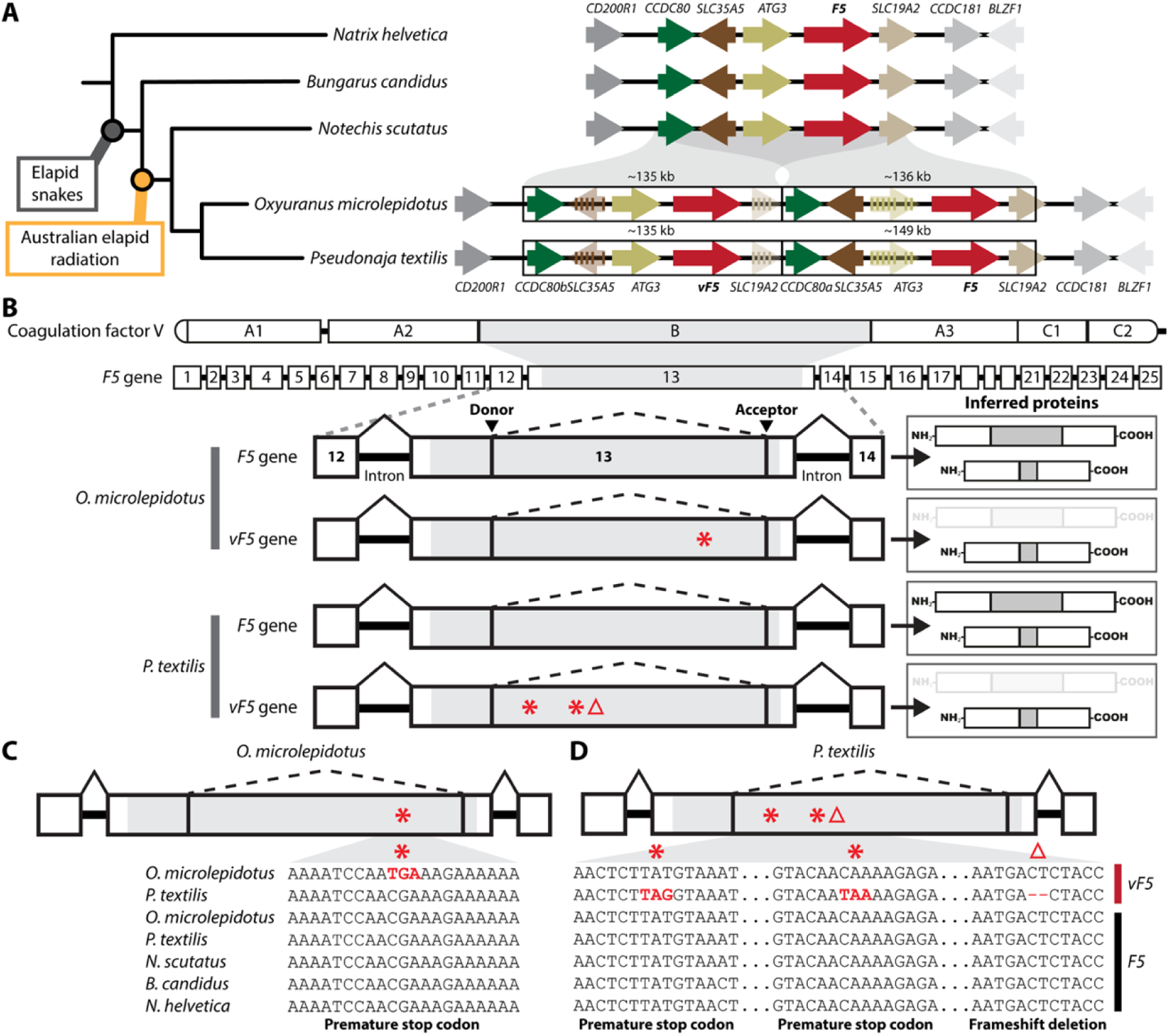
Venom factor V gene arose via a segmental duplication. (A) Schematic showing the genomic regions of the *F5* locus and its flanking genes in several representative snake species. The *F5* gene (red arrow) is located in a syntenic region flanked by *CD200R1* (dark grey), *CCDC80* (dark green), *SLC35A5* (dark brown), *ATG3* (olive green), *SLC19A2* (beige), *CCDC181* (grey), and *BLZF1* (light grey). A single *F5* gene exists in both non-Australian snakes (*N*. *helvetica* and *B*. *candidus*) and one Australian terrestrial snake (*N*. *scutatus*), whereas two *F5* genes are found in one Australian terrestrial clade (*O*. *microlepidotus* and *P*. *textilis*). The duplicated *F5* genes share the same flanking genes as the non-duplicated loci, which shows they arose through a segmental duplication. The paralog expressed in the venom gland is denoted *vF5*, the paralog expressed primarily in the liver is denoted *F5.* Within the duplicated segments one copy of *SLC35A5* and of *ATG3* are pseudogenes (indicated by transparent arrows). The species phylogeny is based on reference (49). (B) Schematic of the domain structure of coagulation factor V: signal peptide, A1, A2, B, A3, C1, and C2 domains. The *F5* gene comprises 25 exons (sizes drawn to relative scale), with the regulatory B-domain encoded by exon 13 (highlighted in grey). Within exon 13, the presence of donor and acceptor splice sites enables alternative splicing, which can remove most of the B-domain region (highlighted in grey). In the *F5* gene of *O*. *microlepidotus* and *P*. *textilis*, this event is predicted to produce two protein isoforms with B-domains of varying lengths. While these splice sites are also present in the duplicated *vF5* genes, exon 13 contains a premature stop codon in *O*. *microlepidotus* (indicated by a red asterisk) and two premature stop codons plus a frameshift deletion in *P*. *textilis* (indicated by a red asterisk and a red triangle). Because the nonsense mutations are located within the region that is removed by RNA splicing, these mutations do not affect translation of the shorter isoform but prevent translation of the canonical FV isoform in venom gland tissue. (C) and (D) Comparison of genomic regions containing nonsense mutations that are unique to the *vF5* genes and absent from the *F5* genes. Nonsense mutations are shown in red and indicated on the exon by a red asterisk for a premature stop codon or a red triangle for a frameshift deletion.

In contrast, we found two *F5* genes in the genomes of the two Australian species that express the factor V toxin (*O. microlepidotus* and *P. textilis*; Fig. 4A) (46). Each *F5* gene copy is located within a syntenic segment with similar flanking genes as observed in an Australian outgroup species (*N. scutatus*; Fig. 4A). These two syntenic segments are approximately 135 kb in length in *O*. *microlepidotus* (∼135 and ∼136 kb) and *P*. *textilis* (∼135 and ∼149 kb) and they share similar genomic boundaries, with the 5′ end located near exon 1 of *CCDC80* and the 3′ end positioned between exons 4 and 5 of *SLC19A2* (Fig. 4A). Therefore, we infer that a large genomic segment containing the *F5* gene was duplicated and retained in the common ancestor of the monophyletic taipan–brown snake clade.

While the duplicated genes *F5* and *CCDC80* are intact, one paralog of *SLC35A5* and of *ATG3* contain deletions in some exons that cause frameshifts and thus appear to have become pseudogenes in both taipan and brown snake genomes (Fig. 4A). Since these genes are intact, single-copy loci in species lacking the duplicated segment (*N. helvetica, B. candidus* and *N. scutatus*), we infer that, after the segmental duplication, some duplicates may have been functionally advantageous and were retained while others were dispensable or deleterious and were inactivated by mutations.

The presence of two *F5* gene copies raises the question of in which tissue (liver and/or venom gland) each paralog is transcriptionally active. Because previous work suggested that the *F5* gene may express multiple isoforms (50), we analyzed transcripts by long-read Iso-seq of *O*. *microlepidotus* liver and venom gland RNA. In both tissues, we detected RNA transcript isoforms encoding two protein isoforms (1919 versus 1459 amino acids) that share the same domain structure (A1–A2–B–A3–C1–C2) but differ in the length of their B-domain (Table 1; Fig. 4B). We identified one paralog as the liver-expressed *F5* gene because it encodes the two RNA isoforms found in liver tissue, with the long B-domain isoform encoding the canonical coagulation factor V protein, while the shorter B-domain isoform, hereafter termed FV-short, is of unknown function (*SI Appendix*, Fig. S8). The second paralog was identified as the venom-expressed *F5* gene (*vF5*) because we detected two RNA isoforms in venom gland tissue that aligned to this gene, with the shorter RNA isoform encoding FV-short, the venom toxin (Table 1; *SI Appendix*, Fig. S8). The orthologous *P*. *textilis F5* and *vF5* genes show the same pattern of tissue expression (Table 1; *SI Appendix*, Fig. S8).

These results show that the *F5* gene was co-opted and duplicated in the most recent common ancestor of the taipan/brown snake clade, but they do not reveal the order in which these events occurred. Importantly, similar to the *F10* gene, we also found significant transcription of the hemostatic *F5* gene in venom gland tissue of both *O. microlepidotus and P. textilis* (Table 1, bold), which was confirmed by full-length transcripts in *O. microlepidotus* venom gland RNA (*SI Appendix*, Fig. S9) as well as by short-read mapping of *P. textilis* venom gland RNA (*SI Appendix*, Fig. S10). We also detected the corresponding factor V protein in venom (*SI Appendix*, Fig. S11). These observations are consistent with *F5* expression first evolving in the venom gland when it was a single-copy gene, before the duplication of the locus, and persisting after gene duplication.

### Neofunctionalization of the venom factor V protein through loss of a regulatory domain

Activation of the hemostatic form of factor V requires proteolytic removal of its long B-domain, whereas the factor V venom toxin has a very short B-domain that has been suggested to underlie its constitutive activity (51). The mechanism underlying the truncation of venom factor V’s B-domain region could involve genomic deletion, alternative splicing, or a post-translational event. To investigate these possibilities, we first sought to understand how the two different-sized B-domains are encoded by both *F5* paralogs. Since the entire B-domain is encoded by exon 13, we analyzed how the two RNA isoforms map to this exonic region. We found that the longer RNA isoform spans the entire exon 13, but the shorter variant includes only two short coding sequences at the 5′ and 3′ ends of the exon, flanked by a donor and acceptor splice site (*SI Appendix*, Fig. S12). This observation suggests that the shorter RNA isoform encoding the venom toxin is generated through alternative splicing within exon 13. We infer then that the venom form became constitutively active due to a splice-based excision of the region encoding the regulatory B-domain.

This inference in turn raises the question of the fate of the long protein isoform in venom gland tissue. Close inspection of exon 13 of the *vF5* reveals a premature stop codon in *O. microlepidotus*, and two premature stop codons as well as a frameshift deletion in *P*. *textilis*; none of which are shared between the two species (Fig. 4C). In contrast, this exon remains intact in the liver-expressed *F5* paralog (Fig. 4C). These mutations are predicted to result in the premature termination of the translation process, producing truncated proteins. The genomic positions of these inactivating mutations, however, are all within the region encoding the long protein isoform, none are in the region encoding the short venom form because they are removed during the splicing of the shorter venom transcript (Fig. 4C). Therefore, we conclude that the *vF5* gene lost the ability to produce the canonical coagulation factor in the venom gland due to distinct mutations that appear to have occurred independently in taipan and brown snakes.

### A novel procoagulant phospholipase A_2_ arose via neofunctionalization of a venom neurotoxin gene

Elapid venoms typically contain PLA_₂_-type toxins that are derived from an ancestral group I PLA_2_ protein, some of which inhibit blood clotting and some of which act as neurotoxins. Australian brown snakes, however, have also evolved PLA_2_ toxin(s) that promote coagulation (41). To investigate the origin(s) of these novel procoagulant toxins, we first sought to identify the number, location, and arrangement of their genes in the brown snake genome.

In a cluster of seven PLA_2_ genes in *P*. *textilis*, we identified two venom-expressed genes that encode known procoagulant PLA_2_ proteins (41) (red arrows; Fig. 5A; *SI Appendix*, table. S1). We also detected one candidate procoagulant toxin in a complex of ten PLA_2_ genes in *O*. *microlepidotus* based on shared genomic synteny and sequence similarity (although the biochemical activity of this taipan protein has yet to be functionally validated) (red arrow; Fig. 5A; *SI Appendix*, Fig. S13). The *P*. *textilis* and *O*. *microlepidotus* PLA_2_ gene complexes also contain the non-venom expressed phospholipase A_2_ group IB gene (*PLA2G1B*; orange arrows) as well as four (*P. textilis*) or eight (*O. microlepidotus*) venom-expressed PLA_2_ genes that encode either monomeric or multimeric neurotoxins (52, 53) (light and dark blue arrows; Fig. 5A; *SI Appendix*, Tables S1 and S2). The arrangement of the procoagulant PLA_2_ genes adjacent to these other PLA_2_ genes suggests that they arose from some gene(s) in the complex, although the identity of that gene is not initially apparent due to the number and diversity of PLA_2_ genes in each complex.

**Fig. 5.**
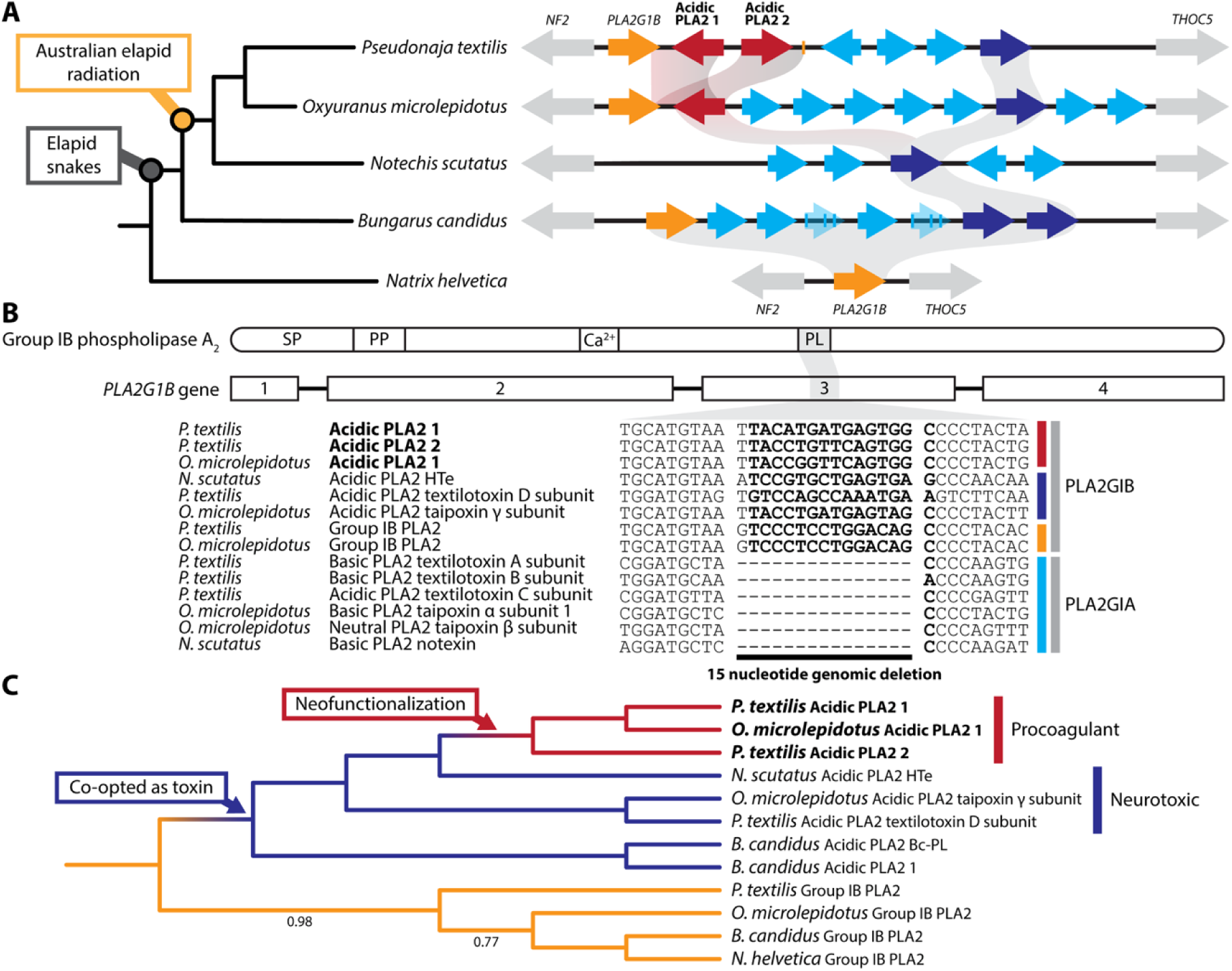
The procoagulant phospholipase A_2_ evolved through modification of a neurotoxin gene. (A) Schematic showing the genomic regions of the group I phospholipase A_₂_ (PLA_₂_) locus and its flanking genes in several representative snake species. Venom-expressed genes encode diverse PLA_₂_ toxins, including procoagulant PLA2GIB proteins (red arrows), PLA2GIB proteins with neurotoxic or unknown activity (dark blue), and neurotoxic PLA2GIA proteins (light blue). Non-venom expressed genes encode the related *PLA2G1B* gene (orange) and two unrelated *NF2* and *THOC5* flanking genes (grey). The *O*. *microlepidotus* Acidic PLA2 1 is a putative procoagulant protein based on its orthologous relationship with *P*. *textilis* Acidic PLA2 1, but the protein has not been biochemically characterized. Group I PLA_2_ toxins evolved from an ancestral single-copy *PLA2G1B* gene and subsequently expanded into numerous neurotoxins in elapid snakes. From one of these PLA2GIB neurotoxins, the procoagulant variants later evolved in brown snakes (and presumably taipans). Pseudogene fragments are indicated by transparent arrows. The species phylogeny was based on reference (49). Full nomenclature for all genes shown is in *SI Appendix*, Fig S15. (B) Schematic of the domain structure of group IB PLA_2_: signal peptide (SP), propeptide (PP), calcium binding domain (Ca^2+^), and the pancreatic loop (PL). The *PLA2G1B* gene comprises 4 exons (sizes drawn to relative scale), with the pancreatic loop encoded by exon 3 (highlighted in grey). In the genes encoding PLA2GIA toxins (light blue), exon 3 contains a 15-nucleotide deletion, whereas it remains intact in the ancestral PLA2GIB genes (orange, dark blue and red). (C) Protein phylogeny of PLA2GIB sequences. The procoagulant PLA_2_s cluster with the neurotoxic PLA_2_s suggesting that the procoagulant proteins evolved from a neurotoxic protein (neofunctionalization). All of these venom-expressed PLA2G1B sequences cluster with non-venom expressed group IB PLA_2_s, indicates that venom-expressed group I PLA_2_ genes evolved through the co-option of an ancestral, single-copy *PLA2G1B* gene. Only Bayesian posterior probabilities below 1.00 are shown.

To help resolve the ancestry of the procoagulant PLA_2_ genes we used both comparative genomic and phylogenetic analyses of group I PLA_2_ genes from taxa at various evolutionary distances from the brown snake-taipan clade. In the Australian elapid *N*. *scutatus* that lacks procoagulant PLA_2_s, we identified five PLA_2_ genes, several of which encode potent neurotoxins (54, 55) (blue arrows), whereas the *PLA2G1B* gene is absent from this species (Fig. 5A). In the Asian elapid *B*. *candidus*, we identified the *PLA2G1B* gene as well as five venom-expressed PLA_2_ genes, three of which encode subunits of a heterodimeric neurotoxin with a Kunitz-type toxin (56) (light blue arrows; Fig. 5A). Finally, in a non-venomous colubrid (*N*. *helvetica*), an outgroup to the elapids, we identified a single *PLA2G1B* gene (Fig. 5A). The arrangement and distribution of PLA_2_ genes suggests that the PLA_2_ gene complexes expanded from the *PLA2GIB* gene and we infer that the venom-expressed genes arose by co-option of *PLA2G1B* in an early elapid ancestor.

A second and key clue to the evolutionary history of these genes emerged from alignment of the PLA_2_ genes which revealed that they fall into two distinct size classes based on variation in the length of exon 3 due to the presence/absence of a 15 bp sequence that encodes a five amino acid “pancreatic loop” in PLA_2_ group IB proteins (*SI Appendix*, Fig. S13). We found that this sequence is present in the procoagulant toxin genes (red arrows) as well as *PLA2G1B* (orange arrows), and toxin paralogs with either neurotoxic or unknown activity (dark blue arrows), but absent from numerous other neurotoxins (light blue arrows; Fig. 5B). Based on the presence of the sequence in the ancestral *PLA2G1B* gene, we infer that the lack of the sequence in these neurotoxin genes is due to a deletion event.

The presence of this sequence feature in the procoagulant toxin proteins led us to hypothesize two possible scenarios for their origin: either a second co-option of the adjacent, non-venom expressed *PLA2G1B* gene or the neofunctionalization of a pre-existing venom neurotoxin that retained the pancreatic loop. To distinguish between these hypotheses, we constructed a protein phylogeny of hypothetical translations of all group IB genes. At the base of our phylogeny, we observed a clade comprised of *PLA2G1B* genes with a sister cluster of all venom-expressed genes (Fig. 5C). Situated within this venom toxin clade is a well-supported cluster of procoagulant toxin genes that group with the neurotoxin genes and not with the ancestral *PLA2G1B* genes (Bayesian posterior probability = 1.00; Fig. 5C), which would have been expected had there been a second co-option of *PLA2G1B*. Therefore, we conclude that the procoagulant toxins are derived from and evolved through neofunctionalization of a pre-existing venom neurotoxin gene.

### Neofunctionalization of a pre-existing venom Kunitz-type toxin resulted in a potent plasmin inhibitor

Brown snake and taipan venoms not only contain toxins that trigger uncontrolled blood clotting, but some species also have toxins that slow the breakdown of these harmful clots. This function is carried out by Kunitz-type toxins that inhibit plasmin, the serine protease responsible for dissolving fibrin clots (42–44). To investigate the genomic origin(s) of these plasmin-inhibiting toxins, we examined the genomic organization of the Kunitz-type inhibitor locus in the brown snake and taipan genomes. We uncovered an evolutionary history similar to that of the procoagulant PLA_2_ proteins in that the Kunitz-type plasmin inhibitors evolved from a venom-expressed Kunitz-type toxin.

The *P*. *textilis* genome harbors four venom-expressed genes, two of which encode known plasmin inhibitors (textilinin-2; Fig. 6A, red arrows), while the other two genes encode known non-plasmin inhibitors (textilinin-3 and −7; Fig. 6A, blue arrows; *SI Appendix*, Fig S14 and Table S1) (43). In comparison, the *O*. *microlepidotus* genome contains two venom-expressed genes that encode proteins related to non-plasmin inhibitors (microlepidin-7 and −5; Fig. 6A, blue arrows; *SI Appendix*, Fig S14 and Table S2). Next to those genes, we observed a pseudogene related to the textilinin-7 gene, which appears to have been lost through a genomic deletion of exon 2 (transparent blue arrow; Fig. 6A). The gene encoding a previously characterized plasmin inhibitor in the coastal taipan (*Oxyuranus scutellatus*) venom (44) was not detected, suggesting that this locus is not present in our inland taipan genome assembly.

**Fig. 6.**
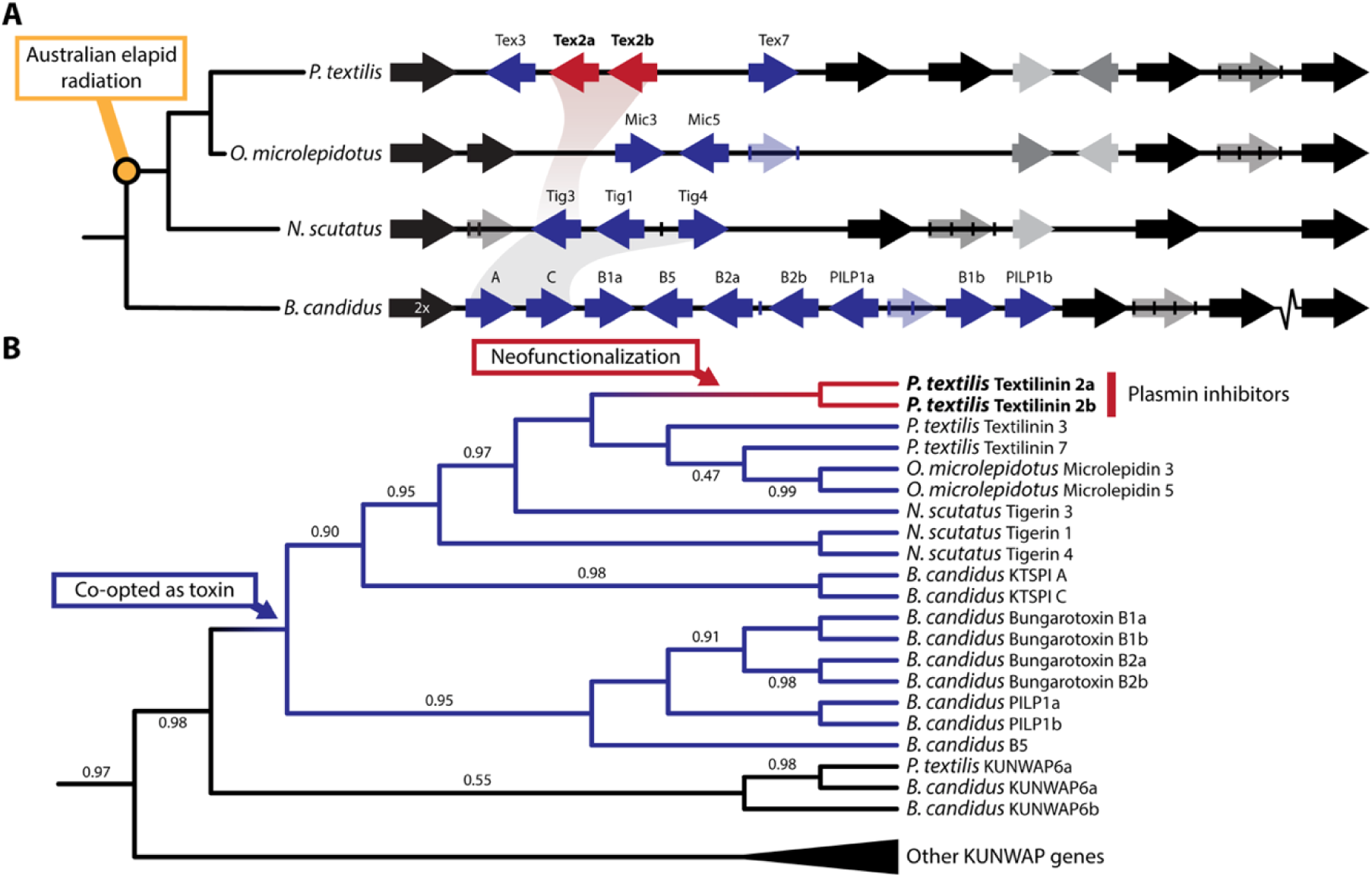
Neofunctionalization of venom-expressed Kunitz-type toxins into plasmin-inhibiting proteins. (A) Schematic showing the gene arrangement of venom-expressed and non-venom-expressed Kunitz-type serine protease inhibitor genes among different elapid snakes. The venom-expressed genes are either encoding plasmin-inhibiting toxins (red arrows) or putative neurotoxins (dark blue), or, waprins (dark grey). The non-venom-expressed encode either Kunitz proteins (light grey) or Kunitz–Waprin proteins (black). Pseudogenes are indicated by transparent arrows. The break-line symbol represents a genomic region with low confidence. The species phylogeny is based on reference (49). (B) Protein phylogeny of Kunitz-type inhibitors reveals that the plasmin-inhibiting proteins (red branches) cluster within a clade of venom-expressed Kunitz-type toxins (blue branches), suggesting that the plasmin-inhibiting function evolved via neofunctionalization. This monophyletic clade of Kunitz-type toxins clusters with a Kunitz–Waprin protein (KUNWAP6; black branches), suggesting that this ortholog represents the ancestral protein co-opted as a venom toxin. For the full phylogeny that includes all Kunitz–Waprin protein sequences see *SI Appendix*, Fig. S16). The phylogeny was rooted using the single-copy KUNWAP1 ortholog from *C*. *bottae* as the outgroup. Only Bayesian posterior probabilities less than 1.00 are shown.

Adjacent to the venom-expressed Kunitz genes, we identified seven non-venom-expressed genes in *P*. *textilis* and six in *O*. *microlepidotus* (Fig. 6A). Within this gene cluster, most genes encode both a Kunitz and WAP domain (black arrows), whereas a few encode either only a Kunitz domain (light grey arrows) or only a WAP domain (dark grey arrows; Fig. 6A).

To trace the ancestral gene(s) from which the two plasmin-inhibiting toxins evolved, we examined this multigene complex in both closely and more distantly related species. In the two venomous elapids *N*. *scutatus* and *B*. *candidus*, we identified large gene clusters composed of mainly non-venom-expressed genes (black and grey arrows), as well as several Kunitz-type toxin genes (blue arrows), including those that encode neurotoxins (57) (Fig. 6A).

The two genes encoding plasmin inhibitors in the *P*. *textilis* genome are flanked by both venom-expressed genes encoding non-plasmin-inhibitory proteins (43), as well as by non-venom-expressed genes encoding physiological proteins. These observations suggest two possible evolutionary scenarios: the plasmin inhibitors either evolved through co-option of a non–venom-expressed gene or through neofunctionalization of a pre-existing venom-expressed protein. To distinguish between these alternatives, we constructed a phylogenetic tree using protein sequences of our annotated Kunitz-type toxin genes from the different elapid species. This phylogeny formed multiple monophyletic clades, with varying degrees of statistical support, of which most genes encode Kunitz–Waprin proteins (Fig. 6B). Within one of these clades, we observed a strongly supported clade comprising all elapid Kunitz-type toxin genes (Bayesian posterior probability = 1.00; Fig. 6B). In this venom-expressed clade, the two plasmin inhibitors cluster with tigerin 3 (Bayesian posterior probability = 1.00; Fig. 6B). Because Tigerin 3 (Tig3) appears to be the closest relative of the plasmin-inhibiting toxins, we infer that these novel plasmin-inhibiting paralogs evolved through neofunctionalization of a pre-existing venom toxin, and not through the *de novo* co-option of a non-venom expressed gene.

## Discussion

In this study, we traced the evolutionary origin of four procoagulant toxins that collectively encode a key biochemical innovation in a group of Australian elapids, and discovered evidence for a previously unknown, putative fifth procoagulant toxin (coagulation factor VII). We found that these toxins evolved through two distinct genetic paths. The venom forms of coagulation factor X and coagulation factor V (and coagulation factor VII) evolved via the *de novo* co-option of physiological proteins via their heterotopic expression in the venom gland, the fixation of segmental duplications containing each locus, and subsequent gain-of-function mutations that rendered the factor X and factor V proteins constitutively active. By contrast, procoagulant PLA_2_s and plasmin-inhibiting Kunitz-type toxins did not evolve by *de novo* co-option of physiological proteins, but by the neofunctionalization of pre-existing venom toxins that had different biological and biochemical activities.

These observations raise questions concerning the order of genetic events along each path, and how this specific example might inform our general understanding of the genetic origins of novel proteins. They also raise the question of why Australian elapids took these specific genetic paths, which we explore by comparing them with the distinct path taken by some vipers that convergently evolved a procoagulant venom.

### The genetic path of innovation

Our inference that the procoagulant toxins evolved through two distinct genetic paths is based on our observations that different sets of genetic events occurred in the evolution of venom factors V and X (and VII) than in the evolution of the PLA_2_ and Kunitz-type toxins. To understand the origins of novelties, it is of great interest to attempt to determine the order of genetic events, and whether certain steps must occur or are more likely to have occurred before others. One central issue concerns the role and relative order of gene duplication.

In his landmark book, Ohno proposed that gene duplication was a necessary prerequisite for the origin of new protein functions because duplication would allow one copy to retain the ancestral function while the second copy could incorporate new functional mutations (neofunctionalization) (17). However, this classic model has been challenged on the theoretical grounds that if newly duplicated genes are selectively neutral, they must acquire new, innovative, and selectable mutations before acquiring inactivating mutations (nonsense mutations, frameshifts and deletions) (58–60). It is now recognized that the latter is much more probable than the former. Therefore, it has been proposed that innovative mutations may occur initially in single-copy genes and impart some new function such that subsequent gene duplicates may then be positively selected for and fixed on the basis of the new function (60, 61). A variety of empirical studies have demonstrated innovative mutations occurring before gene duplication (62–65). Several observations here prompt us to postulate the relative order of major genetic events in the evolution of procoagulant venom toxins that bear on these general models of genetic innovation.

### The de novo co-option path

Once elapids reached the shores of Australia, a continent-wide radiation venomous snakes unfolded. One striking innovation that emerged early during their colonization was the recruitment of coagulation factor X into their venom. This novel venom toxin enabled these snakes to target a different physiological pathway in the new prey they encountered as they spread across Australia. The role of factor X in the radiation of the Australian elapids raises the central question of what initial mutation(s) occurred that enabled the *F10* gene to become part of the venom arsenal?

We suggest that the first genetic change in the ancestral liver-expressed, clotting factor gene was the gain of its heterotopic expression in venom gland tissue (co-option; Fig. 7, step i) (our line of reasoning also applies to venom factor V, discussed below). Two key pieces of evidence for this scenario are the low but significant RNA expression of the predominantly liver-expressed *F10* orthologs in venom gland tissues and the presence of the corresponding hemostatic proteins in venom. If expression in the venom gland evolved after gene duplication, then one of two *F10* genes (the liver-expressed gene) would not be expected to be transcriptionally active in venom gland tissue. The observed expression of the liver-expressed *F10* paralog in the venom gland is consistent with the ancestral single-copy *F10* gene evolving expression in the venom gland before gene duplication, then decaying to some degree in the liver-expressed paralog after duplication.

**Fig. 7.**
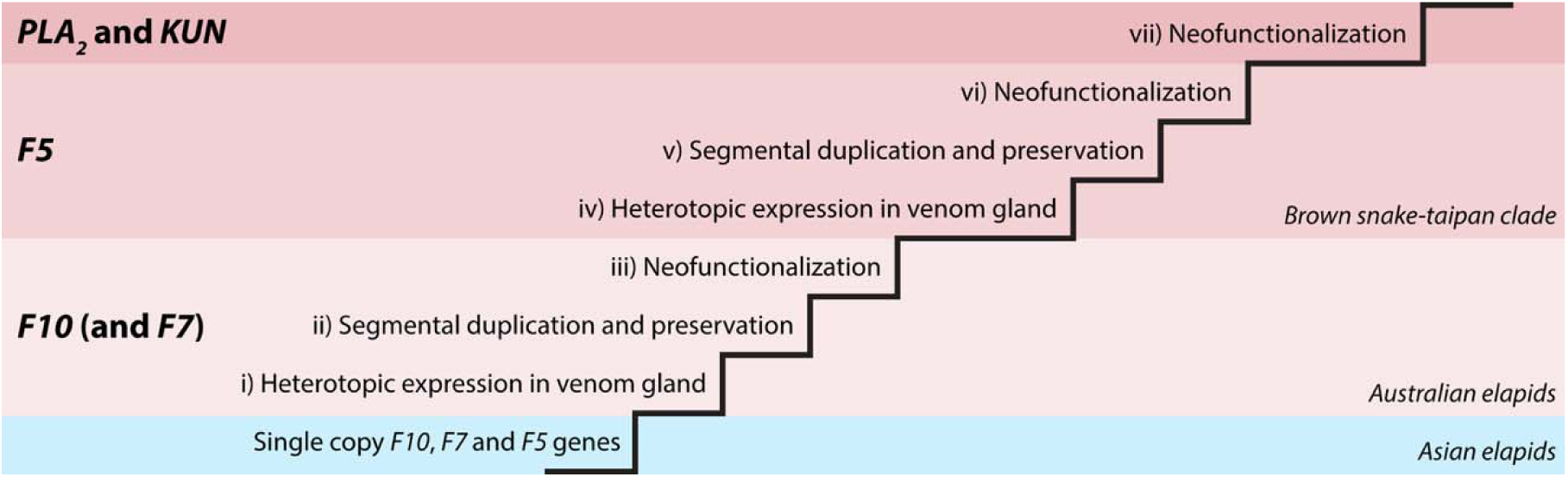
Proposed evolutionary and genetic paths to a potent procoagulant venom in the Australian brown snake and taipan clade. The evolutionary path from a neurotoxic ancestral Asian elapid venom to a highly potent procoagulant venom in the Australian brown snake-taipan clade involved a sequence of changes in toxin composition beginning with the incorporation of Factor X (and FVII) in the most recent common ancestor of Australian elapids, followed by the incorporation of factor V, procoagulant PLA_2_, and plasmin-inhibiting Kunitz proteins in the brown snake-taipan clade. The inferred genetic paths by which brown snakes and taipan evolved their procoagulant venom are shown schematically as a staircase, where each step represents a key genetic event (see text).

A third piece of evidence supporting *F10* expression in the venom gland as the initial genetic change concerns the fixation of the segmental duplication containing the *F10* locus as well as *PROZ* and *F7*. The duplication of large chromosomal regions alters the copy number of genes within the duplication and may be deleterious if copy number variation at any locus negatively affects fitness, which is often the case (66, 67). Furthermore, large duplications are expected to be unstable due to increased rates of non-allelic recombination that generate deletions and duplications (68). Therefore, segmental duplications are unlikely to become fixed in the genome under neutral or deleterious conditions. The preservation of the segmental duplications in Australian elapids suggests that some gene(s) within each duplicated region conferred a selective advantage sufficient to drive fixation of the entire genomic segment (Fig. 7, step ii).

If, as we have suggested above, the single-copy coagulation factor genes were already expressed in the venom gland, then the immediate functional consequence of their segmental duplication would be to increase toxin gene expression (via increased gene dosage). Selection for increased expression of *F10* (and possibly *F7*) in venom gland tissue may then have driven the fixation of the large chromosomal duplicates; an evolutionary force previously suggested for the retention of duplicated venom genes (23, 69). The co-duplication of two venom-expressed genes (*F10* and *F7*), both of which are known to activate one another (47), may have further increased the probability of their fixation relative to either gene alone.

We note that the duplication of the *PROZ* gene probably did not contribute to the fixation of the segmental duplication, because one paralog became pseudogenized, which is a typical signature of the neutral or deleterious effect of a duplication. In contrast, the absence of pseudogenization in the *F10* and *F7* paralogs in our terrestrial elapid genomes, observed in not just one, but two segmental duplication events in the *P. textilis* lineage, suggests that selection has maintained these paralogs in an intact state.

After venom factor X was established as a procoagulant toxin early in the Australian elapid radiation, factor X’s endogenous cofactor, coagulation factor V, was recruited into the venom of the common ancestor of the *Pseudonaja*-*Oxyuranus* clade. For similar reasons as described for *F10* - the observed expression of the liver-expressed *F5* paralog in extant venom glands, the detection of coagulation factor V protein in venom, and the improbability of fixing a segmental duplication absent positive selection, we infer that venom factor V evolved through a similar series of genetic steps, the first being the expression of the single *F5* ortholog in venom glands (Fig. 7, step iv). Once *F5* expression was established in the venom gland, it would have enabled the formation of the prothrombinase complex with venom factor X. This new interaction would have then favored the preservation of the subsequent *F5*-containing segmental duplication (Fig. 7, step v).

Once these duplicated segments were preserved, new innovative mutations could then accumulate that optimize the expression and function of venom-expressed toxins without compromising the ancestral hemostatic clotting factors. One such mutation was the deletion within the activation peptide encoded by the *vF10* gene, which is suggested to render venom factor X constitutively active (neofunctionalization) (Fig. 7, step iii) (48). Since this deletion disrupts the hemostatic function of factor X (48), it may have also marked the evolutionary fate of one paralog to become the venom-expressed gene. We also found possible fate-determining mutations in one *F5* paralog in both *P. textilis* and *O*. *microlepidotus*, where the *vF5* gene acquired premature stop codon(s) in exon 13 (Fig. 7, step vi). These inactivating mutations prevent the translation of the full-length hemostatic protein, but do not affect the translation of the shorter venom protein, as this region of the RNA transcript is removed through alternative splicing.

This order of key events that we postulate for the genetic paths of venom factor X and venom factor V entailing: i) the co-option of the coagulation factor ortholog into venom; ii) segmental gene duplication; and iii) mutations that render the proteins constitutively active (Fig. 7, steps i-vi) are consistent with models that suggest that innovative mutations may precede gene duplication (60–62, 65). Indeed, co-option has occurred without gene duplication in the origin of some wasp (70) and snake venom toxins (71, 72).

### The neofunctionalization path

The genetic path of procoagulant PLA_₂_ proteins and plasmin-inhibiting Kunitz-type toxins did not start with *de novo* co-option of a physiological protein into venom. Rather, these venom proteins evolved by directly modifying venom-expressed genes in ways that enabled them to exert procoagulant activities (neofunctionalization) (Fig. 7, step vii).

We found striking parallels in the origins of the PLA_₂_ and Kunitz-type toxin proteins. Our phylogenetic analysis of group I PLA_₂_ loci suggests that the procoagulant variants evolved from a clade of genes encoding neurotoxins, of which the closest extant relative is a potent neurotoxin in *Notechis scutatus* (“Acidic PLA_₂_ HTe”) (55). We infer a similar evolutionary origin for the plasmin-inhibiting Kunitz-type proteins, which are also likely to have evolved from a neurotoxin ancestor. While none of the individual Kunitz-type toxins of Australian elapids have been functionally tested to date, the proposed neurotoxin ancestry is supported by several observations.

First, neurotoxic Kunitz-type toxins are found across elapids worldwide, from New World coral snakes (*Micrurus* spp.) (73), to African mambas (*Dendroaspis* spp.) (74, 75), and Asian kraits (*Bungarus* spp.) (57, 76), suggesting that neurotoxic forms were likely present in a common ancestor prior to the Australian elapid radiation. This inference is further supported by our finding that the KTSPI-A and KTSPI-C paralogs in the Asian krait (*Bungarus* candidus) appear to be the closest paralogs to all Australian elapid proteins, and that the KTSPI-C appears to be an ortholog of a potent potassium-channel blocking toxin in the banded krait (*Bungarus fasciatus*) (76). Finally, multiple residues that have been functionally shown to be important for binding and blocking potassium-channel activity (74, 75) are also present in the protein sequences of Australian elapid toxins examined in this study (*SI Appendix*, Fig. SX). For example, the key K5 and L9 residues that α-dendrotoxins use for channel binding are also found in Textilinin 7, Microlepidin 3 and Microlepidin 5 (75). The preservation of the L9 residue in all other Australian elapid proteins investigated here, including the plasmin-inhibitors (Textilinin 2a and Textilinin 2b), is consistent with an ancestral neurotoxin function. Therefore, we suggest that the procoagulant activities in Kunitz-type toxins and PLA_2_ toxins evolved through functional shifts in ancestral neurotoxins.

### Evolution takes the shorter path by working with the variation that is available

When similar traits evolve independently in unrelated groups of animals, they provide windows into the genetic paths that are possible for a given trait (77, 78). Whether such convergent traits evolved along the same genetic path or an entirely different one can help illuminate why evolution has unfolded in the ways it has. Some vipers such as Russell’s viper (*Daboia russelii*) independently evolved procoagulant venoms, so it is instructive to compare and contrast the genetic path of the origin of viper and elapid venoms. The procoagulant vipers did not co-opt their endogenous *F10* and *F5* genes, as the brown snakes/taipans did. Instead, these vipers evolved a way to activate the prey factor X and factor V proteins by modifying three pre-existing venom toxins: a metalloprotease and C-type lectin that form a complex that activates factor X, and a serine protease that activates prey factor V (79) (Fig. 8). Through different genetic paths, the elapid and viper toxins converge on the same functional outcome: prothrombin activation.

**Fig. 8.**
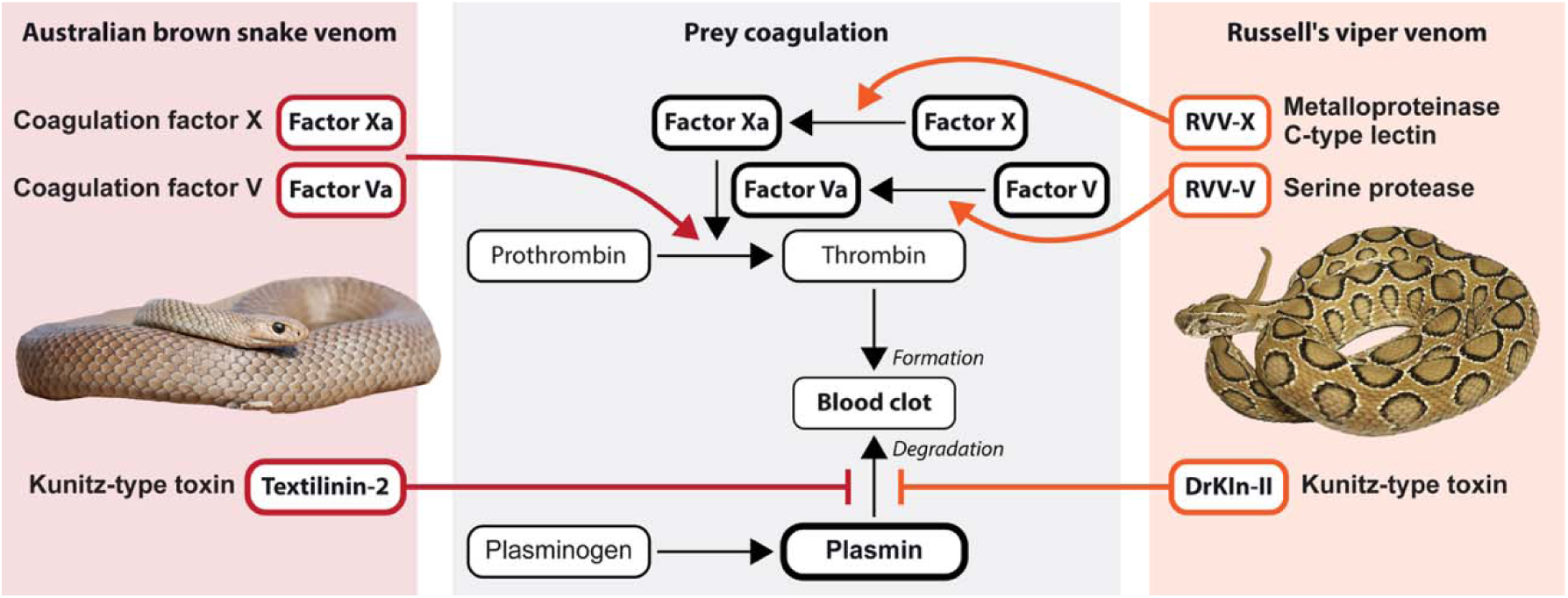
Convergent evolution of functionally analogous procoagulant venoms in brown snakes and certain vipers. The common pathway of the coagulation system involves the activation of factor X to activated factor X (Xa) and factor V to activated factor V (Va), which in turn form a complex that cleaves prothrombin to thrombin and promotes blood coagulation in prey (center panel, black arrows). Brown snake and taipan venoms activate the clotting system by injecting activated forms of factor X and factor V, thereby promoting thrombin generation (left panel, red arrow). In contrast, some viper venoms contain both a metalloproteinase that activates factor X and a serine protease that activates factor V, which together also promote thrombin generation (right panel, orange arrows). Blood clots are normally degraded by plasmin (center panel, black arrow), but both elapid and viper venoms contain Kunitz-type toxins that inhibit plasmin, thereby preventing the breakdown of blood clot (red and orange T-lines). [Photo credits: Eastern brown snake: Ákos Lumnitzer; Russell’s viper: Klaus Roemer.]

The different genomic origins of these functionally analogous toxins raise the question of why each snake lineage followed their respective genetic paths. We suggest that elapids and vipers took different paths due to differences in the genetic variation available for innovation in their respective ancestors. Viper genomes contain highly-expanded gene families of both venom-expressed metalloproteinases and serine proteases (23, 72, 80–82), several of which interact with components of the hemostatic system. Thus, vipers evolved prothrombinase activity by modifying extant toxins. By contrast, the Australian elapid *P*. *textilis* appears to possess only two venom metalloproteinase genes and no venom serine protease genes (72); the expansion and diversification of these toxin families is largely restricted to vipers. It appears then that no toxins were available in Australian elapid ancestors like those that existed in viper ancestors such that *de novo* co-option of the endogenous clotting factors was the only available path to evolving the ecologically novel prothrombin-activating toxins.

However, both elapids and vipers took very similar paths to evolving plasmin-inhibiting venom components. To prevent clot breakdown, both snake lineages evolved venom plasmin-inhibitors - not by *de novo* co-option of an endogenous plasmin inhibitor, but by modifying venom-expressed Kunitz-type toxins (79).

We suggest that this observation is best explained by the shorter mutational path being the more probable path to the evolution of novelty. Kunitz-type toxins are encoded by a multigene family of venom-expressed serine protease inhibitors that can interact with a broad range of serine proteases (44, 83, 84). Because of this apparent promiscuity, a small number of mutations may have been sufficient to shift their activity and specificity toward plasmin inhibition (i.e. R17 and V18) (85). In contrast, the *de novo* co-option of a liver-expressed plasmin inhibitor to be expressed in the venom gland may require certain kinds of mutational events (the gain of a venom gland enhancer) that may occur less frequently than substitutions in venom-expressed proteins.

The inference that *de novo* co-option is the less available path compared to the modification of existing toxins is also supported by two observations about the evolution of snake venoms in general. First, that key snake venom toxin families are descended and co-opted from a single member of an ancestral gene family, and second, we do not observe additional co-option events within the same protein family in a given lineage. For example, in elapid venoms, three-finger toxins evolved from a squamate-specific *LY6* gene (25) and phospholipases from group I *PLAG1B* (this study), while in viper venoms, their metalloproteinases evolved from *ADAM28* (23), phospholipases from group II *Pla2g* (22), and serine proteases from a reptilian kallikrein gene (24). The expansion and diversification of these toxin families in the two major venomous snake lineages have occurred exclusively through changes in the number and structures of the venom-expressed gene clade, rather than via the independent co-option of any of the numerous other metalloproteinase, phospholipase, or serine protease genes in their genomes.

Thus, we can appreciate that when these relatively rare *de novo* co-option events do occur, they can have an outsized impact by pioneering new venom activities that exploit new ecological opportunities. The *de novo* co-options of the five ancestral genes noted above were seminal events in the evolution and radiation of elapids and vipers, as were the *de novo* co-option of *F10* in what has been described as one of the most prolific radiations of venomous snakes (28), and of *F5* in endowing the brown snake-taipan clade with the most potent of all venoms.

## Materials and Methods

### Blood and tissue collection

Inland taipan (*Oxyuranus microlepidotus*) and Malayan krait (*Bungarus candidus*) blood were provided by Kentucky Reptile Zoo (Slade, Kentucky, USA). Eastern brown snake (*Pseudonaja textilis*) blood was provided by the Australian Reptile Academy (Whiterock, Queensland, Australia). This specimen was a captive adult male originating from Cryna, southeast Queensland. Furthermore, a captive-bred male *O. microlepidotus* was kept at the Viper Resource Center of the National Natural Toxins Research Center, Texas A&M University-Kingsville (Kingsville, Texas, USA), and its tissues were collected in accordance with the protocols reviewed and approved by the Texas A&M Kingsville Institutional Animal Care and Use Committee in compliance with all applicable federal regulations governing the protection of research animals (Viper Resource Center at Texas A&M University-Kingsville, IACUC #2018-11-09-A3 and #:2021-11-29/1474). The venom gland and liver tissues were dissected, cut into 0.5 cm pieces, and stored in 1800 µL of RNAprotect Tissue Reagent (Ref. 76104. QIAGEN, Hilden, Germany) at 4 °C.

### DNA isolation, library preparation, and DNA sequencing

High-molecular-weight DNA was isolated from *O. microlepidotus* and *B. candidus* blood with the NucleoBond HMW DNA kit (Ref. 740160.20; Macherey-Nagel, Düren, Germany) according to the manufacturer’s instructions, and its purity and integrity was validated using gel electrophoresis.

Subsequently, library preparation and DNA sequencing were performed by Cold Spring Harbor Laboratory (Laurel Hollow, New York, USA). For *P*. *textilis*, high-molecular-weight DNA extraction, library preparation, and genome sequencing were commercially performed by the Australian Genome Research Facility (Melbourne, Victoria, Australia). The input DNA for the sequencing libraries was size selected for fragments of ∼15 kilo base pairs. The libraries were then sequenced using Pacific Biosciences HiFi technology on a Revio system, with three 25M SMRT Cells for *O*. *microlepidotus* and *P*. *textilis*, or two 25M SMRT Cells for *B*. *candidus*.

### RNA isolation, library preparation, and RNA sequencing

Total RNA was isolated from *O. microlepidotus* venom gland and liver tissue using RNeasy Plus Mini Kit (Ref. 74134. QIAGEN, Hilden, Germany) according to the manufacturer’s instructions, and its purity and integrity was validated using gel electrophoresis. Each tissue was processed separately to prevent cross-contamination. Library preparation and mRNA sequencing were performed by Cold Spring Harbor Laboratory (Laurel Hollow, New York, USA) using Illumina short-read and PacBio long-read technologies. For short-read Illumina RNA sequencing, two Kapa mRNA libraries were prepared, and each library (representing one tissue) was sequenced (2 x 100 cycles) on a NextSeq 550 system. For Pacific Biosciences Iso-Seq sequencing, two SMRTbell Express kits were used to prepare libraries, and each library (representing one tissue) was sequenced in a PacBio Sequel II cell using 30-hour movie times. Reads were processed using the Pacific Biosciences processing pipeline (https://isoseq.how). The data processing included five steps: consensus generation, demultiplexing, refinement, clustering and collapsing of mapped reads. The genome mapping of clustered reads permitted the identification of sets of gene associated read clusters. Collapsing of many reads into a single consensus sequence occurred only within clustered read sets.

### Genome assembly

The *O. microlepidotus* and *P. textilis* genomes were assembled using Canu (version 2.1.1) (86) with the settings “genomeSize=1.8G” for estimated genome size and “pacbio-hifi” for the input reads. The *B*. *candidus* genome was assembled using HIFIasm (version 0.25.0-r726) (87). Because the *F10* gene region was absent from existing *N*. *scutatus* assemblies, despite transcript and protein evidence indicating its presence in this species (30, 32, 46), we reassembled the genome using an alternative assembler to recover this missing region. To do so, Oxford Nanopore Technologies (ONT) genomic reads (SRR30595366) were retrieved from the Sequence Read Archive (SRA) and assembled with Canu (version 2.3) (88) using the settings “genomeSize=1.8G” and “-nanopore”.

### Annotation of venom loci

Homology-based manual gene annotations were performed on both newly generated and publicly available genome assemblies. The barred grass snake (*Natrix helvetica*) assembly (GCF_964273705.1) was used as the non-venomous outgroup species for the *F10*, *F5*, and *PLA2G1B* loci. The newly generated assembly for *N*. *scutatus* was used exclusively for the *F10* region, whereas an existing assembly (GCA_049901735.1) was used for the other venom gene families. The *F10* gene region was also annotated in several publicly available genomes, including *Aipysurus laevis* (GCA_040207615.1) (89), *Hydrophis major* (GCA_033807585.1) (90), *Hydrophis elegans* (GCA_033807725.1) (90), *Hydrophis curtus* (GCA_037043045.1) (90), and *Hydrophis ornatus* (GCA_037055225.1) (90). Furthermore, the Kunitz-type toxin locus was also annotated in *Charina bottae* (GCA_023362775.1) (91). First, we used the “tblastn” command to identify the contigs, scaffolds, or chromosomes containing candidate venom gene(s) by comparison with reference sequences (92). In these genomic regions, we manually annotated the venom genes and their flanking genes using SnapGene (version 8.0) (GSL Biotech LLC, San Diego, Calfornia, USA). The exons were manually identified through an iterative process of either “blastn” or “tblastn”, with reference protein or transcript sequences as queries, and were later refined by identifying exon–intron boundaries. Because of the error-prone nature of the *N*. *scutatus* ONT reads, small exonic indels that disrupted an otherwise intact reading frames were manually corrected. Most of these indels occurred in homopolymeric regions and were therefore considered to be technical artifacts rather than true biological variation. Finally, we obtained the coding DNA sequences (CDSs), corresponding to the mRNA coding regions, and their predicted protein translations using the “Splice to Remove Introns…” function implemented in SnapGene.

### Transcriptomics

To qualitatively assess whether the B-domains of both *F5* paralogs are alternatively spliced, we analyzed full-length RNA transcripts in both venom gland and liver tissues of *O*. *microlepidotus*. From this curated Isoseq dataset, all *F5* transcripts were identified using the “blastn” command with the *F5* paralogs from *O*. *microlepidotus* genome as the query sequences (92). The identified transcripts were extracted, aligned with *F5* gene annotations in AliView (version 1.30) (93), and subsequently manually analyzed for variation in exon 13 (which encodes the entire B-domain).

To quantify relative expression levels of *F10*, *F7* and *F5* genes and respective isoforms, we counted full-length RNA reads in the venom gland and liver tissues of *O*. *microlepidotus*. We used full-length RNA transcript counts as a semi-quantitative proxy for their gene expression, which allowed us to unambiguously identify and discriminate between highly similar paralogs and isoforms. All transcripts corresponding to each gene were identified using the “blastn” command (92), extracted, aligned with annotated CDSs in AliView (93), and quantified by the same software, with manual verification of transcript identity to prevent any misassignments. To validate this approach, we used several known liver-expressed genes (*F2*, *F9*, *PROZ* and *ALB*) as controls.

To verify venom gland expression of annotated venom toxin genes other than *F10*, *F5*, and *F7*, short RNA reads were mapped to the annotated CDSs. RNA-seq data from *O*. *microlepidotus* were generated in this study, whereas equivalent data for adult *P*. *textilis* were retrieved from the SRA database (accessions SRR33084406 and SRR33084395). First, low-quality reads and adapter sequences were removed using Trimmomatic (version 0.40-rc1) (94) with the following settings: “ILLUMINACLIP:TruSeq3-PE-2.fa:2:30:10”, “LEADING:5”, “TRAILING:5”, “SLIDINGWINDOW:4:30”, “MINLEN:40”. Next, we evaluated read quality and confirmed that adapters had been correctly removed using FastQC (version 0.12.1). We then prepared transcript reference sequences from CDSs of all annotated genes using RSEM (version v1.3.1) (95). Subsequently, reads were mapped using RSEM with Bowtie2 (version 2.4.4) (96) with the sensitivity level set to “very sensitive” for the CDS datasets to estimate relative gene expression levels in venom gland and liver tissues of *O*. *microlepidotus* and *P. textilis*.

### Proteomics

In order to search for evidence of the hemostatic clotting factor proteins (Factor X, Factor V, and Factor VII) in venoms, we reanalyzed publicly available proteomic data from seven *P*. *textilis* venom samples. Raw mass spectrometry data from trypsin-digested venom protein peptides were retrieved from the PRoteomics IDEntifications (PRIDE) database (dataset identifier PXD063001). Fragmentation spectra were searched against hypothetical protein translations derived from our *P*. *textilis* gene annotations using the MSFragger-based FragPipe computational platform (97, 98).

### Phylogenetic analysis

Bayesian phylogenetic analyses were performed for factor X, group I PLA_2_, and Kunitz-type toxins. Hypothetical protein translations from our genome annotations were aligned using the Multiple Sequence Comparison by Log-Expectation (MUSCLE) algorithm (99) in AliView (version 1.30) (93). Phylogenetic analyses were conducted in MrBayes (version 3.2.7a) (100). Signal peptides of group I PLA_2_ sequences were manually removed from the alignments. Amino-acid substitution models were sampled using Bayesian model averaging (aamodelpr = mixed). The Markov chain Monte Carlo algorithm was run for 15 million generations, sampling every 100 generations, with four independent runs of four chains each (three hot, one cold), a heating temperature of 0.2, and a burn-in of 25%. All analyses reached convergence, as indicated by meeting the convergence threshold (stopval = 0.01). The consensus phylogenies were constructed using contype=halfcompat (for factor X and group I PLA_₂_) and contype=allcompat (for Kunitz-type toxins).

## Acknowledgments

We thank Christina N. Zdenek and Chris Hay of Australian Reptile Academy and Kristen Wiley of Kentucky Reptile Zoo for providing blood samples; Josh Llinas of Unusual Pet Vets for collecting the Eastern brown snake blood sample; Sara Goodwin of the Cold Spring Harbor Sequencing Core for technical consultation; Lucas Trevisan of the Australian Genome Research Facility for assistance with DNA sequencing; Somasekar Seshagiri for discussion of snake genomics; the staff of the Texas A&M University-Kingsville National Natural Toxins Research Center for their technical assistance and support; and Matt Giorgianni, Fiona Ukken, Yetunde Ayinuola, and Rabindra Thakur for technical advice and comments on the manuscript. This work was supported by the Viper Resource Grant at Texas A&M University-Kingsville (#P40OD010960-22; E.E.S), the Howard Hughes Medical Institute (S.B.C.) and by the Andrew and Mary Balo and Nicholas and Susan Simon Endowed Professorship in Biology at the University of Maryland (S.B.C.).

## Data, Materials and Software availability

The whole-genome PacBio HiFi reads, short-read RNA-sequencing (Illumina RNA-seq) data, and long-read RNA sequencing (PacBio Iso-seq) data have been deposited in the National Center for Biotechnology Information Sequence Read Archive (SRA) and are available under the BioProject accession number PRJNAXXXXXXX. This Whole Genome Shotgun project has been deposited at DDBJ/ENA/GenBank under the accession XXXXXX000000000. The version described in this paper is version XXXXXX010000000.

## Author Contributions

J.v.T. and S.B.C. designed research; J.v.T., N.L.D., and E.E.S, performed research; J.v.T., and C.F.S. analyzed data; and J.v.T. and S.B.C. wrote the paper.

## References

1. S. B. Carroll, Evo-Devo and an Expanding Evolutionary Synthesis: A Genetic Theory of Morphological Evolution. Cell 134, 25–36 (2008).

2. G. P. Wagner, V. J. Lynch, Evolutionary novelties. Current Biology 20, R48–R52 (2010).

3. N. Shubin, C. Tabin, S. Carroll, Deep homology and the origins of evolutionary novelty. Nature 457, 818–823 (2009).

4. A. P. Moczek, D. J. Rose, Differential recruitment of limb patterning genes during development and diversification of beetle horns. Proceedings of the National Academy of Sciences 106, 8992–8997 (2009).

5. T. Werner, S. Koshikawa, T. M. Williams, S. B. Carroll, Generation of a novel wing colour pattern by the Wingless morphogen. Nature 464, 1143–1148 (2010).

6. R. D. Reed et al., optix Drives the Repeated Convergent Evolution of Butterfly Wing Pattern Mimicry. Science 333, 1137–1141 (2011).

7. W. J. Glassford et al., Co-option of an Ancestral Hox-Regulated Network Underlies a Recently Evolved Morphological Novelty. Developmental Cell 34, 520–531 (2015).

8. F. Leal, Martin J. Cohn, Loss and Re-emergence of Legs in Snakes by Modular Evolution of Sonic hedgehog and HOXD Enhancers. Current Biology 26, 2966–2973 (2016).

9. E. Z. Kvon et al., Progressive Loss of Function in a Limb Enhancer during Snake Evolution. Cell 167, 633–642.e611 (2016).

10. S. N. Murugesan et al., Butterfly eyespots evolved via cooption of an ancestral gene-regulatory network that also patterns antennae, legs, and wings. Proceedings of the National Academy of Sciences 119, e2108661119 (2022).

11. R. T. Kellenberger et al., Multiple gene co-options underlie the rapid evolution of sexually deceptive flowers in Gorteria diffusa. Current Biology 33, 1502–1512.e1508 (2023).

12. J. W. Satterlee et al., Convergent evolution of plant prickles by repeated gene co-option over deep time. Science 385, eado1663 (2024).

13. A. L. Herbert et al., Ancient developmental genes underlie evolutionary novelties in walking fish. Current Biology 34, 4339–4348.e4336 (2024).

14. E. McQueen, M. Rebeiz, “Chapter Twelve - On the specificity of gene regulatory networks: How does network co-option affect subsequent evolution?” in Current Topics in Developmental Biology, I. S. Peter, Ed. (Academic Press, 2020), vol. 139, pp. 375–405.

15. N. Gompel, B. Prud’homme, P. J. Wittkopp, V. A. Kassner, S. B. Carroll, Chance caught on the wing: cis-regulatory evolution and the origin of pigment patterns in Drosophila. Nature 433, 481–487 (2005).

16. L. Arnoult et al., Emergence and Diversification of Fly Pigmentation Through Evolution of a Gene Regulatory Module. Science 339, 1423–1426 (2013).

17. S. Ohno, Evolution by gene duplication (Springer Science & Business Media, 1970).

18. G. C. Conant, K. H. Wolfe, Turning a hobby into a job: How duplicated genes find new functions. Nature Reviews Genetics 9, 938–950 (2008).

19. M. Soskine, D. S. Tawfik, Mutational effects and the evolution of new protein functions. Nature Reviews Genetics 11, 572–582 (2010).

20. A. F. Cisneros et al., Evolutionary causes and consequences of gene duplication. Nature Reviews Genetics 10.1038/s41576-026-00935-5 (2026).

21. N. R. Casewell, W. Wüster, F. J. Vonk, R. A. Harrison, B. G. Fry, Complex cocktails: the evolutionary novelty of venoms. Trends in Ecology & Evolution 28, 219–229 (2013).

22. N. L. Dowell et al., The Deep Origin and Recent Loss of Venom Toxin Genes in Rattlesnakes. Current Biology 26, 2434–2445 (2016).

23. M. W. Giorgianni et al., The origin and diversification of a novel protein family in venomous snakes. Proceedings of the National Academy of Sciences 117, 10911–10920 (2020).

24. A. Barua, I. Koludarov, A. S. Mikheyev, Co-option of the same ancestral gene family gave rise to mammalian and reptilian toxins. BMC Biology 19, 268 (2021).

25. I. Koludarov et al., Domain loss enabled evolution of novel functions in the snake three-finger toxin gene superfamily. Nature Communications 14, 4861 (2023).

26. J. D. Galbraith et al., Horizontal Transposon Transfer and Its Implications for the Ancestral Ecology of Hydrophiine Snakes. 10.3390/genes13020217.

27. H. Harold, G. Alana, M. Helene, Paleoclimatology, Paleogeography, and the Evolution and Distribution of Sea Kraits (Serpentes; Elapidae; Laticauda). Herpetological Monographs 31, 1–17 (2017).

28. K. L. Sanders, M. S. Y. Lee, R. Leys, R. Foster, J. Scott Keogh, Molecular phylogeny and divergence dates for Australasian elapids and sea snakes (hydrophiinae): evidence from seven genes for rapid evolutionary radiations. Journal of Evolutionary Biology 21, 682–695 (2008).

29. P. M. Oliver, A. F. Hugall, Phylogenetic evidence for mid-Cenozoic turnover of a diverse continental biota. Nature Ecology & Evolution 1, 1896–1902 (2017).

30. G. Tans, J. W. Govers-Riemslag, J. L. van Rijn, J. Rosing, Purification and properties of a prothrombin activator from the venom of Notechis scutatus scutatus. Journal of Biological Chemistry 260, 9366–9372 (1985).

31. H. Speijer, J. Govers-Riemslag, R. Zwaal, J. Rosing, Prothrombin activation by an activator from the venom of Oxyuranus scutellatus (Taipan snake). Journal of Biological Chemistry 261, 13258–13267 (1986).

32. V. S. Rao, J. S. Joseph, R. M. Kini, Group D prothrombin activators from snake venom are structural homologues of mammalian blood coagulation factor Xa. Biochemical Journal 369, 635–642 (2003).

33. V. S. Rao, S. Swarup, M. R. Kini, The catalytic subunit of pseutarin C, a group C prothrombin activator from the venom of Pseudonaja textilis, is structurally similar to mammalian blood coagulation factor Xa. Thromb Haemost 92, 509–521 (2004).

34. M. A. Reza, S. Swarup, M. R. Kini, Two parallel prothrombin activator systems in Australian rough-scaled snake, Tropidechis carinatus. Structural comparison of venom prothrombin activator with blood coagulation factor X 93, 40–47 (2005).

35. M. A. Reza, T. N. Minh Le, S. Swarup, R. Manjunatha Kini, Molecular evolution caught in action: gene duplication and evolution of molecular isoforms of prothrombin activators in Pseudonaja textilis (brown snake). Journal of Thrombosis and Haemostasis 4, 1346–1353 (2006).

36. C. N. Zdenek et al., Clinical implications of convergent procoagulant toxicity and differential antivenom efficacy in Australian elapid snake venoms. Toxicology Letters 316, 171–182 (2019).

37. T. N. W. Jackson et al., Rapid Radiations and the Race to Redundancy: An Investigation of the Evolution of Australian Elapid Snake Venoms. 10.3390/toxins8110309.

38. V. S. Rao, S. Swarup, R. M. Kini, The nonenzymatic subunit of pseutarin C, a prothrombin activator from eastern brown snake (Pseudonaja textilis) venom, shows structural similarity to mammalian coagulation factor V. Blood 102, 1347–1354 (2003).

39. T. N. Minh Le, M. A. Reza, S. Swarup, R. M. Kini, Gene duplication of coagulation factor V and origin of venom prothrombin activator in Pseudonaja textilis snake. Thromb Haemost 93, 420–429 (2005).

40. M. E. Nesheim, J. B. Taswell, K. G. Mann, The contribution of bovine Factor V and Factor Va to the activity of prothrombinase. Journal of Biological Chemistry 254, 10952–10962 (1979).

41. A. Armugam et al., Group IB phospholipase A2 from Pseudonaja textilis. Archives of Biochemistry and Biophysics 421, 10–20 (2004).

42. P. P. Masci et al., Textilinins from Pseudonaja textilis textilis. Characterization of two plasmin inhibitors that reduce bleeding in an animal model. Blood Coagulation & Fibrinolysis 11 (2000).

43. I. Filippovich et al., A family of textilinin genes, two of which encode proteins with antihaemorrhagic properties. British Journal of Haematology 119, 376–384 (2002).

44. S. T. H. Earl et al., Identification and characterisation of Kunitz-type plasma kallikrein inhibitors unique to Oxyuranus sp. snake venoms. Biochimie 94, 365–373 (2012).

45. A. J. Broad, S. K. Sutherland, A. R. Coulter, The lethality in mice of dangerous Australian and other snake venom. Toxicon 17, 661–664 (1979).

46. L. St. Pierre et al., Comparative Analysis of Prothrombin Activators from the Venom of Australian Elapids. Molecular Biology and Evolution 22, 1853–1864 (2005).

47. M. Hertzberg, Biochemistry of factor X. Blood Reviews 8, 56–62 (1994).

48. M. Schreuder et al., Evolutionary Adaptations in Pseudonaja Textilis Venom Factor X Induce Zymogen Activity and Resistance to the Intrinsic Tenase Complex. Thromb Haemost 120, 1512–1523 (2020).

49. H. Zaher et al., Large-scale molecular phylogeny, morphology, divergence-time estimation, and the fossil record of advanced caenophidian snakes (Squamata: Serpentes). PLOS ONE 14, e0216148 (2019).

50. D. Verhoef et al., Evolutionary conservation of regulatory B-domain elements in blood coagulation factor V in snakes. Lessons from snake venom: New insights into the structural and functional aspects of factor V and factor X, 25 (2021).

51. M. H. A. Bos et al., Venom factor V from the common brown snake escapes hemostatic regulation through procoagulant adaptations. Blood 114, 686–692 (2009).

52. J. A. Pearson, M. I. Tyler, K. V. Retson, M. E. H. Howden, Studies on the subunit structure of textilotoxin, a potent presynaptic neurotoxin from the venom of the Australian common brown snake (Pseudonaja textilis). 3. The complete amino-acid sequences of all the subunits. Biochimica et Biophysica Acta (BBA) - Protein Structure and Molecular Enzymology 1161, 223–229 (1993).

53. J. Fohlman, D. Eaker, E. Karlsson, S. Thesleff, Taipoxin, an Extremely Potent Presynaptic Neurotoxin from the Venom of the Australian Snake Taipan (Oxyuranus s. scutellatus). European Journal of Biochemistry 68, 457–469 (1976).

54. E. Karlsson, D. Eaker, L. Rydén, Purification of a presynaptic neurotoxin from the venom of the Australian tiger snake Notechis scutatus scutatus. Toxicon 10, 405–413 (1972).

55. B. Francis, J. A. Coffield, L. L. Simpson, I. I. Kaiser, Amino Acid Sequence of a New Type of Toxic Phospholipase A2 from the Venom of the Australian Tiger Snake (Notechis scutatus scutatus). Archives of Biochemistry and Biophysics 318, 481–488 (1995).

56. O. Khow et al., Isolation, Toxicity and Amino Terminal Sequences of Three Major Neurotoxins in the Venom of Malayan Krait (Bungarus candidus) from Thailand. The Journal of Biochemistry 134, 799–804 (2003).

57. C. G. Benishin, Potassium channel blockade by the B subunit of beta-bungarotoxin. Molecular Pharmacology 38, 164–169 (1990).

58. J. B. Walsh, How often do duplicated genes evolve new functions? Genetics 139, 421–428 (1995).

59. A. Force et al., Preservation of Duplicate Genes by Complementary, Degenerative Mutations. Genetics 151, 1531–1545 (1999).

60. U. Bergthorsson, D. I. Andersson, J. R. Roth, Ohno’s dilemma: Evolution of new genes under continuous selection. Proceedings of the National Academy of Sciences 104, 17004–17009 (2007).

61. A. L. Hughes, The evolution of functionally novel proteins after gene duplication. Proceedings of the Royal Society B: Biological Sciences 256, 119–124 (1994).

62. J. Piatigorsky, G. Wistow, The Recruitment of Crystallins: New Functions Precede Gene Duplication. Science 252, 1078–1079 (1991).

63. C. T. Hittinger, S. B. Carroll, Gene duplication and the adaptive evolution of a classic genetic switch. Nature 449, 677–681 (2007).

64. D. L. Des Marais, M. D. Rausher, Escape from adaptive conflict after duplication in an anthocyanin pathway gene. Nature 454, 762–765 (2008).

65. J. Näsvall, L. Sun, J. R. Roth, D. I. Andersson, Real-Time Evolution of New Genes by Innovation, Amplification, and Divergence. Science 338, 384–387 (2012).

66. Y.-C. Tang, A. Amon, Gene Copy-Number Alterations: A Cost-Benefit Analysis. Cell 152, 394–405 (2013).

67. A. M. Rice, A. McLysaght, Dosage-sensitive genes in evolution and disease. BMC Biology 15, 78 (2017).

68. R. Koszul, B. Dujon, G. Fischer, Stability of Large Segmental Duplications in the Yeast Genome. Genetics 172, 2211–2222 (2006).

69. M. J. Margres, A. T. Bigelow, E. M. Lemmon, A. R. Lemmon, D. R. Rokyta, Selection To Increase Expression, Not Sequence Diversity, Precedes Gene Family Origin and Expansion in Rattlesnake Venom. Genetics 206, 1569–1580 (2017).

70. E. O. Martinson, Mrinalini, Y. D. Kelkar, C.-H. Chang, J. H. Werren, The Evolution of Venom by Co-option of Single-Copy Genes. Current Biology 27, 2007–2013.e2008 (2017).

71. C.-T. Pan et al., The evolution and structure of snake venom phosphodiesterase (svPDE) highlight its importance in venom actions. eLife 12, e83966 (2023).

72. D. D. Almeida et al., Tracking the recruitment and evolution of snake toxins using the evolutionary context provided by the Bothrops jararaca genome. Proceedings of the National Academy of Sciences 118, e2015159118 (2021).

73. C. J. Bohlen et al., A heteromeric Texas coral snake toxin targets acid-sensing ion channels to produce pain. Nature 479, 410–414 (2011).

74. J. P. Imredy, R. MacKinnon, Energetic and structural interactions between δ-dendrotoxin and a voltage-gated potassium channel11Edited by G. von Heijne. Journal of Molecular Biology 296, 1283–1294 (2000).

75. S. Gasparini et al., Delineation of the functional site of alpha-dendrotoxin: The functional topographies of dendrotoxins are different but share a conserved core with those of other Kv1 potassium channel-blocking toxins. Journal of Biological Chemistry 273, 25393–25403 (1998).

76. W. Yang et al., BF9, the First Functionally Characterized Snake Toxin Peptide with Kunitz-Type Protease and Potassium Channel Inhibiting Properties. Journal of Biochemical and Molecular Toxicology 28, 76–83 (2014).

77. Z. D. Blount, R. E. Lenski, J. B. Losos, Contingency and determinism in evolution: Replaying life’s tape. Science 362, eaam5979 (2018).

78. J. B. Losos, Improbable destinies: Fate, chance, and the future of evolution (Penguin, 2018).

79. B. Xie et al., Dynamic genetic differentiation drives the widespread structural and functional convergent evolution of snake venom proteinaceous toxins. BMC Biology 20, 4 (2022).

80. D. R. Schield et al., The origins and evolution of chromosomes, dosage compensation, and mechanisms underlying venom regulation in snakes. Genome Research 10.1101/gr.240952.118, - (2019).

81. G. Mochales-Riaño et al., Chromosome-level reference genome for the medically important Arabian horned viper (Cerastes gasperettii). GigaScience 14, giaf030 (2025).

82. A. J. Mason et al., Molecular mechanisms underlying early functional divergence in snake venom inferred from the genomes of two pitviper lineages. BMC Biology 23, 366 (2025).

83. A. Ritonja, V. Turk, F. GubenŠEk, Serine Proteinase Inhibitors from Vipera ammodytes Venom. European Journal of Biochemistry 133, 427–432 (1983).

84. A.-C. Cheng, I.-H. Tsai, Functional characterization of a slow and tight-binding inhibitor of plasmin isolated from Russell’s viper venom. Biochimica et Biophysica Acta (BBA) - General Subjects 1840, 153–159 (2014).

85. E.-K. I. Millers et al., The structure of Human Microplasmin in Complex with Textilinin-1, an Aprotinin-like Inhibitor from the Australian Brown Snake. PLOS ONE 8, e54104 (2013).

86. S. Nurk et al., HiCanu: accurate assembly of segmental duplications, satellites, and allelic variants from high-fidelity long reads. Genome Research 30, 1291–1305.

87. H. Cheng, G. T. Concepcion, X. Feng, H. Zhang, H. Li, Haplotype-resolved de novo assembly using phased assembly graphs with hifiasm. Nature Methods 18, 170–175 (2021).

88. S. Koren et al., Canu: scalable and accurate long-read assembly via adaptive k-mer weighting and repeat separation. Genome Research 10.1101/gr.215087.116, gr.215087.215116 (2017).

89. I. H. Rossetto, A. J. Ludington, B. F. Simões, N. Van Cao, K. L. Sanders, Dynamic Expansions and Retinal Expression of Spectrally Distinct Short-Wavelength Opsin Genes in Sea Snakes. Genome Biology and Evolution 16 (2024).

90. A. J. Ludington, J. M. Hammond, J. Breen, I. W. Deveson, K. L. Sanders, New chromosome-scale genomes provide insights into marine adaptations of sea snakes (Hydrophis: Elapidae). BMC Biology 21, 284 (2023).

91. J. L. Grismer et al., Reference genome of the rubber boa, Charina bottae (Serpentes: Boidae). Journal of Heredity 113, 641–648 (2022).

92. S. F. Altschul et al., Gapped BLAST and PSI-BLAST: a new generation of protein database search programs. Nucleic Acids Research 25, 3389–3402 (1997).

93. A. Larsson, AliView: a fast and lightweight alignment viewer and editor for large datasets. Bioinformatics 30, 3276–3278 (2014).

94. A. M. Bolger, M. Lohse, B. Usadel, Trimmomatic: a flexible trimmer for Illumina sequence data. Bioinformatics 30, 2114–2120 (2014).

95. B. Li, C. N. Dewey, RSEM: accurate transcript quantification from RNA-Seq data with or without a reference genome. BMC Bioinformatics 12, 323 (2011).

96. B. Langmead, S. L. Salzberg, Fast gapped-read alignment with Bowtie 2. Nature Methods 9, 357–359 (2012).

97. A. T. Kong, F. V. Leprevost, D. M. Avtonomov, D. Mellacheruvu, A. I. Nesvizhskii, MSFragger: ultrafast and comprehensive peptide identification in mass spectrometry–based proteomics. Nature Methods 14, 513–520 (2017).

98. F. Yu et al., Identification of modified peptides using localization-aware open search. Nature Communications 11, 4065 (2020).

99. R. C. Edgar, MUSCLE: multiple sequence alignment with high accuracy and high throughput. Nucleic Acids Research 32, 1792–1797 (2004).

100. F. Ronquist et al., MrBayes 3.2: Efficient Bayesian Phylogenetic Inference and Model Choice Across a Large Model Space. Systematic Biology 61, 539–542 (2012).

101. N. L. Bray, H. Pimentel, P. Melsted, L. Pachter, Near-optimal probabilistic RNA-seq quantification. Nature Biotechnology 34, 525–527 (2016).

