## Supplementary Information for "The genomic origins and evolutionary path to a key innovation in the world’s most venomous snakes"

**This PDF file includes:**

Supporting text
Figures S1 to S16
Tables S1 to S2
SI References

**Supporting text**

*Short-read RNA-seq mapping*

To search for evidence of *F10* and *F5* expression in *P. textilis* venom gland tissue, we mapped short RNA-seq reads to the corresponding genomic loci. Adult *P. textilis* venom gland RNA-seq data were retrieved from the SRA database (accession SRR33084395). Low-quality reads and adapter sequences were removed using Trimmomatic (version 0.40-rc1) (1) with the following parameters: “ILLUMINACLIP:TruSeq3-PE-2.fa:2:30:10”, “LEADING:5”, “TRAILING:5”, “SLIDINGWINDOW:4:30”, and “MINLEN:40”. Read quality was subsequently assessed, and adapter removal was confirmed using FastQC (version 0.12.1). The processed RNA-seq reads were aligned to all contigs in our *P. textilis* genome assembly using the BWA-MEM algorithm (2). Using the Samtools “view” command (3), paired-end reads with a mapping quality (MAPQ) score greater than 20 were retained and sorted, and only the reads mapping to the two contigs containing the *F10* and *F5* genes were extracted. Mapped reads were subsequently visualized and inspected using Integrative Genomics Viewer (IGV; version 2.16.0) (4, 5) to search for reads mapping to regions unique to the *F10* or *F5* genes and not to their respective paralogs.

### Figures

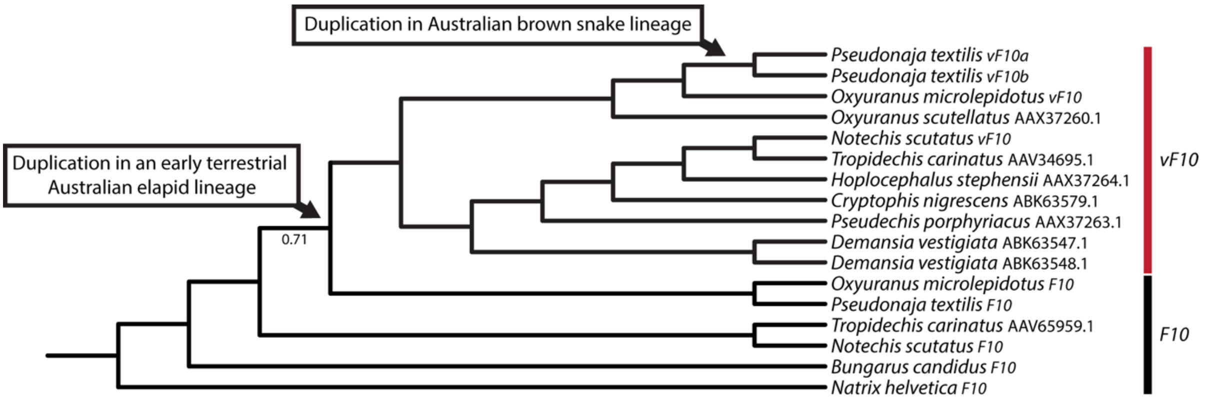

**Fig. S1. A venom-expressed *F10* gene was duplicated in the brown snake lineage (*Pseudonaja* spp.).** A protein phylogeny based on hypothetical translations of the *F10* genes and full-length cDNA sequences from distinct Australian elapid species. At the base of the phylogeny, there is a cluster of *F10* genes (encoding the hemostatic protein), and nested within it, a well-supported monophyletic clade consisting of *vF10* (encoding the venom toxin). In the venom gene clade, the *vF10b* gene clusters with *vF10a*, indicating that *vF10b* arose through duplication of the venom-expressed *vF10a* and not from the liver-expressed *F10* gene, a scenario that has previously been proposed (6). The phylogeny was rooted with *N. helvetica F10* protein sequence as the outgroup. Only Bayesian posterior probabilities below 1.00 are shown.

**1 — 76**

| | |
|---|---|
| *Pseudonaja textilis vF10a* | MAPQLLLCLILTFLWSLPEAESNVFLKSKVANRFLQRTKRANSLVEEFKSGNIERECIEERCSKEEAREVFEDDEK |
| *Pseudonaja textilis vF10b* | MAPQLLLCLILTFLWSLPEAESNVFLKSKVANRFLQRTKRANSLVEEFKSGNIERECIEERCSKEEAREVFEDDEK |
| *Oxyuranus microlepidotus vF10* | MAPQLLLCLILTFLWSLPEAESNVFLKSKVANRFLQRTKRANSLFEEFRSGNIERECIEERCSKEEAREVFEDDEK |
| *Notechis scutatus vF10* | MAPQLLLCLILTFLWSLPEAESNVFLKSKVANRFLQRTKRSNSLFEEIRPGNIERECIEEKCSKEEAREVFEDNEK |
| *Pseudonaja textilis F10* | MAPQLLLCLILTFLWSLPEAESNVFLKSKVANRFLQRTKRANSLFEEFKPGNIERECIEERCSKEEAREAFEDDEK |
| *Oxyuranus microlepidotus F10* | MAPQLLLCLILTFLWSLPEAESNVFLKSKVANRFLQRTKRANSLFEEFKPGNIERECIEERCSKEEAREAFEDDEK |
| *Notechis scutatus F10* | MAPQLLLCLILTFLWSLPEAESNVFLKSKVANRFLQRTKRANSLFEEFKAGNIERECIEERCSKEEAREAFEDNEK |
| *Aipysurus laevis F10* | MAPQLLLCLILTFLWSLPEAESNVFLKSKVANRFLQRTKRANSLFEEFKAGNIERECIEERCSKEEAREAFEDNEK |
| *Hydrophis major F10* | MAPQLLLCLIFTFLWSLPEAESNVFLKSKVANRFLQRTKRANSLFEEFKAGNIERECIEERCSKEEAREAFEDNEK |
| *Hydrophis elegans F10* | MAPQLLLCLIFTFLWSLPEAESNVFLKSKVANRFLQRTKRANSLFEEFKAGNIERECIEERCSKEEAREAFEDNEK |
| *Hydrophis curtus F10* | MAPQLLLCLIFTFLWSLPEAESNVFLKSKVANRFLQRTKRANSLFEEFKAGNIERECIEERCSKEEAREAFEDNEK |
| *Hydrophis ornatus F10* | MAPQLLLCLIFTFLWSLPEAESNVFLKSKVANRFLQRTKRANSLFEEFKAGNIERECIEERCSKEEAREAFEDNEK |
| *Bungarus candidus F10* | MAPQLLLCLILTFLWSLPEAESNVFLKSKVANRFLQRTKRANSLFEEIKPGNIERECREERCSKEEAREAFENNEE |
| *Natrix helvetica F10* | MAPQLLLCLILTFLWSLPNAESNVFLKGKVANRFLQRTKRANSIFEEFKAGNIERECIEERCSKEEAREAFEDQEK |

**77 — 152**

| | |
|---|---|
| *Pseudonaja textilis vF10a* | TETFWNVYVDGDQCSSNPCHYRGICKDGIGSYTCTCLSGYEGKNCERVLYKSCRVDNGNCWHFCKSVQNDIQCSCA |
| *Pseudonaja textilis vF10b* | TETFWNVYVDGDQCSSNPCHYRGICKDGIGSYTCTCLSGYEGKNCERVLYKSCRVDNGNCWHFCKHVQNDIQCSCA |
| *Oxyuranus microlepidotus vF10* | TETFWNVYVDGDQCSSNPCHYRGTCKDGIGSYTCTCLFGYEGKNCERVLYKSCRVDNGNCWHFCKPVQNDIQCSCA |
| *Notechis scutatus vF10* | TETFWNVYVDGDQCSSNPCHYRGTCKDGIGSYTCTCLPNYEGKNCEKVLYQSCRVDNGNCWHFCKRVQSETQCSCA |
| *Pseudonaja textilis F10* | TETFWNVYVDGDQCSSNPCHYGGTCKDGIGSYTCTCLSGYEGKNCEYVLYKSCRVDNGDCWHFCKPVQNGIQCSCA |
| *Oxyuranus microlepidotus F10* | TETFWNVYVDGDQCSSNPCHYGGTCKDGIGSYTCTCLSGYEGKNCEYVLYKSCRVDNGNCWHFCKPVQNGIQCSCA |
| *Notechis scutatus F10* | TETFWNVYVDGDQCSSNPCHYGGTCKDGIGSYTCTCLAGYEGKNCQYVLYQSCRVDNGNCWHFCKRVQSEIQCSCA |
| *Aipysurus laevis F10* | TETFWNVYVDGDQCSSNPCHYGGTCKDGIGSYTCTCLAGYEGKNCQYVLYKSCRVDNGNCWHFCKPVQNEIQCSCA |
| *Hydrophis major F10* | TETFWNVYVDGDQCSSNPCHYGGTCKDGIGSYTCTCLAGYEGKNCQYVLYKSCRVDNGNCWHFCKPVQNEIQCSCA |
| *Hydrophis elegans F10* | TETFWNVYVDGDQCSSNPCHYGGTCKDGIGSYTCTCLAGYEGKNCQYVLYKSCRVDNGNCWHFCKPVQNEIQCSCA |
| *Hydrophis curtus F10* | TETFWNVYVDGDQCSSNPCHYGGTCKDGIGSYTCTCLAGYEGKNCQYVLYKSCRVDNGNCWHFCKPVQNEIQCSCA |
| *Hydrophis ornatus F10* | TETFWNVYVDGDQCSSNPCHYGGTCKDGIGSYTCTCLAGYEGKNCQYVLYKSCRVDNGNCWHFCKPVQNEIQCSCA |
| *Bungarus candidus F10* | TEKFWNIYVDGDQCSSNPCHYGGTCKDGIGSYTCTCLAGYQGKNCEYIIYKSCRVDNGNCWHFCKLVQSEIQCSCA |
| *Natrix helvetica F10* | TENFWNVYVDGDQCLSNPCHYGATCKDGIGNFSCTCLAGYQGKTCESFILRSCRLDNGNCWHFCKSDQNKIHCSCA |

**153 — 228**

| | |
|---|---|
| *Pseudonaja textilis vF10a* | EGYLLGEDGHSCVAGGNFSCGRNIKTRNKR**EASLPDF-----------------------------VQSHNATLLK** |
| *Pseudonaja textilis vF10b* | EGYLLGEDGHSCVAGGNFSCGRNIKTRNKR**EANLPDF-----------------------------VQSQNATLLK** |
| *Oxyuranus microlepidotus vF10* | EGYLLGEDGHSCVAGGNFSCGRNIKTRNKR**EASLPDF-----------------------------VQSQNATLLK** |
| *Notechis scutatus vF10* | ESYRLGVDGHSCVAEGDFSCGRNIKARNKR**EASLPDF-----------------------------VQSQKATLLK** |
| *Pseudonaja textilis F10* | ESYLLGEDGHSCVAGGDFSCGRNIKTRNKR**EANLPDFQTDFSDDYDEIDENNFVETPTNFSGLVLTVQSQNATLLK** |
| *Oxyuranus microlepidotus F10* | ESYLLGEDGHSCVAGGDFSCGRNIKTRNKR**EANLPDFQTDFSDDYDEIDENNFVETPTNFSGLVPTVQSQNATLLK** |
| *Notechis scutatus F10* | ESYLLGDDGHSCVAGGDFSCGRNIKARNKR**EASLPDFQTDFSDDYDAIDENNFVETPTNFSGLVPTVQSQNATLLK** |
| *Aipysurus laevis F10* | ESYLLGDDGYSCVAEDDFSCGRNIKARNKR**EASLPDFQTDFSDDYDAIDENNFVETPTNFSGLVPTVQSQNASLLK** |
| *Hydrophis major F10* | ESYLLGDDGYSCVAEDDFSCGKNIKARNKR**EASLPDFQTDFSDDYDAIDENNFVETPTNFSGLVPTVQSQNANLLK** |
| *Hydrophis elegans F10* | ESYLLGDDGYSCVAEDDFSCGKNIKARNKR**EASLPDFQTDFSDDYDAIDENNFVETPTNFSGLVPTVQSQNANLLK** |
| *Hydrophis curtus F10* | ESYLLGDDGYSCVAEDDFSCGKNIKARNKR**EASLPDFQTDFSDDYDAIDENNFVETPTNFSGLVPTVQSQNANLLK** |
| *Hydrophis ornatus F10* | ESYLLGDDGYSCVAEDDFSCGKNIKARNKR**EASLPDFQTDFSDDYDAIDENNFVETPTNFSGLVPTVQSQNANLLK** |
| *Bungarus candidus F10* | ESYVLGDDEYSCVAEDDFSCGRNIKARNKR**EASLPDFQTEFSEDYDAIDENNFVETSTNFSSLVPTLQSRNATSLK** |
| *Natrix helvetica F10* | EGYILGDDGQSCVAGGDFSCGRNIKARNKR**EASLVDFQEDFSDDYDVIEEESFVETPTNFSSLVPIMQHRNATKPR** |

**229 — 304**

| | |
|---|---|
| *Pseudonaja textilis vF10a* | **KSDNPSPDIR**IVNGMDCKLGECPWQAALVDDKKGVFCGGTILSPIYVLTAAHCINETETISVVV-GEIDRSRAETG |
| *Pseudonaja textilis vF10b* | **KSDNPSPDIR**IVNGMDCKLGECPWQAALVDEKEGVFCGGTILSPIYVLTAAHCINETETISVVV-GEIDKSRIETG |
| *Oxyuranus microlepidotus vF10* | **KSDNPSPDIR**IVNGMDCKLGECPWQAVLVDEKEGVFCGGTILSPIYVLTAAHCINQTEKISVVV-GEIDKSRVETG |
| *Notechis scutatus vF10* | **KSDNPSPDIR**IVNGMDCKLGECPWQAVLINEKGEVFCGGTILSPIHVLTAAHCINQTKSVSVIV-GEIDISRKETR |
| *Pseudonaja textilis F10* | **KSDNPSPDIR**VVNGTDCKLGECPWQALLLNDEGDGFCGGTILSPIYVLTAAHCINQTKYITVVV-GEIDISSKKTG |
| *Oxyuranus microlepidotus F10* | **KSDNPSPDIR**VVNGTDCKLGECPWQALLINDQGDGFCGGTILSPIYVLTAAHCINQTKYIRVVV-GEIDISRKKTG |
| *Notechis scutatus F10* | **KSDNPSPDIR**VVNGTDCKLGECPWQALLINDQGDGFCGGTILSPIYVLTAAHCINQTKYIRVVV-EEIDISRKETR |
| *Aipysurus laevis F10* | **KSDNPSPDIR**VVNGTDCKLGECPWQALLINDQGDGFCGGTILSPIYVLTAAHCINQTKYIRVVV-GEIDISRKKTG |
| *Hydrophis major F10* | **KSDNPSPDVR**VVNGTDCKQGECPWQVLLINDRGDGFCGGTILSPIYVLTAAHCINQTKYIRVVVAGEIDISRKKTG |
| *Hydrophis elegans F10* | **KSDNPSPDIR**VVNGTDCKQGECPWQALLINDRGDGFCGGTILSPIYVLTAAHCINQTKYIRVVV-GEIDISRKKTG |
| *Hydrophis curtus F10* | **KSDNPSPDIR**VVNGTDCKQGECPWQALLINDRGDGFCGGTILSPIYVLTAAHCINQTKYIRVVV-GEIDISRKKTG |
| *Hydrophis ornatus F10* | **KSDNPSPDIR**VVNGTDCKQGECPWQALLINDRGDGFCGGTILSPIYVLTAAHCINQTKYIRVVV-GEIDISRKKTG |
| *Bungarus candidus F10* | **KSNNPSPEIR**VVNGTDCKLGECPWQALLLNDQGDGFCGGTILSSIYVLTAAHCINQTKHITVVV-GEVDISSKRNG |
| *Natrix helvetica F10* | **KSDNSNPEGR**VVNGTDCKPGECPWQALLINDQGDGFCGGTVLSPMYVLTAAHCINQTTRIKVVV-GEVDISSKKTG |

```
                                305                                                                       380
Pseudonaja textilis vF10a       PLLSVDKVYVHKKFVPPKKSQEFYEKFDLVSYDYDIAIIQMKTPIQFSENVVPACLPTADFANQVLMKQDFGIVSG
Pseudonaja textilis vF10b       PLLSVDKIYVHKKFVPPQKAY----KFDLAAYDYDIAIIQMKTPIQFSENVVPACLPTADFANQVLMKQDFGIVSG
Oxyuranus microlepidotus vF10   HLLSVDKIYVHKKFVPPKKGYKFYEKFDLVSYDYDIAIIQMKTPIQFSENVVPACLPTADFANQVLMKQDFGIISG
Notechis scutatus vF10          RLLSVDKIYVHTKFVPPNYYY-VHQNFDRVAYDYDIAIIRMKTPIQFSENVVPACLPTADFANEVLMKQDSGIVSG
Pseudonaja textilis F10         RLHSVDKIYVHQKFVP-------------ATYDYDIAIIQLKTPIQFSENVVPACLPTADFANQVLMKQNFGIVSG
Oxyuranus microlepidotus F10    RLLSVDKIYVHQKFVP-------------ATYDYDIAIIQLKTPIQFSENVVPACLPTADFANQVLMKQNFGIVSG
Notechis scutatus F10           RLLSVDKIYVHQKFVP-------------STYDYDIALIQMKTPIQFSENVVPACLPTADFANQVLMKQDFGIVSG
Aipysurus laevis F10            PL-SVDKIYVHKNFVP-------------TTYDYDIALIQLKTAIQFSENVVPACLPTADFANQVLMKQDFGIVSG
Hydrophis major F10             -LLSVDKIYVHKNFVP-------------TTYDYDIALIQLKTAIQFSENVVPACLPTADFANQVLMKQDFGIVSG
Hydrophis elegans F10           -LLSVDKIYVHKNFVP-------------TTYDYDIALIQLKTAIQFSENVVPACLPTADFANQVLMKQDFGIVSG
Hydrophis curtus F10            -LLSVDKIYVHKNFVP-------------TTYDYDIALIQLKTAIQFSENVVPACLPTADFANQVLMKQDFGIVSG
Hydrophis ornatus F10           -LLSVDKIYVHKNFVP-------------TTYDYDIALIQLKTAIQFSENVVPACLPTADFANQVLMKQDFGIVSG
Bungarus candidus F10           RLIAVDKIYVHQKFVP-------------ATYDYDIAIIQLKTPIQFSENVVPACLPTADFANQVLMKQNFGIVSG
Natrix helvetica F10            HLLAVNKIYVHQKFVL-------------ATYDYDIAIIQLKTPIRFSENVIPACLPTADFANQVLMKQDSGVISG

                                381                                                                       456
Pseudonaja textilis vF10a       FGGIFERGPNSKTLKVLKVPYVDRHTCMLSSNFPITPTMFCAGYDTLPQDACQGDSGGPHITAYRDTHFITGIVSW
Pseudonaja textilis vF10b       FGRIFEKGPKSKTLKVLKVPYVDRHTCMVSSETPITPNMFCAGYDTLPRDACQGDSGGPHTTVYRDTHFITGIVSS
Oxyuranus microlepidotus vF10   FGRIFEKGPKSNTLKVLKVPYVDRHTCMVSSESPITPTMFCAGYDTLPRDACQGDSGGPHITAYRDTHFITGIVSW
Notechis scutatus vF10          FGRIRFKEPTSNTLKVITVPYVDRHTCMLSSDFRITQNMFCAGYDTLPQDACQGDSGGPHITAYRDTHFITGIISW
Pseudonaja textilis F10         FGRTRERGKTSNTLKVVTLPYVDRHTCMLSSNFPITQNMFCAGYDTLPQDACQGDSGGPHITAYRDTHFITGIVSW
Oxyuranus microlepidotus F10    FGRTRERGQTSNTLKVVTLPYVDRHTCMLSSNFPITQNMFCAGYDTLPQDACQGDSGGPHITAYRDTHFITGIVSW
Notechis scutatus F10           FGRIRERGQTSNTLKVITLPYVDRHTCMLSSNFPITQNMFCAGYNTLPQDACQGDSGGPHITAYRDTHFITGIISW
Aipysurus laevis F10            FGRTRERGQTSKTLKVVTLPYVDRHTCMLSSNFPITQNMFCAGYNTLPQDACQGDSGGPHITAYRDTHFITGIISW
Hydrophis major F10             FGRTRERGKTSNTLKVVTLPYVDRHTCMLSSNFPITQNMFCAGYNTLPQDACQGDSGGPHITAYRDTHFITGIISW
Hydrophis elegans F10           FGRTRERGKTSNTLKVVTLPYVDRHTCMLSSNFPITQNMFCAGYNTLPQDACQGDSGGPHITAYRDTHFITGIISW
Hydrophis curtus F10            FGRTRERGKTSNTLKVVTLPYVDRHTCMLSSNFPITQNMFCAGYNTLPQDACQGDSGGPHITAYRDTHFITGIISW
Hydrophis ornatus F10           FGRTRERGKTSNTLKVVTLPYVDRHTCMLSSNFPITQNMFCAGYNTLPQDACQGDSGGPHITAYRDTHFITGIISW
Bungarus candidus F10           FGRTRERGRTSNTLKVVTLPYVDRHICKISSNFPITENMFCAGYNTLPQDACQGDSGGPHITAYRDTHFITGIISW
Natrix helvetica F10            FGRIKERGRTSNTLKMVTLPYVDRHVCKLSSNFPITENMFCAGYRTLAQDACQGDSGGPHVTAYSDTYFVTGIISW

                                457                                     497
Pseudonaja textilis vF10a       GEGCARKGRYGIYTKLSKFIPWIKRIMRQKLPSTESSTGRL
Pseudonaja textilis vF10b       GEGCARNGKYGIYTKLSKFIPWIKRIMRQKLPSTESSTGRL
Oxyuranus microlepidotus vF10   GEGCAKKGKYGIYTKVSKFILWIKRIMRQKLPSTESSTGRL
Notechis scutatus vF10          GEGCARKGKYGVYTKVSRFIPWIKKIMSLK-----------
Pseudonaja textilis F10         GEGCAQTGKYGVYTKVSKFILWIKRIIRQKQPSTESSTGQL
Oxyuranus microlepidotus F10    GEGCAQTGKYGVYTKVSKFILWIKRIMRQKLPSTESSTGRL
Notechis scutatus F10           GEGCAQTGKYGVYTKVSRFILWIKRIMRLKLPSTESSTGRL
Aipysurus laevis F10            GEGCAQTGKYGVYTKVSRFILWIKRIMRLKIPSTE------
Hydrophis major F10             GEGCAQTGKYGVYTKVSRFILWIKRIMRLKIPSTESSTGRL
Hydrophis elegans F10           GEGCAQTGKYGVYTKVSRFILWIKRIMRLKIPSTESSTGRL
Hydrophis curtus F10            GEGCAQTGKYGVYTKVSRFILWIKRIMRLKTPSTESSTGRL
Hydrophis ornatus F10           GEGCAQTGKYGVYTKVSRFILWIKRIMRLKIPSTESSTGRL
Bungarus candidus F10           GEGCAQPDKYGVYTKVSKFIPWIKRIMRQKLPGTDSSSGRL
Natrix helvetica F10            GEGCAQKDKYGVYTKVSKFIPWIKRIMRQKVPRTESGTARP
```

**Fig. S2. The venom factor X proteins have a truncated activation peptide region**. Amino acid sequence alignment of hypothetical protein translations of *F10* genes**.** Residues shown in grey encode the signal peptide and propeptide regions, and residues in bold indicate the activation peptide region, which is intact in proteins encoded by the *F10* genes but exhibits a 29-amino-acid deletion in proteins encoded by *vF10* genes.

```
Oxyuranus microlepidotus vF10    MAPQLLLCLILTFLWSLPEAESNVFLKSKVANRFLQRTKRANSLFEEFRSGNIERECIEERCSKEEAREVFEDDEK
Oxyuranus microlepidotus F10     MAPQLLLCLILTFLWSLPEAESNVFLKSKVANRFLQRTKRANSLFEEFKPGNIERECIEERCSKEEAREAFEDDEK
transcript/802606                MAPQLLLCLILTFLWSLPEAESNVFLKSKVANRFLQRTKRANSLFEEFKPGNIERECIEERCSKEEAREAFEDDEK

Oxyuranus microlepidotus vF10    TETFWNVYVDGDQCSSNPCHYRGTCKDGIGSYTCTCLFGYEGKNCERVLYKSCRVDNGNCWHFCKPVQNDIQCSCA
Oxyuranus microlepidotus F10     TETFWNVYVDGDQCSSNPCHYGGTCKDGIGSYTCTCLSGYEGKNCEYVLYKSCRVDNGNCWHFCKPVQNGIQCSCA
transcript/802606                TETFWNVYVDGDQCSSNPCHYGGTCKDGIGSYTCTCLSGYEGKNCEYVLYKSCRVDNGNCWHFCKPVQNGIQCSCA

Oxyuranus microlepidotus vF10    EGYLLGEDGHSCVAGGNFSCGRNIKTRNKREASLPDF-----------------------------VQSQNATLLK
Oxyuranus microlepidotus F10     ESYLLGEDGHSCVAGGDFSCGRNIKTRNKREANLPDFQTDFSDDYDEIDENNFVETPTNFSGLVPTVQSQNATLLK
transcript/802606                ESYLLGEDGHSCVAGGDFSCGRNIKTRNKREANLPDFQTDFSDDYDEIDENNFVETPTNFSGLVPTVQSQNATLLK

Oxyuranus microlepidotus vF10    KSDNPSPDIRIVNGMDCKLGECPWQAVLVDEKEGVFCGGTILSPIYVLTAAHCINQTEKISVVVGEIDKSRVETGH
Oxyuranus microlepidotus F10     KSDNPSPDIRVVNGTDCKLGECPWQALLINDQGDGFCGGTILSPIYVLTAAHCINQTKYIRVVVGEIDISRKKTGR
transcript/802606                KSDNPSPDIRVVNGTDCKLGECPWQALLINDQGDGFCGGTILSPIYVLTAAHCINQTKYIRVVVGEIDISRKKTGR

Oxyuranus microlepidotus vF10    LLSVDKIYVHKKFVPPKKGYKFYEKFDLVSYDYDIAIIQMKTPIQFSENVVPACLPTADFANQVLMKQDFGIISGF
Oxyuranus microlepidotus F10     LLSVDKIYVHQKFVP-------------ATYDYDIAIIQLKTPIQFSENVVPACLPTADFANQVLMKQNFGIVSGF
transcript/802606                LLSVDKIYVHQKFVP-------------ATYDYDIAIIQLKTPIQFSENVVPACLPTADFANQVLMKQNFGIVSGF

Oxyuranus microlepidotus vF10    GRIFEKGPKSNTLKVLKVPYVDRHTCMVSSESPITPTMFCAGYDTLPRDACQGDSGGPHITAYRDTHFITGIVSWG
Oxyuranus microlepidotus F10     GRTRERGQTSNTLKVVTLPYVDRHTCMLSSNFPITQNMFCAGYDTLPQDACQGDSGGPHITAYRDTHFITGIVSWG
transcript/802606                GRTRERGQTSNTLKVVTLPYVDRHTCMLSSNFPITQNMFCAGYDTLPQDACQGDSGGPHITAYRDTHFITGIVSWG

Oxyuranus microlepidotus vF10    EGCAKKGKYGIYTKVSKFILWIKRIMRQKLPSTESSTGRL
Oxyuranus microlepidotus F10     EGCAQTGKYGVYTKVSKFILWIKRIMRQKLPSTESSTGRL
transcript/802606                EGCAQTGKYGVYTKVSKFILWIKRIMRQKLPSTESSTGRL
```

**Fig. S3. The hemostatic *F10* gene is transcribed in inland taipan (*Oxyuranus microlepidotus*) venom gland tissue.** Protein sequence alignment comparing the hypothetical protein translations of the *F10* and *vF10* paralogs of *O. microlepidotus* with a representative full-length translated Iso-seq read corresponding to hemostatic factor X from venom gland tissue. The alignment provides qualitative evidence that full-length *F10* transcripts are expressed in venom gland tissue. Amino acid residues that differ between the *F10* and *vF10* sequences are shown in bold.

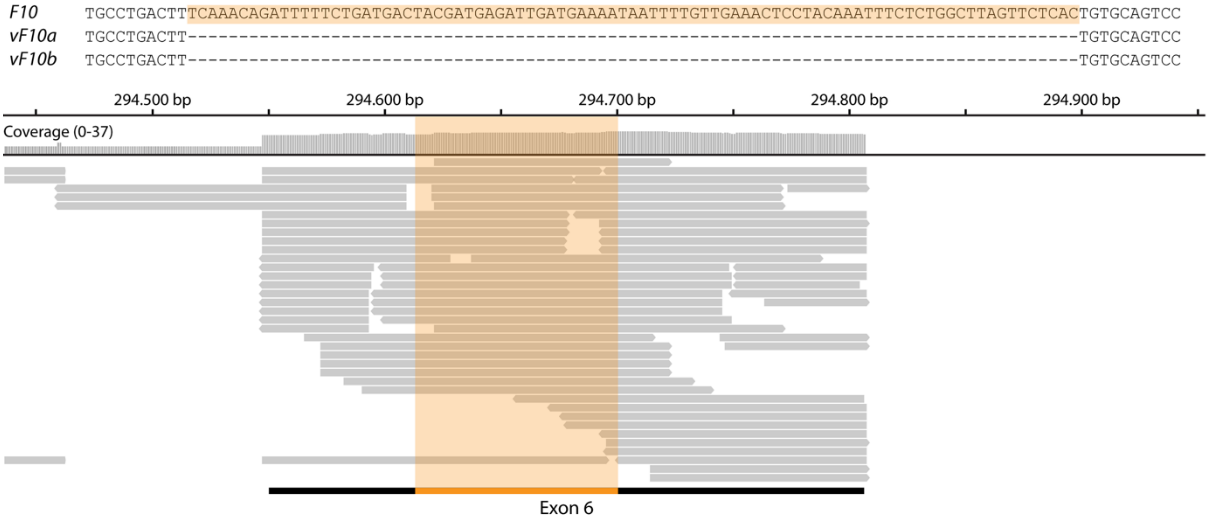

**Fig. S4. The *F10* gene is expressed in Eastern brown snake (*Pseudonaja textilis*) venom gland tissue.** One distinguishing feature between the *F10* and *vF10* paralogs is an 87-nucleotide deletion in exon 6 of the *vF10* gene. The corresponding nucleotide sequence in the *F10* gene (shown in orange in the sequence alignment) is absent in *vF10*. Taking advantage of this structural variant, we identified multiple short-read RNA-seq reads from *P. textilis* venom gland tissue that map to this *F10*-specific region (highlighted in orange), providing evidence that the hemostatic *F10* gene is also transcribed in the venom gland of *P. textilis*.

```
Pseudonaja textilis vF10a    MAPQLLLCLILTFLWSLPEAESNVFLKSKVANRFLQRTKRANSLVEEFKSGNIERECIEERCSKEEAREVFEDDEK
Pseudonaja textilis vF10b    MAPQLLLCLILTFLWSLPEAESNVFLKSKVANRFLQRTKRANSLVEEFKSGNIERECIEERCSKEEAREVFEDDEK
Pseudonaja textilis F10      MAPQLLLCLILTFLWSLPEAESNVFLKSKVANRFLQRTKRANSLFEEFKPGNIERECIEERCSKEEAREAFEDDEK

Pseudonaja textilis vF10a    TETFWNVYVDGDQCSSNPCHYRGICKDGIGSYTCTCLSGYEGKNCERVLYKSCRVDNGNCWHFCKSVQNDIQCSCA
Pseudonaja textilis vF10b    TETFWNVYVDGDQCSSNPCHYRGICKDGIGSYTCTCLSGYEGKNCERVLYKSCRVDNGNCWHFCKHVQNDIQCSCA
Pseudonaja textilis F10      TETFWNVYVDGDQCSSNPCHYGGTCKDGIGSYTCTCLSGYEGKNCEYVLYKSCRVDNGDCWHFCKPVQNGIQCSCA

Pseudonaja textilis vF10a    EGYLLGEDGHSCVAGGNFSCGRNIKTRNKREASLPDF-----------------------------VQSHNATLLK
Pseudonaja textilis vF10b    EGYLLGEDGHSCVAGGNFSCGRNIKTRNKREANLPDF-----------------------------VQSQNATLLK
Pseudonaja textilis F10      ESYLLGEDGHSCVAGGDFSCGRNIKTRNKREANLPDFQTDFSDDYDEIDENNFVETPTNFSGLVLTVQSQNATLLK

Pseudonaja textilis vF10a    KSDNPSPDIRIVNGMDCKLGECPWQAALVDDKKGVFCGGTILSPIYVLTAAHCINETETISVVVGEIDRSRAETGP
Pseudonaja textilis vF10b    KSDNPSPDIRIVNGMDCKLGECPWQAALVDEKEGVFCGGTILSPIYVLTAAHCINETETISVVVGEIDKSRIETGP
Pseudonaja textilis F10      KSDNPSPDIRVVNGTDCKLGECPWQALLLNDEGDGFCGGTILSPIYVLTAAHCINQTKYITVVVGEIDISSKKTGR

Pseudonaja textilis vF10a    LLSVDKVYVHKKFVPPKKSQEFYEKFDLVSYDYDIAIIQMKTPIQFSENVVPACLPTADFANQVLMKQDFGIVSGF
Pseudonaja textilis vF10b    LLSVDKIYVHKKFVPPQKAY----KFDLAAYDYDIAIIQMKTPIQFSENVVPACLPTADFANQVLMKQDFGIVSGF
Pseudonaja textilis F10      LHSVDKIYVHQKFVP-------------ATYDYDIAIIQLKTPIQFSENVVPACLPTADFANQVLMKQNFGIVSGF

Pseudonaja textilis vF10a    GGIFERGPNSKTLKVLKVPYVDRHTCMLSSNFPITPTMFCAGYDTLPQDACQGDSGGPHITAYRDTHFITGIVSWG
Pseudonaja textilis vF10b    GRIFEKGPKSKTLKVLKVPYVDRHTCMVSSETPITPNMFCAGYDTLPRDACQGDSGGPHTTVYRDTHFITGIVSSG
Pseudonaja textilis F10      GRTRERGKTSNTLKVVTLPYVDRHTCMLSSNFPITQNMFCAGYDTLPQDACQGDSGGPHITAYRDTHFITGIVSWG

Pseudonaja textilis vF10a    EGCARKGRYGIYTKLSKFIPWIKRIMRQKLPSTESSTGRL
Pseudonaja textilis vF10b    EGCARNGKYGIYTKLSKFIPWIKRIMRQKLPSTESSTGRL
Pseudonaja textilis F10      EGCAQTGKYGVYTKVSKFILWIKRIIRQKQPSTESSTGQL
```

**Fig S5. Coagulation factor X protein is detected in Eastern brown snake (*Pseudonaja textilis*) venom.** Amino acid sequence alignment of hypothetical *F10* gene translations with tryptic peptides shown in bold. Red residues indicate peptides unique to the venom paralogs (*vF10a* and/or *vF10b*), purple residues indicate peptides unique to the liver protein (*F10*), and black residues indicate peptides shared by all paralogs. Residues shown in grey correspond to the signal peptide and propeptide regions. Tryptic peptides corresponding to the hemostatic protein were detected in the venoms from three of seven individuals.

| | |
|---|---|
| *Oxyuranus microlepidotu F7b* | MISCHFGVFLCLLLLASPIARTSVFLRHKEASNILQRAR**R**ANSFLEELKA**G**SIERECLE**E**KCS**W**EEAREIFQDDLR |
| transcript/829350 | MISCHFGVFLCLLLLASPIARTSVFLRHKEASNILQRAR**R**ANSFLEELKA**G**SIERECLE**E**KCS**W**EEAREIFQDDLR |
| *Oxyuranus microlepidotu F7a* | MISCHFGVFLCLLLLASPIARTSVFLRHKEASNILQRAR**Q**ANSFLEELKA**E**SIERECLE**G**KCS**C**EEAREIFQDDLR |
| transcript/809936 | MISCHFGVFLCLLLLASPIARTSVFLRHKEASNILQRAR**Q**ANSFLEELKA**E**SIERECLE**G**KCS**C**EEAREIFQDDLR |
| *Oxyuranus microlepidotu F7b* | TKEFW**A**IYIDPDQCDSNPCQNGGTCIDRPQDYFCICPPEHEGRNCEIGPEIKLKCF**FK**NGNCEQFCVDSPTTLRQC |
| transcript/829350 | TKEFW**A**IYIDPDQCDSNPCQNGGTCIDRPQDYFCICPPEHEGRNCEIGPEIKLKCF**FK**NGNCEQFCVDSPTTLRQC |
| *Oxyuranus microlepidotu F7a* | TKEFW**E**IYIDPDQCDSNPCQNGGTCIDRPQDYFCICPPEHEGRNCEIGPEIKLKCF**LE**NGNCEQFCVDSPTTLRQC |
| transcript/809936 | TKEFW**E**IYIDPDQCDSNPCQNGGTCIDRPQDYFCICPPEHEGRNCEIGPEIKLKCF**LE**NGNCEQFCVDSPTTLRQC |
| *Oxyuranus microlepidotu F7b* | SCAEGYRLASDQVSCIAEVDYPCGKIPVLAKRPNQQGRIVGGYDCPPGECPWQALITENGNEKCGGVLVAPSWIIS |
| transcript/829350 | SCAEGYRLASDQVSCIAEVDYPCGKIPVLAKRPNQQGRIVGGYDCPPGECPWQALITENGNEKCGGVLVAPSWIIS |
| *Oxyuranus microlepidotu F7a* | SCAEGYRLASDQVSCIAEVDYPCGKIPVLAKRPNQQGRIVGGYDCPPGECPWQALITENGNEKCGGVLVAPSWIIS |
| transcript/809936 | SCAEGYRLASDQVSCIAEVDYPCGKIPVLAKRPNQQGRIVGGYDCPPGECPWQALITENGNEKCGGVLVAPSWIIS |
| *Oxyuranus microlepidotu F7b* | AAHCFSRVRKENLRIRLGEYHLNHRDEGEQERRVDELIIHEKYSFKTVDNDIALLRLSAPVNFSEYVVPICLPPPR |
| transcript/829350 | AAHCFSRVRKENLRIRLGEYHLNHRDEGEQERRVDELIIHEKYSFKTVDNDIALLRLSAPVNFSEYVVPICLPPPR |
| *Oxyuranus microlepidotu F7a* | AAHCFSRVRKENLRIRLGEYHLNHRDEGEQERRVDELIIHEKYSFKTVDNDIALLRLSAPVNFSEYVVPICLPPPR |
| transcript/809936 | AAHCFSRVRKENLRIRLGEYHLNHRDEGEQERRVDELIIHEKYSFKTVDNDIALLRLSAPVNFSEYVVPICLPPPR |
| *Oxyuranus microlepidotu F7b* | FAADILNYVEFSTVSGWGRLLEGGSTSALLMRVEVPKIHKKECVRHTKFNITDNMFCAGYLDGSKDSCEGDSGGPH |
| transcript/829350 | FAADILNYVEFSTVSGWGRLLEGGSTSALLMRVEVPKIHKKECVRHTKFNITDNMFCAGYLDGSKDSCEGDSGGPH |
| *Oxyuranus microlepidotu F7a* | FAADILNYVEFSTVSGWGRLLEGGSTSALLMRVEVPKIHKKECVRHTKFNITDNMFCAGYLDGSKDSCEGDSGGPH |
| transcript/809936 | FAADILNYVEFSTVSGWGRLLEGGSTSALLMRVEVPKIHKKECVRHTKFNITDNMFCAGYLDGSKDSCEGDSGGPH |
| *Oxyuranus microlepidotu F7b* | VTEYKNTWYLTGLVSWGKGCAAIGTYGVYTKVIKYHQWLNNHMDS |
| transcript/829350 | VTEYKNTWYLTGLVSWGKGCAAIGTYGVYTKVIKYHQWLNNHMDS |
| *Oxyuranus microlepidotu F7a* | VTEYKNTWYLTGLVSWGKGCAAIGTYGVYTKVIKYHQWLNNHMDS |
| transcript/809936 | VTEYKNTWYLTGLVSWGKGCAAIGTYGVYTKVIKYHQWLNNHMDS |

**Fig S6. Full-length *F7* transcripts are present in inland taipan (*Oxyuranus microlepidotus*) venom gland tissue.** Protein sequence alignment of the hypothetical protein translations encoded by the *F7a* and *F7b* paralogs of *O. microlepidotus*, together with representative full-length translated Iso-seq reads from venom gland tissue corresponding to each *F7* paralog. The alignment provides qualitative evidence that full-length *F7a and F7b* transcripts are expressed in venom gland tissue. Amino acid residues that differ between the *F7a* and *F7b* sequences are shown in bold.

| | |
|---|---|
| *Pseudonaja textilis F7a* | MISCHFGVFLCLLLLASPIARTSVFLRHKEASNILQRARRANSFLEELKAESIEQECLEQKCSCEEAREIFQDDLR |
| *Pseudonaja textilis F7b1* | MVSCHFGVFLCLLLLASPIARTSVFLRHKEASNILQRARRANSFLEELKAGSIERECLEENCSWEEAREIFQDDMR |
| *Pseudonaja textilis F7b2* | MISWHFGVFLCLLLLASPIARTSVFLRHKEASNILQRARRANSFLEELKAGSIERECLEEKCSWEEAREIFQDDLR |
| *Pseudonaja textilis F7a* | TKEFCEIYIDPDQCDSNPCQNGGTCIDRPQDYFCICPPEHEGRNCEIGPEIKLKCFLENGNCEQFCVDSPTTLRQC |
| *Pseudonaja textilis F7b1* | TKEFWAIYIDPDQCDSNPCQNGGTCIDRPQDYFCICPPEHEGRNCEIGPETKLKCFFKNGNCEQFCVDSPTTLR**QC** |
| *Pseudonaja textilis F7b2* | TKEFWAIYIDPDQCDSNPCQNGGTCIDRPQDYFCICPPEHEGRNCEIGPETKLKCFFKNGNCEQFCVDSPTTLR**QC** |
| *Pseudonaja textilis F7a* | SCAEGYKLASDQVSCIAEVDYPCGKIPVLAKRPNQQGRIVGGYDCPPGECPWQALITENGNEKCGGVLVAPSWIIS |
| *Pseudonaja textilis F7b1* | **SCAEGYR**LASDQVSCIAEVDYPCGKIPVLAKRPNQQGRIVGGYDCPPGECPWQALITENGNEKCGGVLVAPSWVIS |
| *Pseudonaja textilis F7b2* | **SCAEGYR**LASDQVSCIAEVDYPCGKIPVLAKRPNQQGRIVGGYDCPPGECPWQALITENGNEKCGGVLVAPSWVIS |
| *Pseudonaja textilis F7a* | AAHCFSHLRKENLRIR**LGEYHLNHRDEGEQER**RVDELIIHENYSFKTVDNDIALLHLSAPVNFSEYVVPICLPPPR |
| *Pseudonaja textilis F7b1* | AAHCFSHLRKENLRIR**LGEYHLNHRDEGEQER**RVDELIIHEKYSFKTVDNDIALLRLSAPVNFSEYVVPICLPPPR |
| *Pseudonaja textilis F7b2* | AAHCFSRVHKENLRIR**LGEYHLNHRDEGEQER**RVDELIIHEKYSFKTVDNDIALLRLSAPVNFSEYVVPICLPPPR |
| *Pseudonaja textilis F7a* | FAADILNYVEFSTVSGWGRLLEGGSTSALLMRVEVPKIHKKECVRHTKFNITDNMFCAGYLDGSK**DSCEGDSGGPH** |
| *Pseudonaja textilis F7b1* | FAADVLNYVEFSTVSGWGRLLEGGSTSALLMRVQVPKIHKKECVQHTKFNITDNMFCAGYLDGSK**DSCEGDSGGPH** |
| *Pseudonaja textilis F7b2* | FAADVLNYVEFSTVSGWGRLLEGGSTSALLMRVQVPKIHKKECVRHTKFNITDNMFCAGYLDGSK**DSCEGDSGGPH** |
| *Pseudonaja textilis F7a* | **VTEYK**NTWYLTGLVSWGKGCAAIGTYGVYTKVIKYHQWLNNHMDS |
| *Pseudonaja textilis F7b1* | **VTEYK**NTWYLTGLVSWGKGCAAIGTYGVYTKVIKYHQWLNNHMDS |
| *Pseudonaja textilis F7b2* | **VTEYK**NTWYLTGLVSWGKGCAAIGTYGVYTKVIKYHQWLNNHMDS |

**Fig S7. Coagulation factor VII proteins are detected in Eastern brown snake (*Pseudonaja textilis*) venom.** Amino acid sequence alignment of hypothetical F7 protein translations with tryptic peptides shown in bold. Residues shown in grey correspond to the signal peptide and propeptide regions. Tryptic peptides were detected in venom from three of seven individuals.

| | 1 85 |
|---|---|
| *P. textilis vF5* (short) | MGRYSVSPVPKCLLLMFLGWSGLKYYQVNAAQLREYHIAAQLEDWDYNPQPEELSRLSESDLTFKKIVYREYELDFKQEKPRDAL |
| *O. microlepidotus vF5* (short) | MGRYSVSPVPKCLLLMFLGWSGLKYYQVNAAQLREYRIAAQLEDWDYNPQPEELSRLSESELTFKKIVYREYELDFKQEKPRDEL |
| *P. textilis F5* (short) | MGRYSVSPVPKCLLLMFLGWSGLKYYQVNAAQLREYRIAAQLEDWDYNPQPEELSRLSESDLTFKKIVYREYELDFKQEKPRDAL |
| *O. microlepidotus F5* (short) | MGRYSVSPVPKCLLLMFLGWSGLKYYQVNAAQLREYRIAAQLEDWDYSPQPEELSRLSESELTFKKIVYREYELDFKQEKPRDEL |
| *P. textilis F5* | MGRYSVSPVPKCLLLMFLGWSGLKYYQVNAAQLREYRIAAQLEDWDYNPQPEELSRLSESDLTFKKIVYREYELDFKQEKPRDAL |
| *O. microlepidotus F5* | MGRYSVSPVPKCLLLMFLGWSGLKYYQVNAAQLREYRIAAQLEDWDYSPQPEELSRLSESELTFKKIVYREYELDFKQEKPRDEL |
| *N. scutatus F5* | MGRYSVSPVPKCLLLMFLGWSGLKYYQVNAAQLREYRIAAQLEDWDYNLQPEELSRLSESDRTFQKIVYREYELDFKQEKPRDEL |
| *B. candidus F5* | MGRYSVSPVSKCLLLMFLGWSGLKNYQVNAAQLREYRIAAQLEDWDYNLQPEGLSRLSESDRTFRKIVYREYELDFKQEKPRDKL |
| *N. helvetica F5* | MGRYSVSPIPKCLLLVFLGWSGLKYYQVNAAQLREYHIAAQVEDWDYNLQPGGLSRLSESDLAFKKIVYREYEVDFKQEKTRDEL |

| | 86 170 |
|---|---|
| *P. textilis vF5* (short) | SGLLGPTLRGEVGDSLIIYFKNFATQPVSIHPQSAVYNKWSEGSSYSDGTSDVERLDDAVPPGQSFKYVWNITAEIGPKKADPPC |
| *O. microlepidotus vF5* (short) | SGLLGPTLRGEVGDILIIYFKNFATQPVSIHPQSAVYNKWSEGSSYSDGTSDVERLDDAVPPGQSFKYVWNITAEIGPKKADPPC |
| *P. textilis F5* (short) | SGLLGPTLRGEVGDILIIYFKNFATQPVSIHPQSAVYNKWSEGSSYSDGTSDVERLDDAVPPGQSFKYVWNITAEIGPKKADPPC |
| *O. microlepidotus F5* (short) | SGLLGPTLRGEVGDILIIYFKNFATQPVSIHPQSAVYNKWSEGSSYSDGTSDVERLDDAVPPGQSFKYVWNITAEIGPKKADPPC |
| *P. textilis F5* | SGLLGPTLRGEVGDILIIYFKNFATQPVSIHPQSAVYNKWSEGSSYSDGTSDVERLDDAVPPGQSFKYVWNITAEIGPKKADPPC |
| *O. microlepidotus F5* | SGLLGPTLRGEVGDILIIYFKNFATQPVSIHPQSAVYNKWSEGSSYSDGTSDVERLDDAVPPGQSFKYVWNITAEIGPKKADPPC |
| *N. scutatus F5* | SGLLGPTLRGEVGDILIIYFKNFATKPVSIHPQSAVYNKWSEGSSYSDGTSDVERLDDAVPPGQSFKYVWNITAEIGPKKADPPC |
| *B. candidus F5* | SGLLGPTLRGEVGDILIIYFKNFATQPVSIHPQSAVYNKWSEGSSYSDGTSDMERLDNAVPPGQAFKYVWNITAEIGPKKADPPC |
| *N. helvetica F5* | SGLLGPTLRGEVGDVLVIYFKNFATQPVSIHPQSAVYNKWSEGASYSDGTSDVEKLDDAVPPDHSFKYVWNITAEIGPKETDPPC |

| | 171 255 |
|---|---|
| *P. textilis vF5* (short) | LTYAYYSHVNMVRDFNSGLIGALLICKEGSLNANGSQKFFNREYVLMFSVFDESKNWYRKPSLQYTINGFANGTLPDVQACAYDH |
| *O. microlepidotus vF5* (short) | LTYAYYSHVNMVRDFNSGLIGALLICKEGSLNANGAQKFFNREYVLMFSVFDESKNWYRKPSLQYTINGFANGTLPDVQACAYDH |
| *P. textilis F5* (short) | LTYAYYSHVNMVRDFNSGLIGALLICKEGSLNANGAQKFFNREYVLMFSVFDESKNWYRKPSLQYTINGFANGTLPDVQACAYDH |
| *O. microlepidotus F5* (short) | LTYSYYSHVNMVRDFNSGLIGALLICKEGSLNANGAQKFFNREYVLMFSVFDESKNWYRKPSLQYTINGFANGTLPDVQACAYDH |
| *P. textilis F5* | LTYAYYSHVNMVRDFNSGLIGALLICKEGSLNANGAQKFFNREYVLMFSVFDESKNWYRKPSLQYTINGFANGTLPDVQACAYDH |
| *O. microlepidotus F5* | LTYSYYSHVNMVRDFNSGLIGALLICKEGSLNANGAQKFFNREYVLMFSVFDESKNWYRKPSLQYTINGFANGTLPDVQACAYDH |
| *N. scutatus F5* | LTYAYYSHVNMVRDFNSGLIGALLICKEGSLNANGTQKLFNREYVLMFSVFDESKNWYRKPSLQYTINGFANGTLPDVQACAYDH |
| *B. candidus F5* | LTYAYYSHVNMVQDFNSGLIGALLICKKGSLTANGTQKLFNREYVLMFSVFDESKNWYKKPSLQYTINGFANGTLPDIQACAYDR |
| *N. helvetica F5* | LTYAYYSHVNMVRDFNSGLIGALLICKEGSLNADGTQKLFNREYVLMFAVFDESKNWYKKPSLQYTINGFANGTLPDIQACAYDR |

| | 256 340 |
|---|---|
| *P. textilis vF5* (short) | ISWHLIGMSSSPEIFSVHFNGQTLEQNHYKVSTINLVGGASVTADMSVSRTGKWLISSLVAKHLQAGMYGYLNIKDCGNPDTLTR |
| *O. microlepidotus vF5* (short) | ISWHLIGMSSSPEIFSVHFNGQTLEQNHYKVSTINLVGGASVTANMSVSRTGKWLISSLVAKHLQAGMYGYLNIKDCGHPNTLTR |
| *P. textilis F5* (short) | ISWHLIGMSSSPEIFSVHFNGQTLEQNHYKVSTINLVGGASVTANMSVSRTGKWLISSLVAKHLQAGMYGYLNIKDCGNPDTLTR |
| *O. microlepidotus F5* (short) | ISWHLIGMSSSPEIFSVHFNGQTLEQNHYKVSTINLVGGASVTANMSVSRTGKWLISSLVAKHLQAGMYGYLNIKDCGNPDTLTR |
| *P. textilis F5* | ISWHLIGMSSSPEIFSVHFNGQTLEQNHYKVSTINLVGGASVTANMSVSRTGKWLISSLVAKHLQAGMYGYLNIKDCGNPDTLTR |
| *O. microlepidotus F5* | ISWHLIGMSSSPEIFSVHFNGQTLEQNHYKVSTINLVGGASVTANMSVSRTGKWLISSLVAKHLQAGMYGYLNIKDCGNPDTLTR |
| *N. scutatus F5* | ISWHLIGMSSSPEIFSVHFNGQTLEQNHYKVSTINLIGGASVTAKMSVSRTGKWLISSLVAKHLQAGMYGYLNIKDCGNPDTLTR |
| *B. candidus F5* | ISWHLIGMSSSPEIFSVHFNGQTLEQNNYKVSTINLVGGASVTANMSVSRTGKWLISSLVAKHLQAGMYGYLNIKDCGKSDTLTR |
| *N. helvetica F5* | ISWHLIGMSSSPEIFSVHFNGQTLEENHYKVSTINLVGGASVTANMSVSMTGKWLISSLVAKHLQAGMYGYLNIKDCGKPDTLTK |

| | 341 425 |
|---|---|
| *P. textilis vF5* (short) | KLSFRELMKIKNWEYFIAAEEITWDYAPEIPSSVDRRYKAQYLDNFSNFIGKKYKKAVFRQYEDGNFTKPTYAIWPKERGILGPV |
| *O. microlepidotus vF5* (short) | KLSFRELRRIMNWEYFIAAEEITWDYAPEIPSSVDRRYKAQYLDNFSNFIGKKYKKAVFRQYEDGNFTKPTYAIWPKERGILGPV |
| *P. textilis F5* (short) | KLSFRELRRIMNWEYFIAAEEITWDYAPEIPSSVDRRYKAQYLDNFSNFIGKKYKKAVFRQYKDSNFTKPTYAIWPKERGILGPV |
| *O. microlepidotus F5* (short) | KLSFRELRRIMNWEYFIAAEEITWDYAPEIPSSVDRRYKAQYLDNFSNFIGKKYKKAVFRQYKDGNFTKPTYAIWPKERGILGPV |
| *P. textilis F5* | KLSFRELRRIMNWEYFIAAEEITWDYAPEIPSSVDRRYKAQYLDNFSNFIGKKYKKAVFRQYKDSNFTKPTYAIWPKERGILGPV |
| *O. microlepidotus F5* | KLSFRELRRIMNWEYFIAAEEITWDYAPEIPSSVDRRYKAQYLDNFSNFIGKKYKKAVFRQYKDGNFTKPTYAIWPKERGILGPV |
| *N. scutatus F5* | KLSFRELKMIKNWEYFIAAEEITWDYAPEIPRSVDRRYKAQYLDNFSNFIGKKYKKAVFRQYKDSSFSKPTYAIWPKERGILGPV |
| *B. candidus F5* | KLSFKELRMIKNWEYFIAAEEITWDYAPEIPSTVDRRYKAQYLDNFSNFIGKKYKKAVFRQYKDSSFTKPTYAIWPKERGILGPV |
| *N. helvetica F5* | RLTFKELRMIKNWDYFIAAEEITWDYAPEIPSSVDRRYKAQYLDNFSNFIGKKYKKAVFRQYKDSSFSKPTYATWPKERGILGPV |

| | 426 510 |
|---|---|
| *P. textilis vF5* (short) | IKAKVRDTVTIVFKNLASRPYSIYVHGVSVSKDAEGAIYPSDPKENITHGKAVEPGQVYTYKWTVLDTDEPTVKDSECITKLYHS |
| *O. microlepidotus vF5* (short) | IKAKVRDTVTIVFKNLASRPYSIYVHGVSVSKDAEGAIYPSDPKENITHGKAVEPGQVYTYKWTVLDTDEPTVKDSECITKLYHS |
| *P. textilis F5* (short) | IRAKVRDTISIVFKNLASRPYSIYVHGVSVSKDAEGAIYPSDPKENITHGKAVEPGQVYTYKWTVLDTDEPTVKDSECITKLYHS |
| *O. microlepidotus F5* (short) | IRAKVRDTISIVFKNLASRPYSIYVHGVSVSKDAEGAIYPSDPKENITHGKAVEPGQVYTYKWTVLDTDEPTVKDSECITKLYHS |
| *P. textilis F5* | IRAKVRDTISIVFKNLASRPYSIYVHGVSVSKDAEGAIYPSDPKENITHGKAVEPGQVYTYKWTVLDTDEPTVKDSECITKLYHS |
| *O. microlepidotus F5* | IRAKVRDTISIVFKNLASRPYSIYVHGVSVSKDAEGAIYPSDPKENITHGKAVEPGQVYTYKWTVLDTDEPTVKDSECITKLYHS |
| *N. scutatus F5* | IRAEVRDTISIVFKNLASRPYSIYVHGVSVSKDAEGAIYPSDPKENITHGKAVEPGQVYTYKWTVLDTDEPTVKDSQCITKLYHS |
| *B. candidus F5* | IRAEVRDTISIVFKNLASRPYSIYVHGVSVSKDAEGAIYPSDPKENITHGKAVEPGHIYTYKWTVLDTDEPTVKDSQCITKLYHS |
| *N. helvetica F5* | IRAGVRDTISIVFKNLASRPYSIYVHGVSVSKDAEGAVYPSDSKENITQGKAVEPGDVYTYKWTVLDTDEPTAKDSQCITKLYHS |

| | 511 595 |
|---|---|
| *P. textilis vF5* (short) | AVDMTRDIASGLIGPLLVCKHKALSVKGVQNKADVEQHAVFAVFDENKSWYLEDNIKKYCSNPSAVKKDDPKFYKSNVMYTLNGY |
| *O. microlepidotus vF5* (short) | AVDMTRDIASGLIGPLLVCKLKALSVKGVQNKADVEQHAVFAVFDENKSWYLEDNIKKYCSNPSSVKKDDPKFYKSNVMYTLNGY |
| *P. textilis F5* (short) | AVDMTRDIASGLIGPLLVCKHKALSVKGVQNKADVEQHAVFAVFDENKSWYLEDNIKKYCSNPSTVKKDDPKFYKSNVMYTLNGY |
| *O. microlepidotus F5* (short) | AVDMTRDIASGLIGPLLVCKRKALSVKGVQNKADVEQHAVFAVFDENKSWYLEDNIKKYCSNPSSVKKDDPKFYKSNVMYTLNGY |
| *P. textilis F5* | AVDMTRDIASGLIGPLLVCKHKALSVKGVQNKADVEQHAVFAVFDENKSWYLEDNIKKYCSNPSTVKKDDPKFYKSNVMYTLNGY |
| *O. microlepidotus F5* | AVDMTRDIASGLIGPLLVCKRKALSVKGVQNKADVEQHAVFAVFDENKSWYLEDNIKKYCSNPSSVKKDDPKFYKSNVMYTLNGY |
| *N. scutatus F5* | AVDMTRDIASGLIGPLLVCKRKALNSKGVQDKADVEQHAVFAVFDENKSWYLEDNIKKYCSNSSTVKKDDPKFYKSNVMYTLNGY |
| *B. candidus F5* | AVDMTRDIASGLIGPLLVCKRKALDSKGVQKKADVEQHAVFAVFDENKSWYLEDNIKKYCSNSSTVKKDDPKFYKSNVMYTLNGY |
| *N. helvetica F5* | AVDMTRDIASGLIGPLLVCKRKALDNRGVQKKADVEQHAIFAVFDENKSWYLDDNIKKYSSNFSTVKKDDPKFYKSNVMYTLNGY |

```
                                  596                                                                                680
P. textilis vF5 (short)           ASDRTEVLRFHQSEVVQWHLTSVGTVDEIVPVHLSGHTFLSKGKHQDILNLFPMSGESATVTMDNLGTWLLSSWGSCEMSNGMRL
O. microlepidotus vF5 (short)     ASDRTEVLGFHQSEVVQWHLTSVGTVDEIVPVHLSGHTFLSKGKHQDILNLFPMSGESATVTMDNLGTWLLSSWGSCEMSNGMRL
P. textilis F5 (short)            ASDRTEVLGFHQSEVVEWHLTSVGTVDEIVPVHLSGHTFLSKGKHQDILNLFPMSGESATVTMDNLGTWLLSSWGSCEMSNGMRL
O. microlepidotus F5 (short)      ASDRTEVLGFHQSEVVEWHLTSVGTVDEIVPVHLSGHTFLSKGKHQDILNLFPMSGESATVTMDNLGTWLLSSWGSCEMSNGMRL
P. textilis F5                    ASDRTEVLGFHQSEVVEWHLTSVGTVDEIVPVHLSGHTFLSKGKHQDILNLFPMSGESATVTMDNLGTWLLSSWGSCEMSNGMRL
O. microlepidotus F5              ASDRTEVLGFHQSEVVEWHLTSVGTVDEIVPVHLSGHTFLSKGKHQDILNLFPMSGESATVTMDNLGTWLLSSWGSCEMSNGMRL
N. scutatus F5                    ASDRTDVLGFRQSEIVEWHLTSVGTVDEIVPVHLSGHTFLSKGKHQDILNLFPMSGESATVTMDNLGTWLLSSWGSREMSNGMRL
B. candidus F5                    ASDRTDVLGFRQSEIVEWHLTSVGTVDEIVPVHLSGHTFLSKGKHQDILNLFPMRGESATVTMDNLGTWLLSSWGSREMSNGMRL
N. helvetica F5                   ASDRTEVLGFHQSEIVEWHLTSVGTMDEIIPVHLSGHTFLSKGKHQDILNLFPMSGESATVTMDNLGTWLLSSWGSREMSNGMRL

                                  681                                                                                765
P. textilis vF5 (short)           RFLDANYDDEDEGNEEEEEDDGDIFADI--FIPSEVVKK---KEEVPVNFVPDPESDALAKELGLIDDEGNPIIQPRREQTEDDE
O. microlepidotus vF5 (short)     RFLDANYDDEDEGNEEEEEDDGDIFADI--FSPPEVVKK---KEEVPVNFVPDPESDALAKELGLLDDEDNP-EQSRSEQTEDDE
P. textilis F5 (short)            RFLDANYDDEDEGNEEEEEDDGDIFADI--FIPPEVVKK---KEKDPVNFVSDPESDKIAKELGLLDDEDNQ-EESHNVQTEDDE
O. microlepidotus F5 (short)      RFLDANYDDEDEGNEEEEEDDGDIFADI--FSPPEVVKK---KEKDPVNFVSDPESDKIAKELGLLDDEDNQ-EESHNEQTEDDE
P. textilis F5                    RFLDANYDDEDEGNEEEEEDDGDIFADI--FIPPEVVKK---KEKDPVNFVSDPESDKIAKELGLLDDEDNQ-EESHNVQTEDDE
O. microlepidotus F5              RFLDANYDDEDEGNEEEEEDDGDIFADI--FSPPEVVKK---KEKDPVNFVSDPESDKIAKELGLLDDEDNQ-EESHNEQTEDDE
N. scutatus F5                    RFLDAKHDDEDEGNEKEEEHYGGIVANIISGGPSEVVKK---KEKDPRNVVSDSEMDKIAKELGLLDDEDNQ-KESSNEQTEDDE
B. candidus F5                    RFLDANYDDEDEGNEKEEEEDGDIVANIVSFSSPDVIKE---KRKDSKSVVSDPETDKIAKELGLLDDEDNQ-KESSNEQIEDDE
N. helvetica F5                   RFLDANYDDEDDRDEK-EEDYDDIVANIVSFDPSEVTEKKEEKEKDLRKVVSDPESDEIAKQLGLLDDEDST-YKSSNEQAEDDE

                                  766                                                                                850
P. textilis vF5 (short)           EQLMKASMLGLRSFKGSVA--EEELKHTALALEEDAHAS----------------------------------------------
O. microlepidotus vF5 (short)     EQLMIASMLGLRSFKGSVA--EEELKHTALALEEDAHAS----------------------------------------------
P. textilis F5 (short)            EQLMIATMLGFRSFKGSVA--EEELNLTALALEEDAHAS----------------------------------------------
O. microlepidotus F5 (short)      EQLMIATMLGLRSFKGSVA--EEELNLTALALEEDAHAS----------------------------------------------
P. textilis F5                    EQLMIATMLGFRSFKGSVA--EEELNLTALALEEDAHASGELSLNTHLVLINSTDAMRGNEELGTQNNSYVNASETIYANYTSST
O. microlepidotus F5              EQLMIATMLGLRSFKGSVA--EEELNLTALALEEDAHASGELSLNTHLVLINSTDAMRGNEELGTRNNSYVNASETIYANYTSST
N. scutatus F5                    EQLMIASMLGLRSFNGSVA--EEELNLTALALEEDAHTSGELSLNTHLVLINSTDAMRGNEELGTQNNSYVNASETIYANYTSST
B. candidus F5                    EQLMIASMLGLRTFNGSVA-EEEELNLTALALQEDAHTSGDVSLNTHLVLINSTDAMHGNEELGTQNNSYVTASETIYANYTSST
N. helvetica F5                   EQLMIASMLGLRSFNGSVAEEEEEELNLTALAIQEDADASGDVSLNDYIALNNSSDAMLGNGELGTQNNSYVTASETIYANCTSLA

                                  851                                                                                935
P. textilis vF5 (short)           -------------------------------------------------------------------------------------
O. microlepidotus vF5 (short)     -------------------------------------------------------------------------------------
P. textilis F5 (short)            -------------------------------------------------------------------------------------
O. microlepidotus F5 (short)      -------------------------------------------------------------------------------------
P. textilis F5                    GEILISNITAMIGILLSKRSQNEENITQISDSYKEMLAAEYLKAGRGLQSKDIEDLMSPEIIPSLKEQEENTFEKDIWDKKGERQ
O. microlepidotus F5              GEILISNITAMIGILLSKKSQNEENITQISDSYKQMLAAEYLKAGRGLQSQDIEDLMSPEIIPSLKEQEENTFEKDIWDKKGERQ
N. scutatus F5                    GEILISNITAMIGILLSKKSQNEENITQISDSYKQMLAAEYLKAGRGLQSKDIEDLMSPEIIPSLKEQEENTFEKDIWDKQGERQ
B. candidus F5                    GEILISNITEMIGILLSKKSQNEENITQILDSYKQMLAAEYLKAGRGLQSKDIEDLISLEIIPSLKKQEENTFEKDIWDKKGERQ
N. helvetica F5                   GEILTSNITAVIDILLSKISQNEENITQISDSCKQFLAAEYLKTDRGPQSKDIEDLMSLEIISSMKEQEENTFEKYIWDKKRETQ

                                  936                                                                               1020
P. textilis vF5 (short)           -------------------------------------------------------------------------------------
O. microlepidotus vF5 (short)     -------------------------------------------------------------------------------------
P. textilis F5 (short)            -------------------------------------------------------------------------------------
O. microlepidotus F5 (short)      -------------------------------------------------------------------------------------
P. textilis F5                    ILTEATLHALKEMHALFFYVQQKRNLSAFNITSEGMKFLPSTENDSTIDNSKANYFSYDYDYDYKEEEATAAANVEFSKVIINTI
O. microlepidotus F5              ILTEATLHALKEMHALFFYVQQKRNLSAFNITSEDMKFLPSTENDSTIDNSKANYFSYDYDYDYKEEEATAAANVEFSKVIINTI
N. scutatus F5                    ILTEATLHALKEMHALFFYVQQKRNLSAFNITSEDMKFLSSTENDSTTDNSKANYFSYDYDYDYKEEEATAAANVEFSKVITNTI
B. candidus F5                    VLSEATLHALKEMYALFFYVQQKRNISAFNITNEDMKFLSSTENDSTLDNSKANYFSYDYDYDYKGGEATAAANVEFPKVITNAT
N. helvetica F5                   VLTETTLYALKEMYALFFHVQQKRNLSAFNITSEDMKFLPSTENDSTLDDSKANYFPYEYDYDYK-EEATAAANVEFSKVIMNIT

                                  1021                                                                              1105
P. textilis vF5 (short)           -------------------------------------------------------------------------------------
O. microlepidotus vF5 (short)     -------------------------------------------------------------------------------------
P. textilis F5 (short)            -------------------------------------------------------------------------------------
O. microlepidotus F5 (short)      -------------------------------------------------------------------------------------
P. textilis F5                    SDDKKTVENNRSEVNLVSVPSIMPQPENLTSDATLDNFVSTTISKTTKTWNSSSHQRQKCLPKNTDMEW----KKENYDTLADIP
O. microlepidotus F5              SDDKKTVENNRSEVNLVSVPSIMPQPENLTSDATLDNFVSTTISKTTKAWNSSSHQRQKCLPKNTDMEW----KKENYDTLADIP
N. scutatus F5                    SDDKKTVENNGSEVNLASVPSIMPQPENLTSDATLDNFVSTTILKTTKTWNSSSHQRQKCLPKNTDMEW----KKENYDTLADIT
B. candidus F5                    SADKKTVENNGSELNLVSVPSIMPQPENLTSDATLDNFVSTTISKTTKTWNSSSHQKQKCLPKNTGMGW----KKENYDTLADIT
N. helvetica F5                   SADKETVENNGSEPNLVSVPSIMPQSENLTSNATLDNFVSTTISNSTKTWKRSSPQRQKCFPKNTDKEWNVVDKKENYDILADIT

                                  1106                                                                              1190
P. textilis vF5 (short)           -------------------------------------------------------------------------------------
O. microlepidotus vF5 (short)     -------------------------------------------------------------------------------------
P. textilis F5 (short)            -------------------------------------------------------------------------------------
O. microlepidotus F5 (short)      -------------------------------------------------------------------------------------
P. textilis F5                    NDVDNKFQNSSRNASFYQENAMFPGKRKEKTDNSNGQLELGQPGITNKHDSQKDENCTINISQSFIKIQRKKKKPKVLTPRTSKH
O. microlepidotus F5              NDVDNKFQNSSRNASFYQENAMFPGKRKEKTDNSNGQLELGQPGITNKHDSQKDENCTINISQSFIKIQRKKKKPKVLTPRTSKH
N. scutatus F5                    NDVDNKFQNSSRNASFYQENAMFPGKRKEKTDNSNGQLELGQPGITNKHDSQKDENCTINISQSFIKIQRKKKKPKVLTPRASKH
B. candidus F5                    NDGDNKFQNSSRNASFYQENAMFPDKRKEKTDNSNGQLELGQPGITNKHDAQKDENCTINISQSFIKIQRKKKKPKVLTPRTSKP
N. helvetica F5                   NDVDNKFQNSSRNASFYQENALFPGKRKEKTDNSNGQLELGQPGINNKNDAQKDENCTINISQPFVKIQRKKKKPNVLAPRTFKP
```

| | 1191 … 1275 |
|---|---|
| *P. textilis vF5* (short) | **------------------------------------------------------------------------------DPRIDSN** |
| *O. microlepidotus vF5* (short) | **------------------------------------------------------------------------------DPRIDSN** |
| *P. textilis F5* (short) | **------------------------------------------------------------------------------DPRIDSN** |
| *O. microlepidotus F5* (short) | **------------------------------------------------------------------------------DPRIDSN** |
| *P. textilis F5* | **LELPRNLNNSQLSNKPNHTKLLNEAKLTRKQDKLIVTIGLPVEDGDYQEYDIDNTDNDRSSSGSFEYELMYYNNPYTTDPRIDSN** |
| *O. microlepidotus F5* | **LELPRNLNNSQLSNKPNHTKLLNEAKLTRKQDKFIVTIGLPVEDGDYQEYEIDNTDNDKSSSGSFEYELMYYNNPYTTDPRIDSN** |
| *N. scutatus F5* | **LELPRNLNHSQLSNKPNHTKLLNEAKLTRKQDK-LVTIGLPVEDGDYQEYEIDNTDNDKSSSGSFEYELMYYDNPYTTDPRIDSN** |
| *B. candidus F5* | **LDLPRNLNNSQFSNKPNHTKLLNEAKLTRKQDKPIVTIGLPVEDGDYQEYEIDNIDNDQSSSGSFEYELMFYDNPYTTDSRLDST** |
| *N. helvetica F5* | **LELPANLNNSQLSNKLNHTKLLNEAKLTQKQDKPIITIGLPVEDGGYQEYEIDNTDNDESSSGSFEYELMYYDNPYTTDSRLDSN** |

| | 1276 … 1360 |
|---|---|
| *P. textilis vF5* (short) | **SARNPDDIAGRYLR**TINRGNKRRYYIAAEEVLWDYSPIGKSQVRSRAAKTTFKKAIFRSYLDDTFQTPSTGGEYEKHLGILGPII |
| *O. microlepidotus vF5* (short) | **SARNSDDIAGRYLR**TINRRNKRRYYIAAEEVLWDYSPIGKSQVRSLPAKTTFKKAIFRSYLDDTFQTPSTGGEYEKHLGILGPII |
| *P. textilis F5* (short) | **SARNPDDIAGRYLR**TINRGNKRRYYIAAEEVLWDYSPIGKSQVRSRAAKTTFKKAIFRSYLDDTFQTPSTGGEYEKHLGILGPII |
| *O. microlepidotus F5* (short) | **SARNPDDIAGRYLR**TINRGNKRRYYIAAEEVLWDYSPIGKSQVRSRAAKTTFKKAIFRSYLDDTFQTPSTGGEYEKHLGILGPII |
| *P. textilis F5* | **SARNPDDIAGRYLR**TINRGNKRRYYIAAEEVLWDYSPIGKSQVRSRAAKTTFKKAIFRSYLDDTFQTPSTGGEYEKHLGILGPII |
| *O. microlepidotus F5* | **SARNPDDIAGRYLR**TINRGNKRRYYIAAEEVLWDYSPIGKSQVRSRAAKTTFKKAIFRSYLDDTFQTPSTGGEYEKHLGILGPII |
| *N. scutatus F5* | **SARNPDDIARHYLR**TTNSGNKRMYYIAAEEVLWDYSPIRKSQVRSHPAKTTFKKAIFRSYLDDTFQTPSTGGEYEKHLGILGPII |
| *B. candidus F5* | **SARNPDDIAGRYMR**TINRGNIRKYYIAAEEVLWDYSPIRKSQVRSRPVKTTFKKAIFRSYLDDTFQTPSTGGEYEKHLGILGPII |
| *N. helvetica F5* | **IARNPDDIAGRYLR**TINRGNIRRYYIAAEEIVWDYSPFRKSTVRSRPLRTTFKKAIFRSYLDDTFQTPSPVGEYEKHLGILGPII |

| | 1361 … 1445 |
|---|---|
| *P. textilis vF5* (short) | RAEVDDVIEIQFRNLASRPYSLHAHGLLYEKSSEGRSYDDKSPELFKKDDAIMPNGTYTYVWQVPPRSGPTDNTEKCKSWAYYSG |
| *O. microlepidotus vF5* (short) | RAEVDDVIEVQFRNLASRPYSLHAHGLLYEKSSEGRSYDDNSPELFKKDDAIMPNGTYTYVWQVPPRSGPTDNTEKCKSWAYYSG |
| *P. textilis F5* (short) | RAEVDDVIEVQFRNLASRPYSLHAHGLLYEKSSEGRSYDDKSPELFKKDDAIMPNGTYTYVWQVPPRSGPTDNTEKCKSWAYYSG |
| *O. microlepidotus F5* (short) | RAEVDDVIEVQFRNLASRPYSLHAHGLLYEKSSEGRSYDDKSPELFKKDDAIMPNGTYTYVWQVPPRSGPTDNTEKCKSWAYYSG |
| *P. textilis F5* | RAEVDDVIEVQFRNLASRPYSLHAHGLLYEKSSEGRSYDDKSPELFKKDDAIMPNGTYTYVWQVPPRSGPTDNTEKCKSWAYYSG |
| *O. microlepidotus F5* | RAEVDDVIEVQFRNLASRPYSLHAHGLLYEKSSEGRSYDDKSPELFKKDDAIMPNGTYTYVWQVPPRSGPTDNTEKCKSWAYYSG |
| *N. scutatus F5* | RAEVDDVIEVQFKNLASRPYSLHAHGLLYDKSSEGRSYDDQSPELFKKDEAIMPNGTYTYVWHVPQRSGPTDNTEKCKSWAYYSG |
| *B. candidus F5* | RAEVDDVIQVQFRNLASRPYSLHAHGLLYEKSSEGRSYDDQSPEFFKMDDAIMPNGTYTYVWQVTKRSGPTDNTEKCKSWAYYSG |
| *N. helvetica F5* | RAEVDDVIQVQFRNLASRPYSLHAHGLLYEKSSEGRSYDDHSPEFFKKDDAIMPNSTYTYVWQVTKRAGPTDDTEKCKTWAYYSG |

| | 1446 … 1530 |
|---|---|
| *P. textilis vF5* (short) | VNPEKDIHSGLIGPILICQKGMIDKYNRTIDIREFVLFFMVFDEEKSWYFPKSDKSTCEEKLIGVQSLHTFPAINGIPYQLQGLT |
| *O. microlepidotus vF5* (short) | VNPEKDIHSGLIGPILICQKGMIDKYNRTIDIREFVLFFMVFDEEKSWYFPKSDKSTCEEKLIGVQSHHTFPAINGIPYQLQGLM |
| *P. textilis F5* (short) | VNPEKDIHSGLIGPILICQKGMIDKYNRTIDIREFVLFFMVFDEEKSWYFPKSDKSTRAEKLIGVQSRHTFPAINGIPYQLQGLT |
| *O. microlepidotus F5* (short) | VNPEKDIHSGLIGPILICQKGMIDKYNRTIDIREFVLFFMVFDEEKSWYFPKSDKSTRAEKLIGVQSRHTFPAINGIPYQLQGLM |
| *P. textilis F5* | VNPEKDIHSGLIGPILICQKGMIDKYNRTIDIREFVLFFMVFDEEKSWYFPKSDKSTRAEKLIGVQSRHTFPAINGIPYQLQGLT |
| *O. microlepidotus F5* | VNPEKDIHSGLIGPILICQKGMIDKYNRTIDIREFVLFFMVFDEEKSWYFPKSDKSTRAEKLIGVQSRHTFPAINGIPYQLQGLM |
| *N. scutatus F5* | VNPEKDIHSGLIGPILICQKGMIDKYNKTIDIREFVLFFMVFDEERSWYFAKSDKRTRAEKLIGVQSRHTFPAINGIPYQLQGLK |
| *B. candidus F5* | VNPEKDIHSGLIGPILICQKGMIDEYNRPIDIREFVLFFMVFDEERSWYFAKSDKRTRAEKLIGVQSRHTFPAINGIPYQLQGLK |
| *N. helvetica F5* | VNPEKDIHSGLIGPILICHKHMIDEYNRPIDIREFVLLFMVFDEERSWYFEKSDKRTRTERLLGVQSRHTFPAINGIPYQLQGLK |

| | 1531 … 1615 |
|---|---|
| *P. textilis vF5* (short) | MYKDENVHWHLLNMGGPKDIHVVNFHGQTFTEEGREDNQLGVLPLLPGTFASIKMKPSKIGTWLLETEVGENQERGMQALFTVID |
| *O. microlepidotus vF5* (short) | MYKDENVHWHLLNMGGPKDIHVVNFHGQTFTEEGREDNQLGVLPLLPGTFASIKMKPSKIGTWLLETEVGENQERGMQALFTVID |
| *P. textilis F5* (short) | MYKDENVHWHLLNMGGPKDIHVVNFHGQTFTEEGREDNQLGVLPLLPGTFASIKMKPSKIGTWLLETEVGENQERGMQALFTVID |
| *O. microlepidotus F5* (short) | MYKDENVHWHLLNMGGPKDIHVVNFHGQTFTEEGREDNQLGVLPLLPGTFTSIKMKPSKIGTWLLETEVGENQERGMQALFTVID |
| *P. textilis F5* | MYKDENVHWHLLNMGGPKDIHVVNFHGQTFTEEGREDNQLGVLPLLPGTFASIKMKPSKIGTWLLETEVGENQERGMQALFTVID |
| *O. microlepidotus F5* | MYKDENVHWHLLNMGGPKDIHVVNFHGQTFTEEGREDNQLGVLPLLPGTFTSIKMKPSKIGTWLLETEVGENQERGMQALFTVID |
| *N. scutatus F5* | MYKDENVHWHLLNMGGPKDIHVVKFHGQTFTEEGMEDNQLGVLPLLPGTFTSIKMKPSKIGTWLLETEVGENRERGMQALFTVID |
| *B. candidus F5* | MYKDENVHWHLLNMGGPKDIHVVNFHGQTFTEEGREDNQLGVLPLLPGTFTSIKMKPSKIGTWLLETEVGEYQERGMQALFTVID |
| *N. helvetica F5* | MYKEENVHWHLLNMGGPKDIHVVHFHGQTFTEEGREDNQLGVLPLLPGTFTTIKMKPSKIGTWLLKTEVGEYQERGMKALFTVID |

| | 1616 … 1700 |
|---|---|
| *P. textilis vF5* (short) | KDCKLPMGLASGIIQDSQISASGHVGYWEPKLARLNNTAIFNAWSIIKKEHEHPWIQIDLQRQVVITGIQTQGTVQLLQHSYTVE |
| *O. microlepidotus vF5* (short) | KDCKLPMGLASGIIQDSQISASGHVEYWEPKLARLNNTGMFNAWSIIKKEHEHPWIQIDLQRQVVITGIQTQGTVQLLKHSYTVE |
| *P. textilis F5* (short) | KDCKLPMGLASGIIQDSQISASGHVGYWEPKLARLNNTGKYNAWSIIKKEHEHPWIQIDLQRQVVITGIQTQGAMQLLKHLYTVE |
| *O. microlepidotus F5* (short) | KDCKLPMGLASGIIQDSQISASGHVGYWEPKLARLNNPGKYNAWSIIKKEHELPWIQIDLQRQVVITGIQTQGAMQLLKHLYTVE |
| *P. textilis F5* | KDCKLPMGLASGIIQDSQISASGHVGYWEPKLARLNNTGKYNAWSIIKKEHEHPWIQIDLQRQVVITGIQTQGAMQLLKHLYTVE |
| *O. microlepidotus F5* | KDCKLPMGLASGIIQDSQISASGHVGYWEPKLARLNNPGKYNAWSIIKKEHELPWIQIDLQRQVVITGIQTQGAMQLLKHLYTVE |
| *N. scutatus F5* | KDCKLPMGLASGIIQDSQISASGHVGYWEPKLARLNNPGKYNAWSIIKKEHEHPWIQIDLQRQVVITGIQTQGAMQLLKHLYTVE |
| *B. candidus F5* | KDCKLPMGLASGIIQDSQITASGHVGYWEPKLARLNNTGKYNAWSIIKREHEHPWIQVDLQRQVVITGIQTQGAMQLLKHLYTVE |
| *N. helvetica F5* | KDCKLPMGLESGIIQDSQIIASDHLGFWEPKLARLNNAGKYNAWSTIKKENEHPWIQVDLQREVVITGIQTQGAMQLVKHLYTVE |

| | 1701 … 1785 |
|---|---|
| *P. textilis vF5* (short) | YFVTYSEDGQNWITFKGRHSETQMHFEGNSDGTTVKENHIDPPIIARYIRLHPTKFYNRPTFRIELLGCEVEGCSVPLGMESGAI |
| *O. microlepidotus vF5* (short) | YFVTYSKDGQNWITFKGRHSETQMHFEGNSDGTTVKENHIDPPIIARYIRLHPTKFYNTPTFRIELLGCEVEGCSVPLGMESGAI |
| *P. textilis F5* (short) | YFFTYSKDGQNWITFKGRHSETQMHFEGNSDGTTVKENHIDPPIIARYIRLHPTKFHNRPTFRIELLGCEVEGCSVPLGMESGAI |
| *O. microlepidotus F5* (short) | YFVTYSKDGQNWITFKGRHSETQMHFEGNSDGTTVKENHTDPPIIARYIRLHPTKFHNRPTFRIELLGCEVEGCSVPLGMESGAI |
| *P. textilis F5* | YFFTYSKDGQNWITFKGRHSETQMHFEGNSDGTTVKENHIDPPIIARYIRLHPTKFHNRPTFRIELLGCEVEGCSVPLGMESGAI |
| *O. microlepidotus F5* | YFVTYSKDGQNWITFKGRHSETQMHFEGNSDGTTVKENHTDPPIIARYIRLHPTKFHNRPTFRIELLGCEVEGCSVPLGMESGAI |
| *N. scutatus F5* | YFVTYSKDGQNWITFKGRHSETQMHFEGNSDGSTVKENHIDPPIIARYIRLHPTKFHNRPTFRIELLGCEVEGCSVPLGMESGAI |
| *B. candidus F5* | YFVTYSKDGQKWITFKGRHSKTQMHFKGNSDGTKVKENHIDPPIIARYIRLHPTKFYNRPTFRIELLGCEVEGCSVPLGMESGAI |
| *N. helvetica F5* | YCVAYSKDGRKWITFKGRHPKTEMHFEGNSDGTKVKENHIDPPIIARYIRLHPTKFYNRPTFRIELLGCEVEGCSMPLGMESGAI |

| | 1786 … 1870 |
|---|---|
| *P. textilis vF5* (short) | KNSEITASSYKKTWWSSWEPFLARLNLEGGTNAWQPEVNNKDQWLQIDLQHLTKITSIITQGATSMTTSMYVKTFSIHYTDDNST |
| *O. microlepidotus vF5* (short) | KNSEITASSYKKTWWSSWEPFLARLNLEGGTNAWQPEVNNKDQWLQIDLQHLTKITSIITQGATSMTTAMYVKTFSIHYTDDNST |
| *P. textilis F5* (short) | KNSEITASSYKKTWWSSWEPSLARLNLKGRTNAWQPKVNNKDQWLQIDLQHLTKITSIITQGATSMTTSMYVKTFSIHYTDDNST |
| *O. microlepidotus F5* (short) | KDSEITASSYKKTWWSSWEPFLARLNLKGRTNAWQPKVNNKDQWLQIDLQHLTKITSIITQGATSMTTAMYVKTFSIHYTDDNST |
| *P. textilis F5* | KNSEITASSYKKTWWSSWEPSLARLNLKGRTNAWQPKVNNKDQWLQIDLQHLTKITSIITQGATSMTTSMYVKTFSIHYTDDNST |
| *O. microlepidotus F5* | KDSEITASSYKKTWWSSWEPFLARLNLKGRTNAWQPKVNNKDQWLQIDLQHLTKITSIITQGATSMTTAMYVKTFSIHYTDDNST |
| *N. scutatus F5* | KNSEITASSYKKTWWSSWEPSLARLNLKGRTNAWQPKVNNKDQWLQIDLQHLTKITSIITQGATSMSTAMYVKTFSIHYTDDNST |
| *B. candidus F5* | KNSEITASSYKKTWWSSWEPSLARLNLKGRTNAWQPKVNNKDQWLQIDLQHLTKITSIVTQGATSMSTPMYVKTFSIHYSDDNST |
| *N. helvetica F5* | KNSEITASSYKKTWWSSWEPSLARLNLKGRTNAWQPKVNNKDQWLQIDLKHLTKITSIITQGATSMSTEMYVKTFSIHYSDENST |

| | 1871 … 1931 |
|---|---|
| *P. textilis vF5* (short) | WKPYLDVRTSMEKVFTGNINSDGHVKHFFKPPILSRFIRIIPKTWNQYIALRIELFGCEVF |
| *O. microlepidotus vF5* (short) | WKPYLDVRTSMEKVFTGNINSDGHVKHFFKPPILSRFIRIIPKTWNQYIALRIELFGCEVF |
| *P. textilis F5* (short) | WKPYLDVRTSMEKVFTGNINSDGHVKHFFKPPILSRFIRIIPKTWNQYIALRIELFGCEVF |
| *O. microlepidotus F5* (short) | WKPYLDVRTSMEKVFTGNINTDGHVKHFFKPPILSRFIRIIPKTWNQYIALRIELFGCEVF |
| *P. textilis F5* | WKPYLDVRTSMEKVFTGNINSDGHVKHFFKPPILSRFIRIIPKTWNQYIALRIELFGCEVF |
| *O. microlepidotus F5* | WKPYLDVRTSMEKVFTGNINTDGHVKHFFKPPILSRFIRIIPKTWNQYIALRIELFGCEVF |
| *N. scutatus F5* | WKPYLDVCTSMEKVFTGNINTDGHVKHFFKPPILSRFIRIIPKTWNQYIALRIELFGCEVF |
| *B. candidus F5* | WKPYLDVCTSMEKVFTGNMNTDGHVKHFFIPPILSRFIRIIPKTWNQYIALRIELFGCDVF |
| *N. helvetica F5* | WKPYLDVCTSTEKVFTGNVNTDGHVKHFFKPPILSRFIRIIPKTWNRYIALRIELFGCDVF |

**Fig S8. The alternatively spliced venom factor V protein lacks a large portion of its B-domain region.** Amino acid sequence alignment of hypothetical protein translations of *F5* genes**.** The canonical coagulation factor V isoforms are denoted as *vF5* or *F5*, whereas the shorter alternatively spliced isoforms are denoted as *vF5* short or *F5* short. Residues shown in grey correspond to the signal peptide, and residues shown in bold indicate the B-domain region.

| | |
|---|---|
| *O.microlepidotus vF5* | MGRYSVSPVPKCLLLMFLGWSGLKYYQVNAAQLREYRIAAQLEDWDY**N**PQPEELSRLSESELTFKKIVYREYELDFKQEKPRDEL |
| *O. microlepidotus vF5* (short) | MGRYSVSPVPKCLLLMFLGWSGLKYYQVNAAQLREYRIAAQLEDWDY**N**PQPEELSRLSESELTFKKIVYREYELDFKQEKPRDEL |
| *O. microlepidotus F5* | MGRYSVSPVPKCLLLMFLGWSGLKYYQVNAAQLREYRIAAQLEDWDY**S**PQPEELSRLSESELTFKKIVYREYELDFKQEKPRDEL |
| transcript/12353 | MGRYSVSPVPKCLLLMFLGWSGLKYYQVNAAQLREYRIAAQLEDWDY**N**PQPEELSRLSESELTFKKIVYREYELDFKQEKPRDEL |
| *O. microlepidotus F5* (short) | MGRYSVSPVPKCLLLMFLGWSGLKYYQVNAAQLREYRIAAQLEDWDY**S**PQPEELSRLSESELTFKKIVYREYELDFKQEKPRDEL |
| transcript/61854 | MGRYSVSPVPKCLLLMFLGWSGLKYYQVNAAQLREYRIAAQLEDWDY**N**PQPEELSRLSESELTFKKIVYREYELDFKQEKPRDEL |

| | |
|---|---|
| *O.microlepidotus vF5* | SGLLGPTLRGEVGDILIIYFKNFATQPVSIHPQSAVYNKWSEGSSYSDGTSDVERLDDAVPPGQSFKYVWNITAEIGPKKADPPC |
| *O. microlepidotus vF5* (short) | SGLLGPTLRGEVGDILIIYFKNFATQPVSIHPQSAVYNKWSEGSSYSDGTSDVERLDDAVPPGQSFKYVWNITAEIGPKKADPPC |
| *O. microlepidotus F5* | SGLLGPTLRGEVGDILIIYFKNFATQPVSIHPQSAVYNKWSEGSSYSDGTSDVERLDDAVPPGQSFKYVWNITAEIGPKKADPPC |
| transcript/12353 | SGLLGPTLRGEVGDILIIYFKNFATQPVSIHPQSAVYNKWSEGSSYSDGTSDVERLDDAVPPGQSFKYVWNITAEIGPKKADPPC |
| *O. microlepidotus F5* (short) | SGLLGPTLRGEVGDILIIYFKNFATQPVSIHPQSAVYNKWSEGSSYSDGTSDVERLDDAVPPGQSFKYVWNITAEIGPKKADPPC |
| transcript/61854 | SGLLGPTLRGEVGDILIIYFKNFATQPVSIHPQSAVYNKWSEGSSYSDGTSDVERLDDAVPPGQSFKYVWNITAEIGPKKADPPC |

| | |
|---|---|
| *O.microlepidotus vF5* | LTY**A**YYSHVNMVRDFNSGLIGALLICKEGSLNANGAQKFFNREYVLMFSVFDESKNWYRKPSLQYTINGFANGTLPDVQACAYDH |
| *O. microlepidotus vF5* (short) | LTY**A**YYSHVNMVRDFNSGLIGALLICKEGSLNANGAQKFFNREYVLMFSVFDESKNWYRKPSLQYTINGFANGTLPDVQACAYDH |
| *O. microlepidotus F5* | LTY**S**YYSHVNMVRDFNSGLIGALLICKEGSLNANGAQKFFNREYVLMFSVFDESKNWYRKPSLQYTINGFANGTLPDVQACAYDH |
| transcript/12353 | LTY**A**YYSHVNMVRDFNSGLIGALLICKEGSLNANGAQKFFNREYVLMFSVFDESKNWYRKPSLQYTINGFANGTLPDVQACAYDH |
| *O. microlepidotus F5* (short) | LTY**S**YYSHVNMVRDFNSGLIGALLICKEGSLNANGAQKFFNREYVLMFSVFDESKNWYRKPSLQYTINGFANGTLPDVQACAYDH |
| transcript/61854 | LTY**A**YYSHVNMVRDFNSGLIGALLICKEGSLNANGAQKFFNREYVLMFSVFDESKNWYRKPSLQYTINGFANGTLPDVQACAYDH |

| | |
|---|---|
| *O.microlepidotus vF5* | ISWHLIGMSSSPEIFSVHFNGQTLEQNHYKVSTINLVGGASVTANMSVSRTGKWLISSLVAKHLQAGMYGYLNIKDCG**HPN**TLTR |
| *O. microlepidotus vF5* (short) | ISWHLIGMSSSPEIFSVHFNGQTLEQNHYKVSTINLVGGASVTANMSVSRTGKWLISSLVAKHLQAGMYGYLNIKDCG**HPN**TLTR |
| *O. microlepidotus F5* | ISWHLIGMSSSPEIFSVHFNGQTLEQNHYKVSTINLVGGASVTANMSVSRTGKWLISSLVAKHLQAGMYGYLNIKDCG**NPD**TLTR |
| transcript/12353 | ISWHLIGMSSSPEIFSVHFNGQTLEQNHYKVSTINLVGGASVTANMSVSRTGKWLISSLVAKHLQAGMYGYLNIKDCG**NPD**TLTR |
| *O. microlepidotus F5* (short) | ISWHLIGMSSSPEIFSVHFNGQTLEQNHYKVSTINLVGGASVTANMSVSRTGKWLISSLVAKHLQAGMYGYLNIKDCG**NPD**TLTR |
| transcript/61854 | ISWHLIGMSSSPEIFSVHFNGQTLEQNHYKVSTINLVGGASVTANMSVSRTGKWLISSLVAKHLQAGMYGYLNIKDCG**NPD**TLTR |

| | |
|---|---|
| *O.microlepidotus vF5* | KLSFRELRRIMNWEYFIAAEEITWDYAPEIPSSVDRRYKAQYLDNFSNFIGKKYKKAVFRQY**E**DGNFTKPTYAIWPKERGILGPV |
| *O. microlepidotus vF5* (short) | KLSFRELRRIMNWEYFIAAEEITWDYAPEIPSSVDRRYKAQYLDNFSNFIGKKYKKAVFRQY**E**DGNFTKPTYAIWPKERGILGPV |
| *O. microlepidotus F5* | KLSFRELRRIMNWEYFIAAEEITWDYAPEIPSSVDRRYKAQYLDNFSNFIGKKYKKAVFRQY**K**DGNFTKPTYAIWPKERGILGPV |
| transcript/12353 | KLSFRELRRIMNWEYFIAAEEITWDYAPEIPSSVDRRYKAQYLDNFSNFIGKKYKKAVFRQY**K**DGNFTKPTYAIWPKERGILGPV |
| *O. microlepidotus F5* (short) | KLSFRELRRIMNWEYFIAAEEITWDYAPEIPSSVDRRYKAQYLDNFSNFIGKKYKKAVFRQY**K**DGNFTKPTYAIWPKERGILGPV |
| transcript/61854 | KLSFRELRRIMNWEYFIAAEEITWDYAPEIPSSVDRRYKAQYLDNFSNFIGKKYKKAVFRQY**K**DGNFTKPTYAIWPKERGILGPV |

| | |
|---|---|
| *O.microlepidotus vF5* | I**K**AKVRDT**VT**IVFKNLASRPYSIYVHGVSVSKDAEGAIYPSDPKENITHGKAVEPGQVYTYKWTVLDTDEPTVKDSECITKLYHS |
| *O. microlepidotus vF5* (short) | I**K**AKVRDT**VT**IVFKNLASRPYSIYVHGVSVSKDAEGAIYPSDPKENITHGKAVEPGQVYTYKWTVLDTDEPTVKDSECITKLYHS |
| *O. microlepidotus F5* | I**R**AKVRDT**IS**IVFKNLASRPYSIYVHGVSVSKDAEGAIYPSDPKENITHGKAVEPGQVYTYKWTVLDTDEPTVKDSECITKLYHS |
| transcript/12353 | I**R**AKVRDT**IS**IVFKNLASRPYSIYVHGVSVSKDAEGAIYPSDPKENITHGKAVEPGQVYTYKWTVLDTDEPTVKDSECITKLYHS |
| *O. microlepidotus F5* (short) | I**R**AKVRDT**IS**IVFKNLASRPYSIYVHGVSVSKDAEGAIYPSDPKENITHGKAVEPGQVYTYKWTVLDTDEPTVKDSECITKLYHS |
| transcript/61854 | I**R**AKVRDT**IS**IVFKNLASRPYSIYVHGVSVSKDAEGAIYPSDPKENITHGKAVEPGQVYTYKWTVLDTDEPTVKDSECITKLYHS |

| | |
|---|---|
| *O.microlepidotus vF5* | AVDMTRDIASGLIGPLLVCK**L**KALSVKGVQNKADVEQHAVFAVFDENKSWYLEDNIKKYCSNPSSVKKDDPKFYKSNVMYTLNGY |
| *O. microlepidotus vF5* (short) | AVDMTRDIASGLIGPLLVCK**L**KALSVKGVQNKADVEQHAVFAVFDENKSWYLEDNIKKYCSNPSSVKKDDPKFYKSNVMYTLNGY |
| *O. microlepidotus F5* | AVDMTRDIASGLIGPLLVCK**R**KALSVKGVQNKADVEQHAVFAVFDENKSWYLEDNIKKYCSNPSSVKKDDPKFYKSNVMYTLNGY |
| transcript/12353 | AVDMTRDIASGLIGPLLVCK**R**KALSVKGVQNKADVEQHAVFAVFDENKSWYLEDNIKKYCSNPSSVKKDDPKFYKSNVMYTLNGY |
| *O. microlepidotus F5* (short) | AVDMTRDIASGLIGPLLVCK**R**KALSVKGVQNKADVEQHAVFAVFDENKSWYLEDNIKKYCSNPSSVKKDDPKFYKSNVMYTLNGY |
| transcript/61854 | AVDMTRDIASGLIGPLLVCK**R**KALSVKGVQNKADVEQHAVFAVFDENKSWYLEDNIKKYCSNPSSVKKDDPKFYKSNVMYTLNGY |

| | |
|---|---|
| *O.microlepidotus vF5* | ASDRTEVLGFHQSEVV**Q**WHLTSVGTVDEIVPVHLSGHTFLSKGKHQDILNLFPMSGESATVTMDNLGTWLLSSWGSCEMSNGMRL |
| *O. microlepidotus vF5* (short) | ASDRTEVLGFHQSEVV**Q**WHLTSVGTVDEIVPVHLSGHTFLSKGKHQDILNLFPMSGESATVTMDNLGTWLLSSWGSCEMSNGMRL |
| *O. microlepidotus F5* | ASDRTEVLGFHQSEVV**E**WHLTSVGTVDEIVPVHLSGHTFLSKGKHQDILNLFPMSGESATVTMDNLGTWLLSSWGSCEMSNGMRL |
| transcript/12353 | ASDRTEVLGFHQSEVV**E**WHLTSVGTVDEIVPVHLSGHTFLSKGKHQDILNLFPMSGESATVTMDNLGTWLLSSWGSCEMSNGMRL |
| *O. microlepidotus F5* (short) | ASDRTEVLGFHQSEVV**E**WHLTSVGTVDEIVPVHLSGHTFLSKGKHQDILNLFPMSGESATVTMDNLGTWLLSSWGSCEMSNGMRL |
| transcript/61854 | ASDRTEVLGFHQSEVV**E**WHLTSVGTVDEIVPVHLSGHTFLSKGKHQDILNLFPMSGESATVTMDNLGTWLLSSWGSCEMSNGMRL |

| | |
|---|---|
| *O.microlepidotus vF5* | RFLDANYDDEDEGNEEEEEDDGDIFADIFSPPEVVKKKE**EV**PVNFV**P**DPESD**AL**AKELGLLDDEDN**P**E**Q**S**RS**EQTEDDEEQLMIA |
| *O. microlepidotus vF5* (short) | RFLDANYDDEDEGNEEEEEDDGDIFADIFSPPEVVKKKE**EV**PVNFV**P**DPESD**AL**AKELGLLDDEDN**P**E**Q**S**RS**EQTEDDEEQLMIA |
| *O. microlepidotus F5* | RFLDANYDDEDEGNEEEEEDDGDIFADIFSPPEVVKKKE**KD**PVNFV**S**DPESD**KI**AKELGLLDDEDN**Q**E**E**S**HN**EQTEDDEEQLMIA |
| transcript/12353 | RFLDANYDDEDEGNEEEEEDDGDIFADIFSPPEVVKKKE**KD**PVNFV**S**DPESD**KI**AKELGLLDDEDN**Q**E**E**S**HN**EQTEDDEEQLMIA |
| *O. microlepidotus F5* (short) | RFLDANYDDEDEGNEEEEEDDGDIFADIFSPPEVVKKKE**KD**PVNFV**S**DPESD**KI**AKELGLLDDEDN**Q**E**E**S**HN**EQTEDDEEQLMIA |
| transcript/61854 | RFLDANYDDEDEGNEEEEEDDGDIFADIFSPPEVVKKKE**KD**PVNFV**S**DPESD**KI**AKELGLLDDEDN**Q**E**E**S**HN**EQTEDDEEQLMIA |

| | |
|---|---|
| *O.microlepidotus vF5* | **S**MLGLRSFKGSVAEEEL**KH**TALALEEDAHASGELSLNTHLVLINSTDAMRGNEELGT**Q**NNSYVNASETIYANYTSSTGEILISNI |
| *O. microlepidotus vF5* (short) | **S**MLGLRSFKGSVAEEEL**KH**TALALEEDAHAS------------------------------------------------------ |
| *O. microlepidotus F5* | **T**MLGLRSFKGSVAEEEL**NL**TALALEEDAHASGELSLNTHLVLINSTDAMRGNEELGT**R**NNSYVNASETIYANYTSSTGEILISNI |
| transcript/12353 | **T**MLGLRSFKGSVAEEEL**NL**TALALEEDAHASGELSLNTHLVLINSTDAMRGNEELGT**R**NNSYVNASETIYANYTSSTGEILISNI |
| *O. microlepidotus F5* (short) | **T**MLGLRSFKGSVAEEEL**NL**TALALEEDAHAS------------------------------------------------------ |
| transcript/61854 | **T**MLGLRSFKGSVAEEEL**NL**TALALEEDAHAS------------------------------------------------------ |

| | |
|---|---|
| *O.microlepidotus vF5* | TAMIGILL**S**KKSQNEENITQISDSYKQMLAAEYLKAGRGLQS**K**DIEDLMSPEIIPSLKEQEENTFEKDIWDKKGERQILTEATLH |
| *O. microlepidotus vF5* (short) | ------------------------------------------------------------------------------------- |
| *O. microlepidotus F5* | TAMIGILL**S**KKSQNEENITQISDSYKQMLAAEYLKAGRGLQS**Q**DIEDLMSPEIIPSLKEQEENTFEKDIWDKKGERQILTEATLH |
| transcript/12353 | TAMIGILL**F**KKSQNEENITQISDSYKQMLAAEYLKAGRGLQS**Q**DIEDLMSPEIIPSLKEQEENTFEKDIWDKKGERQILTEATLH |
| *O. microlepidotus F5* (short) | ------------------------------------------------------------------------------------- |
| transcript/61854 | ------------------------------------------------------------------------------------- |

| | |
|---|---|
| *O.microlepidotus vF5* | ALKEMHALFFYVQQKRNLSAFNITSEDMKFLPSTENDSTIDNSKANYFS**YD**YDYDYDYKEEEATAAANVEFSKVIINTISDDKKT |
| *O. microlepidotus vF5* (short) | ------------------------------------------------------------------------------------- |
| *O. microlepidotus F5* | ALKEMHALFFYVQQKRNLSAFNITSEDMKFLPSTENDSTIDNSKANYFS--YDYDYDYKEEEATAAANVEFSKVIINTISDDKKT |
| transcript/12353 | ALKEMHALFFYVQQKRNLSAFNITSEDMKFLPSTENDSTIDNSKANYFS--YDYDYDYKEEEATAAANVEFSKVIINTISDDKKT |
| *O. microlepidotus F5* (short) | ------------------------------------------------------------------------------------- |
| transcript/61854 | ------------------------------------------------------------------------------------- |
| *O.microlepidotus vF5* | VENNRSEVNLVSVPSIMPQPENLTSDATLDNFVSTTISKTTK**P**WNSSSHQRQKCLPKNTDMEWKKENYDTLADIPNDVDNKFQNS |
| *O. microlepidotus vF5* (short) | ------------------------------------------------------------------------------------- |
| *O. microlepidotus F5* | VENNRSEVNLVSVPSIMPQPENLTSDATLDNFVSTTISKTTK**A**WNSSSHQRQKCLPKNTDMEWKKENYDTLADIPNDVDNKFQNS |
| transcript/12353 | VENNRSEVNLVSVPSIMPQPENLTSDATLDNFVSTTISKTTK**A**WNSSSHQRQKCLPKNTDMEWKKENYDTLADIPNDVDNKFQNS |
| *O. microlepidotus F5* (short) | ------------------------------------------------------------------------------------- |
| transcript/61854 | ------------------------------------------------------------------------------------- |
| *O.microlepidotus vF5* | SRNASFYQENAMFPGKRKEKTDNSNGQLELGQPGITNKHDSQKDENCTINISQSFIKIQ*KKKKPKVLTPRTSKHLELPRNLNNS |
| *O. microlepidotus vF5* (short) | ------------------------------------------------------------------------------------- |
| *O. microlepidotus F5* | SRNASFYQENAMFPGKRKEKTDNSNGQLELGQPGITNKHDSQKDENCTINISQSFIKIQ**R**KKKKPKVLTPRTSKHLELPRNLNNS |
| transcript/12353 | SRNASFYQENAMFPGKRKEKTDNSNGQLELGQPGITNKHDSQKDENCTINISQSFIKIQ**R**KKKKPKVLTPRTSKHLELPRNLNNS |
| *O. microlepidotus F5* (short) | ------------------------------------------------------------------------------------- |
| transcript/61854 | ------------------------------------------------------------------------------------- |
| *O.microlepidotus vF5* | QLSNKPNHTKLLNEAKLTRKQDK**L**IVTIGLPVEDGDYQEYEIDNTDNDKSSSGSFEYELMYYNNPYTTDPRIDSNSARN**S**DDIAG |
| *O. microlepidotus vF5* (short) | --------------------------------------------------------------------DPRIDSNSARN**S**DDIAG |
| *O. microlepidotus F5* | QLSNKPNHTKLLNEAKLTRKQDK**F**IVTIGLPVEDGDYQEYEIDNTDNDKSSSGSFEYELMYYNNPYTTDPRIDSNSARN**P**DDIAG |
| transcript/12353 | QLSNKPNHTKLLNEAKLTRKQDK**L**IVTIGLPVEDGDYQEYEIDNTDNDKSSSGSFEYELMYYNNPYTTDPRIDSNSARN**P**DDIAG |
| *O. microlepidotus F5* (short) | --------------------------------------------------------------------DPRIDSNSARN**P**DDIAG |
| transcript/61854 | --------------------------------------------------------------------DPRIDSNSARN**P**DDIAG |
| *O.microlepidotus vF5* | RYLRTINR**R**NKRRYYIAAEEVLWDYSPIGKSQVRS**LP**AKTTFKKAIFRSYLDDTFQTPSTGGEYEKHLGILGPIIRAEVDDVIEV |
| *O. microlepidotus vF5* (short) | RYLRTINR**R**NKRRYYIAAEEVLWDYSPIGKSQVRS**LP**AKTTFKKAIFRSYLDDTFQTPSTGGEYEKHLGILGPIIRAEVDDVIEV |
| *O. microlepidotus F5* | RYLRTINR**G**NKRRYYIAAEEVLWDYSPIGKSQVRS**RA**AKTTFKKAIFRSYLDDTFQTPSTGGEYEKHLGILGPIIRAEVDDVIEV |
| transcript/12353 | RYLRTINR**G**NKRRYYIAAEEVLWDYSPIGKSQVRS**RA**AKTTFKKAIFRSYLDDTFQTPSTGGEYEKHLGILGPIIRAEVDDVIEV |
| *O. microlepidotus F5* (short) | RYLRTINR**G**NKRRYYIAAEEVLWDYSPIGKSQVRS**RA**AKTTFKKAIFRSYLDDTFQTPSTGGEYEKHLGILGPIIRAEVDDVIEV |
| transcript/61854 | RYLRTINR**G**NKRRYYIAAEEVLWDYSPIGKSQVRS**RA**AKTTFKKAIFRSYLDDTFQTPSTGGEYEKHLGILGPIIRAEVDDVIEV |
| *O.microlepidotus vF5* | QFRNLASRPYSLHAHGLLYEKSSEGRSYDD**N**SPELFKKDDAIMPNGTYTYVWQVPPRSGPTDNTEKCKSWAYYSGVNPEKDIHSG |
| *O. microlepidotus vF5* (short) | QFRNLASRPYSLHAHGLLYEKSSEGRSYDD**N**SPELFKKDDAIMPNGTYTYVWQVPPRSGPTDNTEKCKSWAYYSGVNPEKDIHSG |
| *O. microlepidotus F5* | QFRNLASRPYSLHAHGLLYEKSSEGRSYDD**K**SPELFKKDDAIMPNGTYTYVWQVPPRSGPTDNTEKCKSWAYYSGVNPEKDIHSG |
| transcript/12353 | QFRNLASRPYSLHAHGLLYEKSSEGRSYDD**K**SPELFKKDDAIMPNGTYTYVWQVPPRSGPTDNTEKCKSWAYYSGVNPEKDIHSG |
| *O. microlepidotus F5* (short) | QFRNLASRPYSLHAHGLLYEKSSEGRSYDD**K**SPELFKKDDAIMPNGTYTYVWQVPPRSGPTDNTEKCKSWAYYSGVNPEKDIHSG |
| transcript/61854 | QFRNLASRPYSLHAHGLLYEKSSEGRSYDD**K**SPELFKKDDAIMPNGTYTYVWQVPPRSGPTDNTEKCKSWAYYSGVNPEKDIHSG |
| *O.microlepidotus vF5* | LIGPILICQKGMIDKYNRTIDIREFVLFFMVFDEEKSWYFPKSDKST**CE**EKLIGVQS**H**HTFPAINGIPYQLQGLMMYKDENVHWH |
| *O. microlepidotus vF5* (short) | LIGPILICQKGMIDKYNRTIDIREFVLFFMVFDEEKSWYFPKSDKST**CE**EKLIGVQS**H**HTFPAINGIPYQLQGLMMYKDENVHWH |
| *O. microlepidotus F5* | LIGPILICQKGMIDKYNRTIDIREFVLFFMVFDEEKSWYFPKSDKST**RA**EKLIGVQS**R**HTFPAINGIPYQLQGLMMYKDENVHWH |
| transcript/12353 | LIGPILICQKGMIDKYNRTIDIREFVLFFMVFDEEKSWYFPKSDKST**RA**EKLIGVQS**R**HTFPAINGIPYQLQGLMMYKDENVHWH |
| *O. microlepidotus F5* (short) | LIGPILICQKGMIDKYNRTIDIREFVLFFMVFDEEKSWYFPKSDKST**RA**EKLIGVQS**R**HTFPAINGIPYQLQGLMMYKDENVHWH |
| transcript/61854 | LIGPILICQKGMIDKYNRTIDIREFVLFFMVFDEEKSWYFPKSDKST**RA**EKLIGVQS**R**HTFPAINGIPYQLQGLMMYKDENVHWH |
| *O.microlepidotus vF5* | LLNMGGPKDIHVVNFHGQTFTEEGREDNQLGVLPLLPGTF**A**SIKMKPSKIGTWLLETEVGENQERGMQALFTVIDKDCKLPMGLA |
| *O. microlepidotus vF5* (short) | LLNMGGPKDIHVVNFHGQTFTEEGREDNQLGVLPLLPGTF**A**SIKMKPSKIGTWLLETEVGENQERGMQALFTVIDKDCKLPMGLA |
| *O. microlepidotus F5* | LLNMGGPKDIHVVNFHGQTFTEEGREDNQLGVLPLLPGTF**T**SIKMKPSKIGTWLLETEVGENQERGMQALFTVIDKDCKLPMGLA |
| transcript/12353 | LLNMGGPKDIHVVNFHGQTFTEEGREDNQLGVLPLLPGTF**T**SIKMKPSKIGTWLLETEVGENQERGMQALFTVIDKDCKLPMGLA |
| *O. microlepidotus F5* (short) | LLNMGGPKDIHVVNFHGQTFTEEGREDNQLGVLPLLPGTF**T**SIKMKPSKIGTWLLETEVGENQERGMQALFTVIDKDCKLPMGLA |
| transcript/61854 | LLNMGGPKDIHVVNFHGQTFTEEGREDNQLGVLPLLPGTF**T**SIKMKPSKIGTWLLETEVGENQERGMQALFTVIDKDCKLPMGLA |
| *O.microlepidotus vF5* | SGIIQDSQISASGHV**E**YWEPKLARLNN**T**G**MF**NAWSIIKKEHE**H**PWIQIDLQRQVVITGIQTQG**TV**QLLKH**S**YTVEYFVTYSKDGQ |
| *O. microlepidotus vF5* (short) | SGIIQDSQISASGHV**E**YWEPKLARLNN**T**G**MF**NAWSIIKKEHE**H**PWIQIDLQRQVVITGIQTQG**TV**QLLKH**S**YTVEYFVTYSKDGQ |
| *O. microlepidotus F5* | SGIIQDSQISASGHV**G**YWEPKLARLNN**P**G**KY**NAWSIIKKEHE**L**PWIQIDLQRQVVITGIQTQG**AM**QLLKH**L**YTVEYFVTYSKDGQ |
| transcript/12353 | SGIIQDSQISASGHV**G**YWEPKLARLNN**P**G**KY**NAWSIIKKEHE**H**PWIQIDLQRQVVITGIQTQG**AM**QLLKH**L**YTVEYFVTYSKDGQ |
| *O. microlepidotus F5* (short) | SGIIQDSQISASGHV**G**YWEPKLARLNN**P**G**KY**NAWSIIKKEHE**L**PWIQIDLQRQVVITGIQTQG**AM**QLLKH**L**YTVEYFVTYSKDGQ |
| transcript/61854 | SGIIQDSQISASGHV**G**YWEPKLARLNN**P**G**KY**NAWSIIKKEHE**H**PWIQIDLQRQVVITGIQTQG**AM**QLLKH**L**YTVEYFVTYSKDGQ |
| *O.microlepidotus vF5* | NWITFKGRHSETQMHFEGNSDGTTVKENH**I**DPPIIARYIRLHPTKF**Y**N**T**PTFRIELLGCEVEGCSVPLGMESGAIKNSEITASSY |
| *O. microlepidotus vF5* (short) | NWITFKGRHSETQMHFEGNSDGTTVKENH**I**DPPIIARYIRLHPTKF**Y**N**T**PTFRIELLGCEVEGCSVPLGMESGAIKNSEITASSY |
| *O. microlepidotus F5* | NWITFKGRHSETQMHFEGNSDGTTVKENH**T**DPPIIARYIRLHPTKF**H**N**R**PTFRIELLGCEVEGCSVPLGMESGAIKDSEITASSY |
| transcript/12353 | NWITFKGRHSETQMHFEGNSDGTTVKENH**I**DPPIIARYIRLHPTKF**H**N**R**PTFRIELLGCEVEGCSVPLGMESGAIKDSEITASSY |
| *O. microlepidotus F5* (short) | NWITFKGRHSETQMHFEGNSDGTTVKENH**T**DPPIIARYIRLHPTKF**H**N**R**PTFRIELLGCEVEGCSVPLGMESGAIKDSEITASSY |
| transcript/61854 | NWITFKGRHSETQMHFEGNSDGTTVKENH**I**DPPIIARYIRLHPTKF**H**N**R**PTFRIELLGCEVEGCSVPLGMESGAIKDSEITASSY |
| *O.microlepidotus vF5* | KKTWWSSWEPFLARLNL**E**G**G**TNAWQP**E**VNNKDQWLQIDLQHLTKITSIITQGATSMTTAMYVKTFSIHYTDDNSTWKPYLDVRTS |
| *O. microlepidotus vF5* (short) | KKTWWSSWEPFLARLNL**E**G**G**TNAWQP**E**VNNKDQWLQIDLQHLTKITSIITQGATSMTTAMYVKTFSIHYTDDNSTWKPYLDVRTS |
| *O. microlepidotus F5* | KKTWWSSWEPFLARLNL**K**G**R**TNAWQP**K**VNNKDQWLQIDLQHLTKITSIITQGATSMTTAMYVKTFSIHYTDDNSTWKPYLDVRTS |
| transcript/12353 | KKTWWSSWEPFLARLNL**K**G**R**TNAWQP**K**VNNKDQWLQIDLQHLTKITSIITQGATSMTTAMYVKTFSIHYTDDNSTWKPYLDVRTS |
| *O. microlepidotus F5* (short) | KKTWWSSWEPFLARLNL**K**G**R**TNAWQP**K**VNNKDQWLQIDLQHLTKITSIITQGATSMTTAMYVKTFSIHYTDDNSTWKPYLDVRTS |
| transcript/61854 | KKTWWSSWEPFLARLNL**K**G**R**TNAWQP**K**VNNKDQWLQIDLQHLTKITSIITQGATSMTTAMYVKTFSIHYTDDNSTWKPYLDVRTS |

| | |
|---|---|
| *O.microlepidotus vF5* | MEKVFTGNIN**S**DGHVKHFFKPPILSRFIRIIPKTWNQYIALRIELFGCEVF |
| *O. microlepidotus vF5* (short) | MEKVFTGNIN**S**DGHVKHFFKPPILSRFIRIIPKTWNQYIALRIELFGCEVF |
| *O. microlepidotus F5* | MEKVFTGNIN**T**DGHVKHFFKPPILSRFIRIIPKTWNQYIALRIELFGCEVF |
| transcript/12353 | MEKVFTGNIN**T**DGHVKHFFKPPILSRFIRIIPKTWNQYIALRIELFGCEVF |
| *O. microlepidotus F5* (short) | MEKVFTGNIN**T**DGHVKHFFKPPILSRFIRIIPKTWNQYIALRIELFGCEVF |
| transcript/61854 | MEKVFTGNIN**T**DGHVKHFFKPPILSRFIRIIPKTWNQYIALRIELFGCEVF |

**Fig S9. Full-length *F5* transcripts are present in inland taipan (*Oxyuranus microlepidotus*) venom gland tissue.** Amino acid alignment of the hypothetical protein isoform translations encoded by the *F5* and *vF5* paralogs of *O. microlepidotus*, together with representative full-length translated Iso-seq reads of both isoforms from venom gland tissue corresponding to each *F5* paralog. The alignment provides qualitative evidence that both *F5* alternative splice variants are expressed in venom gland tissue. A single-base-pair indel in transcript/12353 that was absent in all other transcripts and therefore likely represents a technical artifact, was manually corrected to restore the reading frame. Amino acid residues that differ between the *F5* and vF5 genes are shown in bold.

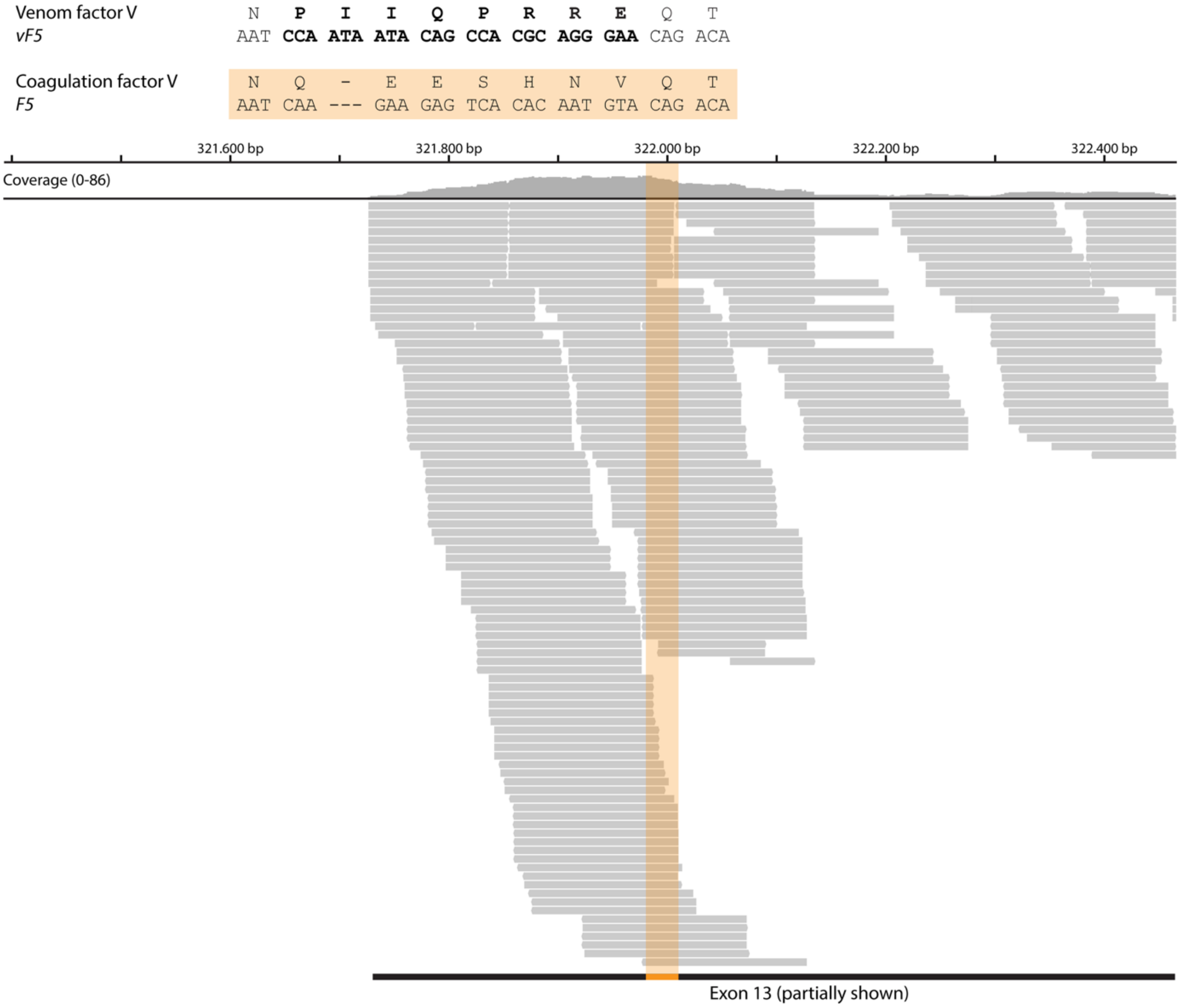

**Fig S10. The hemostatic *F5* gene is expressed in Eastern brown snake (*Pseudonaja textilis*) venom gland tissue.** One distinguishing feature between the *F5* and *vF5* paralogs in *P. textilis* is a unique structural variant in exon 13 of the *vF5* gene. The unique protein sequence and underlying codons are shown in bold for the *vF5* gene, with the corresponding protein sequence and codons in the *F5* gene highlighted in orange. We identified multiple short-read RNA-seq reads from *P. textilis* venom gland tissue that map to/cover this *F5*-specific region (highlighted in orange), providing additional evidence that the hemostatic *F5* gene is also transcribed in the venom gland of *P. textilis*.

| | |
|---|---|
| *Pseudonaja textilis vF5* | MGRYSVSPVPKCLLLMFLGWSGLKYYQVNAAQLREYHIAAQLEDWDYNPQPEELSR**LSESDLTFKK**IVYR**EYELDFK**QEKPRDA |
| *Pseudonaja textilis F5* (short) | MGRYSVSPVPKCLLLMFLGWSGLKYYQVNAAQLREYRIAAQLEDWDYNPQPEELSR**LSESDLTFKK**IVYR**EYELDFK**QEKPRDA |
| *Pseudonaja textilis F5* | MGRYSVSPVPKCLLLMFLGWSGLKYYQVNAAQLREYRIAAQLEDWDYNPQPEELSR**LSESDLTFKK**IVYR**EYELDFK**QEKPRDA |
| *Pseudonaja textilis vF5* | LSGLLGPTLRGEVGDSLIIYFKNFATQPVSIHPQSAVYNK**WSEGSSYSDGTSDVERLDDAVPPGQSFK**YVWNITAEIGPKKADP |
| *Pseudonaja textilis F5* (short) | LSGLLGPTLRGEVGDILIIYFKNFATQPVSIHPQSAVYNK**WSEGSSYSDGTSDVERLDDAVPPGQSFK**YVWNITAEIGPKKADP |
| *Pseudonaja textilis F5* | LSGLLGPTLRGEVGDILIIYFKNFATQPVSIHPQSAVYNK**WSEGSSYSDGTSDVERLDDAVPPGQSFK**YVWNITAEIGPKKADP |
| *Pseudonaja textilis vF5* | PCLTYAYYSHVNMVRDFNSGLIGALLICK**EGSLNANGSQK**FFNREYVLMFSVFDESKNWYRKPSLQYTINGFANGTLPDVQACA |
| *Pseudonaja textilis F5* (short) | PCLTYAYYSHVNMVRDFNSGLIGALLICK**EGSLNANGAQK**FFNREYVLMFSVFDESKNWYRKPSLQYTINGFANGTLPDVQACA |
| *Pseudonaja textilis F5* | PCLTYAYYSHVNMVRDFNSGLIGALLICK**EGSLNANGAQK**FFNREYVLMFSVFDESKNWYRKPSLQYTINGFANGTLPDVQACA |
| *Pseudonaja textilis vF5* | YDHISWHLIGMSSSPEIFSVHFNGQTLEQNHYKVSTINLVGGASVTADMSVSRTGKWLISSLVAKHLQAGMYGYLNIK**DCGNPD** |
| *Pseudonaja textilis F5* (short) | YDHISWHLIGMSSSPEIFSVHFNGQTLEQNHYKVSTINLVGGASVTANMSVSRTGKWLISSLVAKHLQAGMYGYLNIK**DCGNPD** |
| *Pseudonaja textilis F5* | YDHISWHLIGMSSSPEIFSVHFNGQTLEQNHYKVSTINLVGGASVTANMSVSRTGKWLISSLVAKHLQAGMYGYLNIK**DCGNPD** |
| *Pseudonaja textilis vF5* | **TLTR**KLSFRELMKIKNWEYFIAAEEITWDYAPEIPSSVDRRYKAQYLDNFSNFIGKKYKKAVFRQYEDGNFTKPTYAIWPKER**G** |
| *Pseudonaja textilis F5* (short) | **TLTR**KLSFRELRRIMNWEYFIAAEEITWDYAPEIPSSVDRRYKAQYLDNFSNFIGKKYKKAVFR**QYKDSNFTK**PTYAIWPKERG |
| *Pseudonaja textilis F5* | **TLTR**KLSFRELRRIMNWEYFIAAEEITWDYAPEIPSSVDRRYKAQYLDNFSNFIGKKYKKAVFR**QYKDSNFTK**PTYAIWPKERG |
| *Pseudonaja textilis vF5* | **ILGPVIK**AK**VRDTVTIVFK**NLASRPYSIYVHGVSVSKDAEGAIYPSDPKENITHGK**AVEPGQVYTYK**WTVLDTDEPTVK**DSECI** |
| *Pseudonaja textilis F5* (short) | ILGPVIRAK**VRDTISIVFK**NLASRPYSIYVHGVSVSKDAEGAIYPSDPKENITHGK**AVEPGQVYTYK**WTVLDTDEPTVK**DSECI** |
| *Pseudonaja textilis F5* | ILGPVIRAK**VRDTISIVFK**NLASRPYSIYVHGVSVSKDAEGAIYPSDPKENITHGK**AVEPGQVYTYK**WTVLDTDEPTVK**DSECI** |
| *Pseudonaja textilis vF5* | **TKLYHSAVDMTR**DIASGLIGPLLVCKHKALSVKGVQNKADVEQHAVFAVFDENKSWYLEDNIK**KYCSNPSAVKKDDPKFYK**SNV |
| *Pseudonaja textilis F5* (short) | **TKLYHSAVDMTR**DIASGLIGPLLVCKHKALSVKGVQNKADVEQHAVFAVFDENKSWYLEDNIK**KYCSNPSTVKKDDPKFYK**SNV |
| *Pseudonaja textilis F5* | **TKLYHSAVDMTR**DIASGLIGPLLVCKHKALSVKGVQNKADVEQHAVFAVFDENKSWYLEDNIK**KYCSNPSTVKKDDPKFYK**SNV |
| *Pseudonaja textilis vF5* | MYTLNGYASDRTEVLRFHQSEVVQWHLTSVGTVDEIVPVHLSGHTFLSKGKHQDILNLFPMSGESATVTMDNLGTWLLSSWGSC |
| *Pseudonaja textilis F5* (short) | MYTLNGYASDRTEVLGFHQSEVVEWHLTSVGTVDEIVPVHLSGHTFLSKGKHQDILNLFPMSGESATVTMDNLGTWLLSSWGSC |
| *Pseudonaja textilis F5* | MYTLNGYASDRTEVLGFHQSEVVEWHLTSVGTVDEIVPVHLSGHTFLSKGKHQDILNLFPMSGESATVTMDNLGTWLLSSWGSC |
| *Pseudonaja textilis vF5* | EMSNGMRLRFLDANYDDEDEGNEEEEEDDGDIFADIFIPSEVVKKKEEVPVNFVPDPESDALAKELGLIDDEGNPIIQPR**REQT** |
| *Pseudonaja textilis F5* (short) | EMSNGMRLRFLDANYDDEDEGNEEEEEDDGDIFADIFIPPEVVKKKEKDPVNFVSDPESDKIAKELGLLDDEDNQ-EESHNVQT |
| *Pseudonaja textilis F5* | EMSNGMRLRFLDANYDDEDEGNEEEEEDDGDIFADIFIPPEVVKKKEKDPVNFVSDPESDKIAKELGLLDDEDNQ-EESHNVQT |
| *Pseudonaja textilis vF5* | **EDDEEQLMK**ASMLGLRSFKGSVAEEELKHTALALEEDAHAS------------------------------------------- |
| *Pseudonaja textilis F5* (short) | EDDEEQLMIATMLGFRSFKGSVAEEELNLTALALEEDAHAS------------------------------------------- |
| *Pseudonaja textilis F5* | EDDEEQLMIATMLGFRSFKGSVAEEELNLTALALEEDAHASGELSLNTHLVLINSTDAMRGNEELGTQNNSYVNASETIYANYT |
| *Pseudonaja textilis vF5* | ------------------------------------------------------------------------------------ |
| *Pseudonaja textilis F5* (short) | ------------------------------------------------------------------------------------ |
| *Pseudonaja textilis F5* | SSTGEILISNITAMIGILLSKRSQNEENITQISDSYKEMLAAEYLKAGRGLQSKDIEDLMSPEIIPSLKEQEENTFEKDIWDKK |
| *Pseudonaja textilis vF5* | ------------------------------------------------------------------------------------ |
| *Pseudonaja textilis F5* (short) | ------------------------------------------------------------------------------------ |
| *Pseudonaja textilis F5* | GERQILTEATLHALKEMHALFFYVQQKRNLSAFNITSEGMKFLPSTENDSTIDNSKANYFSYDYDYDYKEEEATAAANVEFSKV |
| *Pseudonaja textilis vF5* | ------------------------------------------------------------------------------------ |
| *Pseudonaja textilis F5* (short) | ------------------------------------------------------------------------------------ |
| *Pseudonaja textilis F5* | IINTISDDKKTVENNRSEVNLVSVPSIMPQPENLTSDATLDNFVSTTISKTTKTWNSSSHQRQKCLPKNTDMEWKKENYDTLAD |
| *Pseudonaja textilis vF5* | ------------------------------------------------------------------------------------ |
| *Pseudonaja textilis F5* (short) | ------------------------------------------------------------------------------------ |
| *Pseudonaja textilis F5* | IPNDVDNKFQNSSRNASFYQENAMFPGKRKEKTDNSNGQLELGQPGITNKHDSQKDENCTINISQSFIKIQRKKKKPKVLTPRT |
| *Pseudonaja textilis vF5* | ---------------------------------------------------------------------------------DPR |
| *Pseudonaja textilis F5* (short) | ---------------------------------------------------------------------------------DPR |
| *Pseudonaja textilis F5* | SKHLELPRNLNNSQLSNKPNHTKLLNEAKLTRKQDKLIVTIGLPVEDGDYQEYDIDNTDNDRSSSGSFEYELMYYNNPYTTDPR |
| *Pseudonaja textilis vF5* | **IDSNSARNPDDIAGR**YLRTINRGNKRRYYIAAEEVLWDYSPIGKSQVRSRAAKTTFKKAIFRSYLDDTFQTPSTGGEYEKHLGI |
| *Pseudonaja textilis F5* (short) | **IDSNSARNPDDIAGR**YLRTINRGNKRRYYIAAEEVLWDYSPIGKSQVRSRAAKTTFKKAIFRSYLDDTFQTPSTGGEYEKHLGI |
| *Pseudonaja textilis F5* | **IDSNSARNPDDIAGR**YLRTINRGNKRRYYIAAEEVLWDYSPIGKSQVRSRAAKTTFKKAIFRSYLDDTFQTPSTGGEYEKHLGI |
| *Pseudonaja textilis vF5* | LGPIIRAEVDDVIEIQFRNLASR**PYSLHAHGLLYEK**SSEGR**SYDDKSPELFKK**DDAIMPNGTYTYVWQVPPR**SGPTDNTEK**CKS |
| *Pseudonaja textilis F5* (short) | LGPIIRAEVDDVIEVQFRNLASR**PYSLHAHGLLYEK**SSEGR**SYDDKSPELFKK**DDAIMPNGTYTYVWQVPPR**SGPTDNTEK**CKS |
| *Pseudonaja textilis F5* | LGPIIRAEVDDVIEVQFRNLASR**PYSLHAHGLLYEK**SSEGR**SYDDKSPELFKK**DDAIMPNGTYTYVWQVPPR**SGPTDNTEK**CKS |
| *Pseudonaja textilis vF5* | WAYYSGVNPEKDIHSGLIGPILICQK**GMIDKYNR**TIDIREFVLFFMVFDEEKSWYFPKSDKSTCEEKLIGVQSLHTFPAINGIP |
| *Pseudonaja textilis F5* (short) | WAYYSGVNPEKDIHSGLIGPILICQK**GMIDKYNR**TIDIREFVLFFMVFDEEKSWYFPKSDKSTRAEKLIGVQSRHTFPAINGIP |
| *Pseudonaja textilis F5* | WAYYSGVNPEKDIHSGLIGPILICQK**GMIDKYNR**TIDIREFVLFFMVFDEEKSWYFPKSDKSTRAEKLIGVQSRHTFPAINGIP |
| *Pseudonaja textilis vF5* | YQLQGLTMYKDENVHWHLLNMGGPK**DIHVVNFHGQTFTEEGR**EDNQLGVLPLLPGTFASIKMKPSKIGTWLLETEVGENQERGM |
| *Pseudonaja textilis F5* (short) | YQLQGLTMYKDENVHWHLLNMGGPK**DIHVVNFHGQTFTEEGR**EDNQLGVLPLLPGTFASIKMKPSKIGTWLLETEVGENQERGM |
| *Pseudonaja textilis F5* | YQLQGLTMYKDENVHWHLLNMGGPK**DIHVVNFHGQTFTEEGR**EDNQLGVLPLLPGTFASIKMKPSKIGTWLLETEVGENQERGM |

*Pseudonaja textilis vF5* QALFTVIDKDCKLPMGLASGIIQDSQISASGHVGYWEPKLARLNNTAIFNAWSIIKKEHEHPWIQIDLQRQVVITGIQTQGTVQ
*Pseudonaja textilis F5* (short) QALFTVIDKDCKLPMGLASGIIQDSQISASGHVGYWEPKLARLNNTGKYNAWSIIKKEHEHPWIQIDLQRQVVITGIQTQGAMQ
*Pseudonaja textilis F5* QALFTVIDKDCKLPMGLASGIIQDSQISASGHVGYWEPKLARLNNTGKYNAWSIIKKEHEHPWIQIDLQRQVVITGIQTQGAMQ

*Pseudonaja textilis vF5* LLQHSYTVEYFVTYSEDGQNWITFK**GRHSETQMHFEGNSDGTTVKENHIDPPIIAR**YIRLHPTK**FYNRPTFR**IELLGCEVEGCS
*Pseudonaja textilis F5* (short) LLKHLYTVEYFFTYSKDGQNWITFK**GRHSETQMHFEGNSDGTTVKENHIDPPIIAR**YIRLHPTKFHNRPTFRIELLGCEVEGCS
*Pseudonaja textilis F5* LLKHLYTVEYFFTYSKDGQNWITFK**GRHSETQMHFEGNSDGTTVKENHIDPPIIAR**YIRLHPTKFHNRPTFRIELLGCEVEGCS

*Pseudonaja textilis vF5* VPLGMESGAIK**NSEITASSYKK**TWWSSWEPFLARLNLEGGTNAWQPEVNNKDQWLQIDLQHLTKITSIITQGATSMTTSMYVKT
*Pseudonaja textilis F5* (short) VPLGMESGAIK**NSEITASSYKK**TWWSSWEPSLARLNLKGRTNAWQPKVNNKDQWLQIDLQHLTKITSIITQGATSMTTSMYVKT
*Pseudonaja textilis F5* VPLGMESGAIK**NSEITASSYKK**TWWSSWEPSLARLNLKGRTNAWQPKVNNKDQWLQIDLQHLTKITSIITQGATSMTTSMYVKT

*Pseudonaja textilis vF5* FSIHYTDDNSTWKPYLDVRTSMEK**VFTGNINSDGHVKHFFKPPILSR**FIRIIPKTWNQYIALRIELFGCEVF
*Pseudonaja textilis F5* (short) FSIHYTDDNSTWKPYLDVRTSMEK**VFTGNINSDGHVKHFFKPPILSR**FIRIIPKTWNQYIALRIELFGCEVF
*Pseudonaja textilis F5* FSIHYTDDNSTWKPYLDVRTSMEK**VFTGNINSDGHVKHFFKPPILSR**FIRIIPKTWNQYIALRIELFGCEVF

**Fig. S11. Coagulation factor V protein is detected in Eastern brown snake (*Pseudonaja textilis*) venom.** Amino acid sequence alignment of hypothetical F5 protein translations with tryptic peptides shown in bold. Red residues indicate peptides unique to the venom protein (vF5), purple residues indicate peptides unique to the hemostatic liver isoforms (F5 or F5 short), and black residues indicate peptides shared by both paralogs. Residues shown in grey correspond to the signal peptide. Tryptic peptides corresponding to the hemostatic form were detected in venoms from all individuals (N = 7).

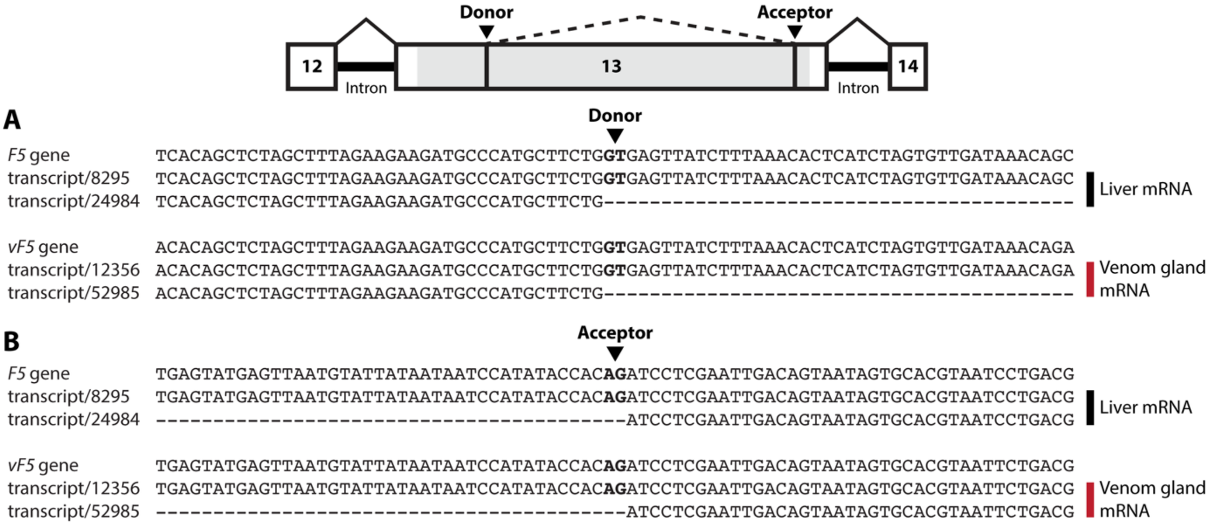

**Fig. S12. *F5* gene paralogs are alternatively spliced within exon 13.** DNA and mRNA sequence comparison at the donor (A) and acceptor (B) splice sites within exon 13 of both *F5* paralogs. The alignments include the *F5* and *vF5* genes as references with each of their corresponding full-length RNA transcripts expressed in liver or venom gland tissue of the inland taipan (*Oxyuranus microlepidotus*). The longer RNA isoform encodes the entire exon 13 (transcripts 8295 and 12356), whereas the shorter isoform (transcripts 24984 and 52985) retains only two short coding sequences at the 5′ and 3′ ends of exon 13. These segments are flanked by two canonical splice sites (donor GT and acceptor AG), indicating that the shorter isoform is generated through alternative splicing within exon 13.

| | 1 … 76 |
|---|---|
| *P. textilis* Acidic PLA2 1 | MHPAHLLVLLGVYVSLLGAARIPPLPLSLDDFSNLITCANRGSRSWLDYAHYGCFCGSGGSGT |
| *P. textilis* Acidic PLA2 2 | MHPAHLLVLLGVCVSLLGAARIPPLPLSLVQFRILIKCANHNSRNVLDYADYGCYCGKGGSGT |
| *O. microlepidotus* Acidic PLA2 1 OS1 | MHPAHLLVLLAVCVSLLGAARIPPLPLSLLDFANLIECANHGTRSALAYVHYGCYCGKGGRGT |
| *N. scutatus* Acidic PLA2 HTe | MYPAHLQVLLAVCVSLLGASSIPARPLNLYQFGNMIQCANHGRRPTRHYMDYGCYCGKGGSGT |
| *P. textilis* Acidic PLA2 textilotoxin D subunit | MHPAHLLVLLGVCVSRLGAASIPRPSLNIMLFGNMIQCTIPCEQSWLGYLDYGCYCGSGSSGI |
| *O. microlepidotus* Acidic PLA2 taipoxin γ subunit | MHPAHLLVLLAVCVSLLGSSEIPQPSLDFEQFSNMIQCTIPCGESCLAYMDYGCYCGPGGSGT |
| *B. candidus* Acidic PLA2 Bc-PL | MNPAHLLVLSALCVSLLGAANIPPQPLNLLQFNEMIECTIPGSFPLLDYMDYGCYCGTGGRGT |
| *B. candidus* Acidic PLA2 1 | MNPAHFLVLAAVCVSLLGAANIPPQPLSFYRYTQMIQCTIRGSLTYLDYMDYGCYCGTGSRGT |
| *B. candidus* Acidic β bungarotoxin A3 | MNPAHLLVLAAVCVSLLGAANIPPHPLNLINFMEMIRYTIPCEKTWGEYTNYGCYCGAGGSGR |
| *B. candidus* Basic β bungarotoxin A2 | MNPAHLLVLSAVCVSLLGAANIPPHPFNLINFMKMIRYTIPCEKTWGEYVDYGCYCGVGGSGR |
| *B. candidus* Basic β bungarotoxin A1 | MNPAHLLVLSAVCVSLLGAANIPPHPLNLINFMEMIRYTIPCEKTWGEYADYGCYCGAGGSGR |
| *N. scutatus* Basic PLA2 notechis 11'2 | MYPAHLLVLLAVCVSLLGAASIPPQPLNLYQFGNMIQCANHGSRATWHYMDYGCYCGAGGSGT |
| *N. scutatus* Basic PLA2 notexin | MYPAHLLVLLAVCVSLLGAASIPPQPLNLVQFSYLIQCANHGSRATWHYMDYGCYCGAGGSGT |
| *N. scutatus* PLA2 1 | MYPAHLLVLLAVCVSLLGATSIPPQPLNLYQFKNMIPCANHGRRPTWHYMDYGCYCSSGGSGT |
| *N. scutatus* Acidic PLA2 | MYPAHLLVLLAVCVSLLGAASIPPLPLNVAQFDNMIECANYGSRPSWHYMEYGCYCGKEGSGT |
| *P. textilis* Basic PLA2 textilotoxin A subunit | MHPAHLLVLLGVCVSLLGASDIPPLPLNLVQFSYLIRCANKYKRPGWHYANYGCYCGSGGRGT |
| *P. textilis* Basic PLA2 textilotoxin B subunit | MHPAHLLVLLAVYVSLLGAARIPPLPLDLVEFGFMIRCANRNSQPAWQYMDYGCYCGKRGSGT |
| *O. microlepidotus* Neutral PLA2 taipoxin β subunit | MHPAHLLVLLAVCVSLLGASDIPPLPLNLLQFGFMIECAIRNRQPALDFMNYGCYCGTVGRGT |
| *P. textilis* Acidic PLA2 textilotoxin C subunit | MHPAHLLVLLGVYVSLLGAARIPPLPLNLIQFSNMIKCTIPGSQPLLDYANYGCYCGPGNNGT |
| *O. microlepidotus* Basic PLA2 taipoxin α subunit 1 | MHPAHLLVLLAVCVSLLGASDIPPLPLNLLQFGFMIRCANRRSRPVWHYMDYGCYCGKGGRGT |
| *O. microlepidotus* Basic PLA2 taipoxin α subunit 2 | MHPAHLLVLLAVCVSLLGASDIPPLPLNLAQFGFMIKCANGGSRPVSHYMDYGCYCGKGGSGT |
| *O. microlepidotus* Basic PLA2 taipoxin α subunit 3 | MHPAHLLVLLAVCVSLLGASDIPPLPLNLAQFGFMIKCANGGSRPVSHYMDYGCYCGKGGSGT |
| *O. microlepidotus* Basic PLA2 taipoxin α subunit 4 | MHPAHLLVLLAVCVSLLGASDIPPLPLNLAQFGFMIKCANGGSRPVSHYMDYGCYCGKGGSGT |
| *O. microlepidotus* Basic PLA2 taipoxin α subunit 5 | MHPAHLLVLLAVCVSLLGASDIPPLPLNLAQFGFMIKCANGGSRPVSHYMDYGCYCGKGGSGT |
| *O. microlepidotus* Basic PLA2 taipoxin α subunit 6 | MHPAHLLVLLAVCVSLLGASDIPPLPLNLAQFGFMIKCANGGSRPVSHYMDYGCYCGKGGSGT |
| *P. textilis* Group IB PLA2 | MNPAHLLVLLAVCVSLLGASDIPPRPLNLIQFGNMIKCTIPGSRPLLDYADYGCYCGPGGSGT |
| *O. microlepidotus* Group IB PLA2 | MNPAHLLVLLAVCVSLLGASDIPPLPLNLIQFGNMIKCTIPGSRPLLDYADYGCYCGPGGSGT |
| *B. candidus* Group IB PLA2 | MNPAHLLVLSAVCVSLLGAANVAPQPRNLYQFRNMIKCTIPGSRPLLDYANYGCYCGKGGRGT |
| *N. helvetica* Group IB PLA2 | MNLAHLLVPAALCVSLVAASALPSLPRNLLQFSNMIKCAIPGSRPVLDYADYGCYCGFGGSGT |

| | 77 … 126 |
|---|---|
| *P. textilis* Acidic PLA2 1 | PVDDLDRCCQVHDNCFGDAEKLPACN**YLFSG**PYWNPYSYKCNEGE-ITCTDDNDECAAFICNC |
| *P. textilis* Acidic PLA2 2 | PVDELDRCCQAHDYCYDDAEKLPACN**YRFSG**PYWNPYSYKCNEGE-VTCTDDNDECKAFICNC |
| *O. microlepidotus* Acidic PLA2 1 OS1 | PVDDLDRCCQVHDDCYGEAEKLPACN**YLMSS**PYFNSYSYKCNEGK-VTCTDDNDECKAFICNC |
| *N. scutatus* Acidic PLA2 HTe | PVDELDRCCKTHDDCYGEAEKLPACN**YMMSG**PYYNTYSYECNEGE-LTCKDDNDECGAFICNC |
| *P. textilis* Acidic PLA2 textilotoxin D subunit | PVDDVDKCCKTHDECYYKAGQIPGCS**VQPNE**VFNVDYSYECNEGQ-LTCNESNNECEMAVCNC |
| *O. microlepidotus* Acidic PLA2 taipoxin γ subunit | PIDDLDRCCKTHDECYAEAGKLSACK**SVLSE**PNNDTYSYECNEGQ-LTCNDDNDECKAFICNC |
| *B. candidus* Acidic PLA2 Bc-PL | PVDMLDRCCKEHDDCYAQIKENPKCS**SLLNV**PYVKQYSFTCSEGD-LTCSDAAGTCARIVCDC |
| *B. candidus* Acidic PLA2 1 | PVDELDRCCQTHDNCYAEAEEHPKCS**SLVKS**PYMNLYSYTCSGGT-ITCNADNDECGAFICNC |
| *B. candidus* Acidic β bungarotoxin A3 | PIDALDRCCYVHDNCYGDAANIRDC-----**N**PKTQSYSYKLTKRT-IICYGAAGTCARVVCDC |
| *B. candidus* Basic β bungarotoxin A2 | PIDALDRCCYVHDNCYGDAEKKHKC-----**N**PKMQSYSYKLTKRT-IICYGAAGTCARIVCDC |
| *B. candidus* Basic β bungarotoxin A1 | PIDALDRCCYVHDNCYGDAEKKHKC-----**N**PKTQSYSYKLTKRT-IICYGAAGTCARIVCDC |
| *N. scutatus* Basic PLA2 notechis 11'2 | PVDELDRCCKAHDDCYTEAGKK-GC-----**S**PKLTLYSWKCIEKT-PTC-NSKTGCERSVCDC |
| *N. scutatus* Basic PLA2 notexin | PVDELDRCCKIHDDCYGDAEKK-GC-----**S**PKMSAYDYYCGENG-PYCRNIKKKCLRFVCDC |
| *N. scutatus* PLA2 1 | PVDELDRCCKTHDDCYTNIGTS-KC-----**N**PKSQIYSWKCGSDG-PTC-DSKTGCERSVCEC |
| *N. scutatus* Acidic PLA2 | PVDELDRCCKAHDDCYTEAEKR-RC-----**H**PKFSAYSWKCGSDG-PTC-DSETGCKRTVCDC |
| *P. textilis* Basic PLA2 textilotoxin A subunit | PVDDVDRCCQAHDKCYEDAEKL-GC-----**Y**PKWTTYYYYCGANG-PYC-KTRTKCQRFVCNC |
| *P. textilis* Basic PLA2 textilotoxin B subunit | PVDDVDRCCQTHNECYDEAAKIPGC-----**K**PKWTFYFWQCGSGSQFTCRKSKDVCRNVVCDC |
| *O. microlepidotus* Neutral PLA2 taipoxin β subunit | PVDDLDRCCKTRNECYAEAEKH-GC-----**Y**PSLTTYRWQCGRVG-LHC-NSKTQCEVFVCAC |
| *P. textilis* Acidic PLA2 textilotoxin C subunit | PVDDVDRCCQAHDECYDEASNH-GC-----**Y**PELTLYDYYCDTGV-PYC-KARTQCQVFVCGC |
| *O. microlepidotus* Basic PLA2 taipoxin α subunit 1 | PVDDLDRCCQVHDECYGEAERRFGC-----**S**PYWTLYSWKCYGTA-PSC-NTKTRCQRFVCRC |
| *O. microlepidotus* Basic PLA2 taipoxin α subunit 2 | PVDELDRCCQVHDECYGEAEKRFKC-----**V**PYMTLYSWKCYGTA-PSC-NTKTDCQRFVCNC |
| *O. microlepidotus* Basic PLA2 taipoxin α subunit 3 | PVDELDRCCQVHDECYGEAEKRFKC-----**V**PYMTLYSWKCYGTA-PSC-NTKTDCQRFVCNC |
| *O. microlepidotus* Basic PLA2 taipoxin α subunit 4 | PVDELDRCCQVHDECYGEAEKRFKC-----**V**PFVTLYSWKCYGKA-PSC-NTKTDCQRFVCNC |
| *O. microlepidotus* Basic PLA2 taipoxin α subunit 5 | PVDELDRCCQVHDECYGEAEKRFKC-----**V**PFVTLYSWKCYGKA-PSC-NTKTDCQRFVCNC |
| *O. microlepidotus* Basic PLA2 taipoxin α subunit 6 | PVDELDRCCQVHDECYGEAEKRFKC-----**V**PYMTLYSWKCYGTA-PSC-NTKTDCQRFVCNC |
| *P. textilis* Group IB PLA2 | PVDELDRCCQTHDNCYSQAKKHPACK**SLLDS**PYTKIYSYTCSGGN-LTCKDDNDECGAFICNC |
| *O. microlepidotus* Group IB PLA2 | PVDELDRCCQTHDNCYSQAKKHPACK**SLLDS**PYTKIYSYTCSGGN-LTCKDDNDECGAFICNC |
| *B. candidus* Group IB PLA2 | PVDMLDRCCQTHDNCYDQAMKHPACK**SFLDS**PYIKTYSYTCSGGT-VTCKDVKDKCARFVCDC |
| *N. helvetica* Group IB PLA2 | PVDQLDRCCQTHDNCYSQAKKHPACK**SLLDS**PYTKTYSYTCSGGN-ITCQDNNDECEAFVCNC |

| | 127 155 |
|---|---|
| *P. textilis* Acidic PLA2 1 | DRTAAICFAGATYNDENFMVTIKKKNICQ |
| *P. textilis* Acidic PLA2 2 | DRTAAICFAGAPYNDENFMITTKKKNICQ |
| *O. microlepidotus* Acidic PLA2 1 OS1 | DRTAAICFAGATYNDENFMISKKRNDICQ |
| *N. scutatus* Acidic PLA2 HTe | DRTAAICFARAPYNDANWDINLIAR--CQ |
| *P. textilis* Acidic PLA2 textilotoxin D subunit | DRAAAICFARFPYNKNYWSINTEIH--CR |
| *O. microlepidotus* Acidic PLA2 taipoxin γ subunit | DRTAVTCFAGAPYNDLNYNIGMIEH--CK |
| *B. candidus* Acidic PLA2 Bc-PL | DRTAALCFAEAPYKRRNFKIDYKTR--CQ |
| *B. candidus* Acidic PLA2 1 | DRTAALCFAKAPYNEENKEIDISKR--CQ |
| *B. candidus* Acidic β bungarotoxin A3 | DRTAALCFGDSEYIEGHKNIDTARF--CQ |
| *B. candidus* Basic β bungarotoxin A2 | DRTAALCFGNSEYIERHKNIDTKRY--CR |
| *B. candidus* Basic β bungarotoxin A1 | DRTAALCFGDSEYIERHKNIDTKRH--CQ |
| *N. scutatus* Basic PLA2 notechis 11'2 | DAAAAKCFAKAPYNKKNYNINTKKR--CQ |
| *N. scutatus* Basic PLA2 notexin | DVEAAFCFAKAPYNNANWNIDTKKR--CQ |
| *N. scutatus* PLA2 1 | DAAAAKCFAKAPYNDANWNIDTEKH--CQ |
| *N. scutatus* Acidic PLA2 | DATAAKCFAKAPFNQANWNIDTETH--CQ |
| *P. textilis* Basic PLA2 textilotoxin A subunit | DVVAADCFASYPYNRRYWFYSNKKR--CR |
| *P. textilis* Basic PLA2 textilotoxin B subunit | DFKAALCLTRARYNSANYNIDIKTH--CR |
| *O. microlepidotus* Neutral PLA2 taipoxin β subunit | DLAAAKCLAQEDYNPAHFNINTKAR--CR |
| *P. textilis* Acidic PLA2 textilotoxin C subunit | DLAVAKCLAGATYNDENKNINTGER--CQ |
| *O. microlepidotus* Basic PLA2 taipoxin α subunit 1 | DAKAAECFARSPYQSSNWNINTKAR--CR |
| *O. microlepidotus* Basic PLA2 taipoxin α subunit 2 | DAKAAECFARSPYQNKNWNINTKAR--CK |
| *O. microlepidotus* Basic PLA2 taipoxin α subunit 3 | DAKAAECFARSPYQNKNWNINTKAR--CK |
| *O. microlepidotus* Basic PLA2 taipoxin α subunit 4 | DAKAAECFARSPYQNKNWNINTKAR--CK |
| *O. microlepidotus* Basic PLA2 taipoxin α subunit 5 | DAKAAECFARSPYQNKNWNINTKAR--CK |
| *O. microlepidotus* Basic PLA2 taipoxin α subunit 6 | DAKAAECFARSPYQNKNWNINTKAR--CK |
| *P. textilis* Group IB PLA2 | DRTAAICFAGAPYNKENYNIDTTKH--CK |
| *O. microlepidotus* Group IB PLA2 | DRTAAICFAGAPYNKENYNIDTTKH--CK |
| *B. candidus* Group IB PLA2 | DRAAAMCFARAPYNKKYKKLDKSKY--CK |
| *N. helvetica* Group IB PLA2 | DRSAAICFAGAPYNEEYKKLDTGKY--CK |

**Fig. S13. The pancreatic loop is intact in group IB but not in group IA phospholipase $A_2$ venom proteins.** Amino acid sequence alignment of hypothetical protein translations of group I phospholipase $A_2$ ($PLA_2$) genes. Residues shown in grey encode the signal peptide and propeptide regions, and residues in bold indicate the pancreatic loop in group IB $PLA_2$, which is absent in group IA $PLA_2$ proteins because of a 5-amino-acid deletion. The procoagulant proteins are "*P. textilis* Acidic PLA2 1", "*P. textilis* Acidic PLA2 2" and putatively "*O. microlepidotus* Acidic PLA2 1 OS1" based on its orthologous relationship to "*P. textilis* Acidic PLA2 1".

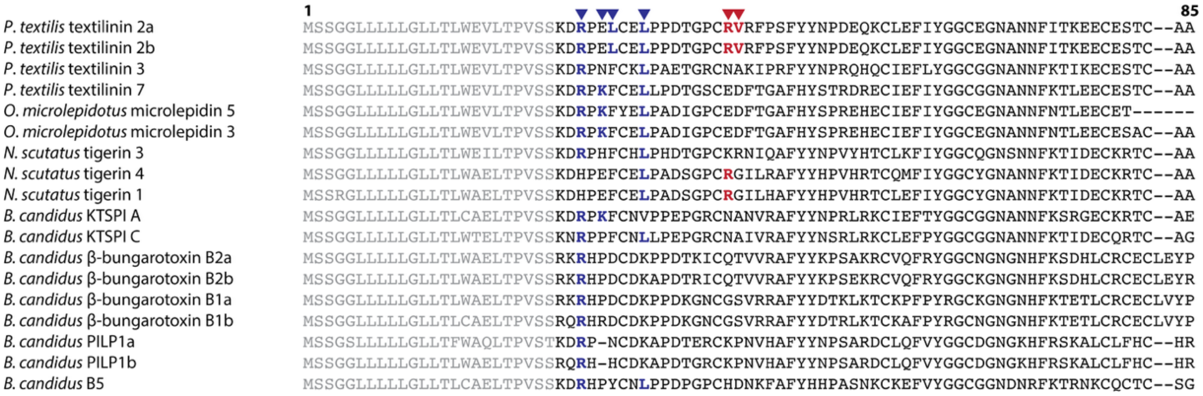

**Fig. S14. Australian elapid Kunitz-type toxins possess key residues associated with Kv1 potassium channel binding.** Amino acid sequence alignment of hypothetical protein translations of Kunitz-type toxin genes**.** Residues shown in grey encode the signal peptide, and residues associated with plasmin inhibition or specificity are highlighted in red (7), whereas residues associated with Kv1 potassium-channel binding are highlighted in blue (8). Note that the β-bungarotoxin paralogs are known potassium-channel inhibiting neurotoxins (9, 10).

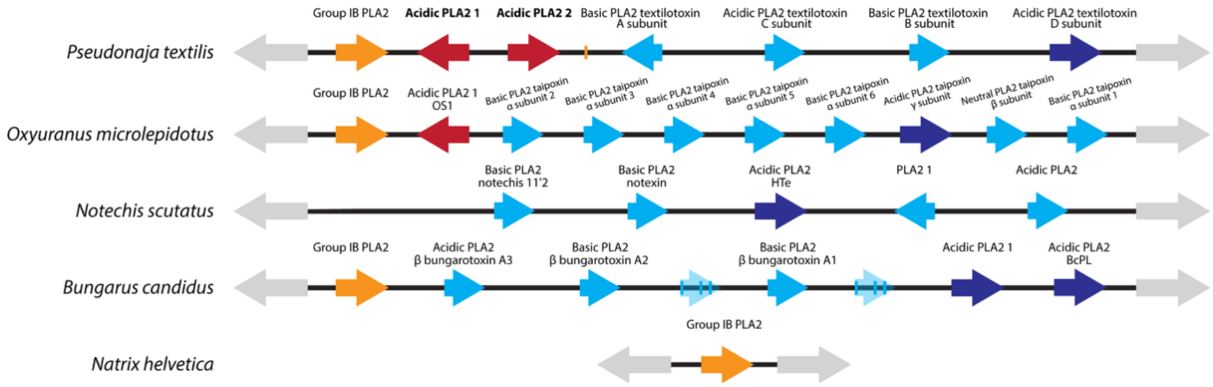

**Fig. S15. Venom-expressed neurotoxic PLA2GIB proteins were neofunctionalized into procoagulant toxins.** Protein identifications for the group I phospholipase genes shown in Fig. 5A, including procoagulant PLA2GIB proteins (red arrows), PLA2GIB proteins with neurotoxic or unknown activity (dark blue), neurotoxic PLA2GIA proteins (light blue), and the non-venom-expressed group IB PLA2 ortholog (orange). The *O. microlepidotus* Acidic PLA2 1 OS1 is a possible procoagulant protein based on its orthologous relationship with *P. textilis* Acidic PLA2 1; however, this protein has not been biochemically characterized.

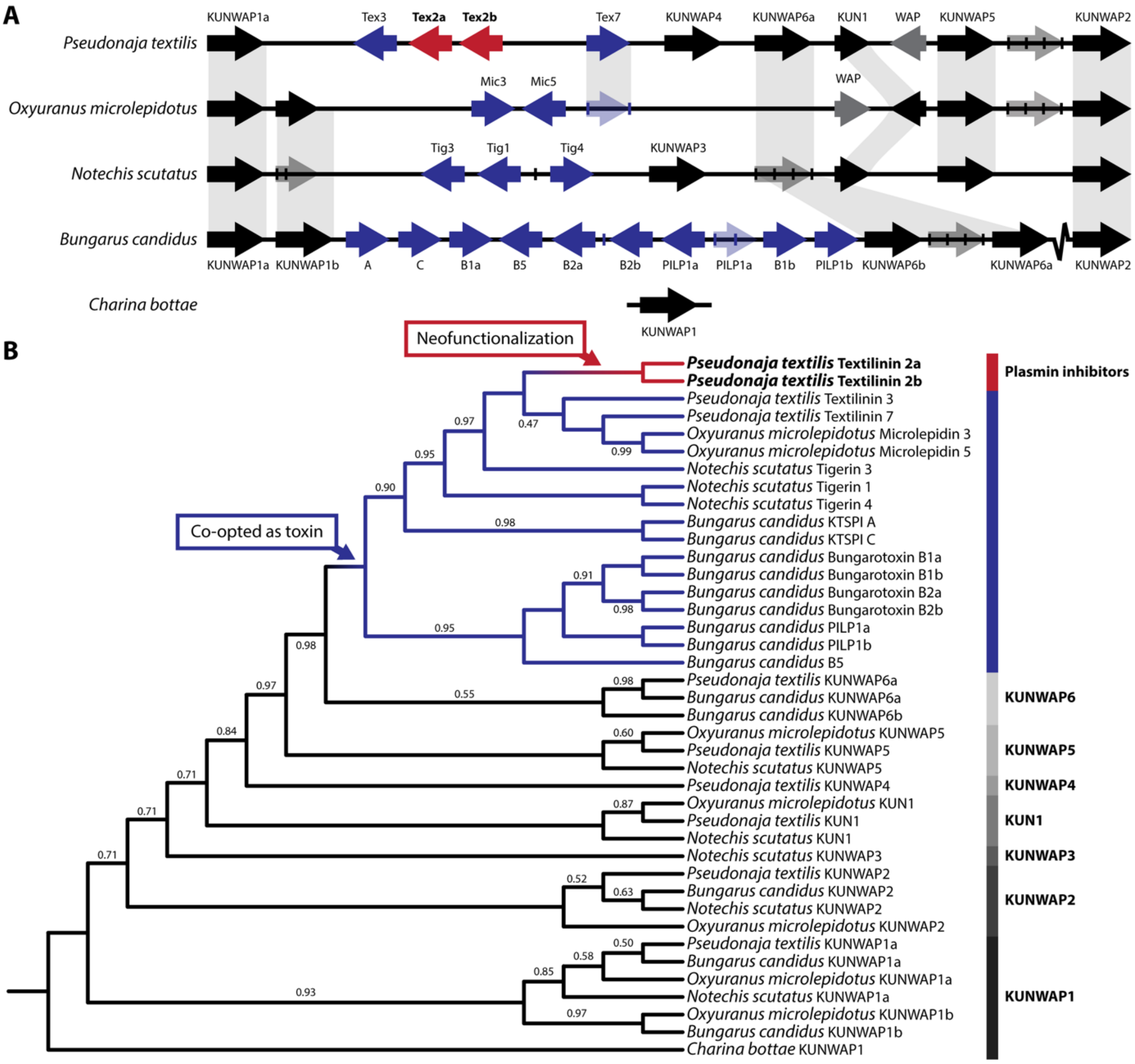

**Fig. S16. Venom-expressed Kunitz-type toxins were neofunctionalized into plasmin-inhibiting proteins.**

(A) A schematic showing the synteny of venom-expressed and non-venom-expressed Kunitz-type serine protease inhibitor genes across different elapid snakes. Pseudogenes are indicated by transparent arrows. The break-line symbol represents a genomic region with low confidence.

(B) A protein phylogeny of Kunitz-type serine protease inhibitor genes. Full phylogeny of Figure 6, with all Kunitz–Waprin protein (KUNWAP) sequences. The plasmin-inhibiting proteins (Textilinin 2a and Textilinin 2b) cluster within the clade of all venom-expressed genes. The phylogeny was rooted with the single-copy gene of *C. bottae* as the outgroup. Only Bayesian posterior probabilities less than 1.00 are shown.

**Tables**

**Table S1. Comparison of gene expression levels between venom gland and liver tissues in *Pseudonaja textilis* using short-read RNA-sequencing.**

| Gene ID | Protein ID | Gene complex | Venom gland (TPM) | Liver (TPM) |
|---|---|---|---|---|
| *NF2* | Merlin | PLA2 | 69,59 | 34,15 |
| *PLA2G1B* | Group IB PLA2 | PLA2 | 3,99 | 0 |
| *PLA-1* | Acidic PLA2 1 | PLA2 | 140262,47 | 120,94 |
| *PLA-2* | Acidic PLA2 2 | PLA2 | 97315,99 | 99,91 |
| *PLA-3* | Basic PLA2 textilotoxin A subunit | PLA2 | 23292,45 | 3 |
| *PLA-4* | Acidic PLA2 textilotoxin C subunit | PLA2 | 42512,56 | 6 |
| *PLA-5* | Basic PLA2 textilotoxin B subunit | PLA2 | 13785,71 | 2,87 |
| *PLA-6* | Acidic PLA2 textilotoxin D subunit | PLA2 | 27505,92 | 0 |
| *THOC5* | THO complex subunit 5 | PLA2 | 394,56 | 335,28 |
| *KUNWAP1a* | Kunitz–Waprin protein 1a | KUN | 12,16 | 314,3 |
| *Textilinin-3* | Textilinin 3 | KUN | 12137,17 | 0 |
| *Textilinin-2a* | Textilinin 2a | KUN | 274025,93 | 167,99 |
| *Textilinin-2b* | Textilinin 2b | KUN | 274025,93 | 167,99 |
| *Textilinin-7* | Textilinin 7 | KUN | 49246,97 | 48 |
| *KUNWAP4* | Kunitz–Waprin protein 4 | KUN | 23,94 | 0 |
| *KUNWAP6a* | Kunitz–Waprin protein 6a | KUN | 0 | 0 |
| *KUN1* | Kunitz protein 1 | KUN | 0 | 0 |
| *WAP1* | Waprin 1 (Textwaprin) | KUN | 0 | 0 |
| *KUNWAP5* | Kunitz–Waprin protein 5 | KUN | 0 | 0 |
| *KUNWAP2* | Kunitz–Waprin protein 2 | KUN | 0 | 0 |

Abbreviation: TPM, Transcripts per Million. Similar TPM values were obtained using Kallisto (data not shown) (11).

**Table S2. Comparison of gene expression levels between venom gland and liver tissues in the Inland taipan (*Oxyuranus microlepidotus*) using short-read RNA-sequencing.**

| Gene ID | Protein ID | Gene complex | Venom gland (TPM) | Liver (TPM) |
|---|---|---|---|---|
| *NF2* | Merlin | PLA2 | 6,94 | 152,88 |
| *PLA2G1B* | Group IB PLA2 | PLA2 | 0,47 | 0 |
| *PLA-1* | Acidic PLA2 1 OS1 | PLA2 | 22561,82 | 0 |
| *PLA-2* | Basic PLA2 taipoxin α subunit 2 | PLA2 | 176034,55 | 0,61 |
| *PLA-3* | Basic PLA2 taipoxin α subunit 3 | PLA2 | 176034,55 | 0,61 |
| *PLA-4* | Basic PLA2 taipoxin α subunit 4 | PLA2 | 166106,5 | 0,3 |
| *PLA-5* | Basic PLA2 taipoxin α subunit 5 | PLA2 | 166106,5 | 0,3 |
| *PLA-6* | Basic PLA2 taipoxin α subunit 6 | PLA2 | 96369,47 | 0 |
| *PLA-7* | Acidic PLA2 taipoxin γ subunit | PLA2 | 74955,84 | 0 |
| *PLA-8* | Neutral PLA2 taipoxin β subunit | PLA2 | 53611,41 | 0 |
| *PLA-9* | Basic PLA2 taipoxin α subunit 1 | PLA2 | 42857,19 | 0 |
| *THOC5* | THO complex subunit 5 | PLA2 | 31,85 | 1273,86 |
| *KUNWAP1a* | Kunitz–Waprin protein 1a | KUN | 0 | 0 |
| *KUNWAP1b* | Kunitz–Waprin protein 1b | KUN | 0 | 0 |
| *Microlepidin-3* | Microlepidin 3 | KUN | 880,78 | 0 |
| *Microlepidin-5* | Microlepidin 5 | KUN | 218,67 | 0 |
| *WAP1* | Waprin 1 (Omwaprin) | KUN | 20991,74 | 0 |
| *KUN1* | Kunitz protein 1 | KUN | 0,81 | 0 |
| *KUNWAP5* | Kunitz–Waprin protein 5 | KUN | 1,98 | 0 |
| *KUNWAP2* | Kunitz–Waprin protein 2 | KUN | 0 | 115,42 |

Abbreviation: TPM, Transcripts per Million. Similar TPM values were obtained using Kallisto (data not shown) (11).

**SI References**

1. A. M. Bolger, M. Lohse, B. Usadel, Trimmomatic: a flexible trimmer for Illumina sequence data. *Bioinformatics* **30**, 2114-2120 (2014).
2. H. Li, Aligning sequence reads, clone sequences and assembly contigs with BWA-MEM. *arXiv preprint arXiv:1303.3997* (2013).
3. P. Danecek *et al.*, Twelve years of SAMtools and BCFtools. *GigaScience* **10**, giab008 (2021).
4. J. T. Robinson *et al.*, Integrative genomics viewer. *Nature Biotechnology* **29**, 24-26 (2011).
5. H. Thorvaldsdóttir, J. T. Robinson, J. P. Mesirov, Integrative Genomics Viewer (IGV): high-performance genomics data visualization and exploration. *Briefings in Bioinformatics* **14**, 178-192 (2013).
6. M. A. Reza, T. N. Minh Le, S. Swarup, R. Manjunatha Kini, Molecular evolution caught in action: gene duplication and evolution of molecular isoforms of prothrombin activators in Pseudonaja textilis (brown snake). *Journal of Thrombosis and Haemostasis* **4**, 1346-1353 (2006).
7. E.-K. I. Millers *et al.*, The structure of Human Microplasmin in Complex with Textilinin-1, an Aprotinin-like Inhibitor from the Australian Brown Snake. *PLOS ONE* **8**, e54104 (2013).
8. S. Gasparini *et al.*, Delineation of the functional site of alpha-dendrotoxin: The functional topographies of dendrotoxins are different but share a conserved core with those of other Kv1 potassium channel-blocking toxins. *Journal of Biological Chemistry* **273**, 25393-25403 (1998).
9. C. G. Benishin, Potassium channel blockade by the B subunit of beta-bungarotoxin. *Molecular Pharmacology* **38**, 164-169 (1990).
10. P.-F. Wu, S.-N. Wu, C.-C. Chang, L.-S. Chang, Cloning and functional expression of B chains of β-bungarotoxins from Bungarus multicinctus (Taiwan banded krait). *Biochemical Journal* **334**, 87-92 (1998).
11. N. L. Bray, H. Pimentel, P. Melsted, L. Pachter, Near-optimal probabilistic RNA-seq quantification. *Nature Biotechnology* **34**, 525-527 (2016).
